# Yeast poly(A)-binding protein (Pab1) controls translation initiation in vivo primarily by blocking mRNA decapping and decay

**DOI:** 10.1101/2024.04.19.590253

**Authors:** Poonam Poonia, Vishalini Valabhoju, Tianwei Li, James Iben, Xiao Niu, Zhenguo Lin, Alan G. Hinnebusch

## Abstract

Poly(A)-binding protein (Pab1 in yeast) is involved in mRNA decay and translation initiation, but its molecular functions are incompletely understood. We found that auxin-induced degradation of Pab1 reduced bulk mRNA and polysome abundance in a manner suppressed by deleting the catalytic subunit of decapping enzyme (*dcp2*Δ), demonstrating that enhanced decapping/degradation is the major driver of reduced mRNA abundance and protein synthesis at limiting Pab1 levels. An increased median poly(A) tail length conferred by Pab1 depletion was also nullified by *dcp2*Δ, suggesting that mRNA isoforms with shorter tails are preferentially decapped/degraded at limiting Pab1. In contrast to findings on mammalian cells, the translational efficiencies (TEs) of many mRNAs were altered by Pab1 depletion; however, these changes were broadly diminished by *dcp2*Δ, suggesting that reduced mRNA abundance is a major driver of translational reprogramming at limiting Pab1. Thus, assembly of the closed-loop mRNP via PABP-eIF4G interaction appears to be dispensable for normal translation of most yeast mRNAs in vivo. Interestingly, histone mRNAs and proteins are preferentially diminished on Pab1 depletion dependent on Dcp2, accompanied by activation of internal cryptic promoters in the manner expected for reduced nucleosome occupancies, revealing a new layer of post-transcriptional control of histone gene expression.

## INTRODUCTION

There is considerable evidence that the poly(A) (pA) tail and poly(A) binding protein (PABP, Pab1 in yeast) stimulate translation initiation, but the molecular mechanisms involved and the biological settings where this control operates are not fully understood (1). One possible mechanism involves formation of a “closed-loop” mRNP with translation initiation factor eIF4E, bound to the m^7^G cap, and Pab1, bound to the pA tail, interacting with adjacent sites on eIF4G, the scaffold subunit of eIF4F. Supporting this model, Pab1 mediates the stimulatory effect of pA tails on mRNA translation in yeast extracts at the step of 43S translation preinitiation complex (PIC) binding to mRNA (2) dependent on eIF4G-PABP interaction (3). Moreover, depleting mammalian PABP or disrupting PABP-eIF4G interaction was shown to weaken eIF4F binding and inhibit translation of polyadenylated reporter mRNAs in mammalian cell extracts (4) in a manner involving increased competition by RNA binding proteins with eIF4G for mRNA (5). Stable eIF4F binding should promote its activities in stimulating helicases eIF4A and Ded1 and recruiting the 43S translation preinitiation complex (PIC) through interactions with other initiation factors bound to the PIC (6,7). PABP can also interact with peptide release factor eRF3, via the C-terminal MLLE domain and eRF3 NTD (8) and might thereby enhance reinitiation by bridging post-termination ribosomes at the stop codon and eIF4F at the cap (9).

At odds with an important role in translation, depletion of PABP from mammalian cells broadly reduced levels of mRNAs and comparably diminished bulk protein synthesis, but had little effect on translational efficiencies (TEs) of individual mRNAs regardless of pA tail lengths (10,11). pA tail lengthening confers increased translation of maternally deposited mRNAs in early embryonic development (1); however, TE is not strongly correlated with pA tail lengths in post-embryonic animal cells (12–14). This uncoupling of TEs from pA tail lengths was attributed partly to the greater abundance of PABP and lack of competition for the factor in somatic cells. Moreover, depleting PABP in somatic cells appeared to confer preferential degradation of mRNAs with short pA tails whose TEs should be preferentially diminished at low PABP levels, precluding detection of their TE reductions on PABP depletion, whereas such mRNAs are apparently more stable in oocytes (10).

WT yeast cells do not exhibit the coupling between TE and pA tail length observed in oocytes (12), suggesting a limited role for yeast Pab1 in translational control. Consistent with this, mutations in yeast eIF4G that specifically disrupt eIF4G-Pab1 interaction did not reduce yeast cell growth unless combined with other eIF4G mutations that impair eIF4G-eIF4E interaction and RNA-binding by the eIF4G NTD (3,15). These last findings suggested that closed loop formation via eIF4G-Pab1 interaction is dispensable for yeast viability and represents only one of several interactions that stabilize eIF4F binding to mRNA. Moreover, RIP-seq analysis indicated that only a fraction of yeast mRNAs exhibits high occupancies of eIF4E, eIF4G, and Pab1, suggesting that closed-loop formation varies greatly among different yeast mRNAs in vivo (16). While not essential for viability and variable in occurrence, closed-loop formation could still influence the wide range of TEs observed among cellular mRNAs in WT yeast cells. Consistent with this, mRNAs exhibiting relatively high occupancies of eIF4E, eIF4G, and Pab1 and low occupancies of inhibitory eIF4E binding proteins (4E-BPs) Caf20 and Eap1, expected to have the greatest potential for closed-loop formation, are among the most highly translated mRNAs in yeast (16). The translation of mRNAs of lowest Pab1 and eIF4E/eIF4G occupancies could be disproportionately impaired by Pab1 depletion by eliminating their already compromised ability to form the closed-loop and achieve a high initiation rate.

PABP is also heavily involved in mRNA turnover but can exert opposing influences on this process. Pab1 can recruit or activate the deadenylases Pan2-Pan3 and the Ccr4 subunit of the Ccr4-Not complex, which cooperate in shortening the pA tail, thought to be a prerequisite for degradation of the transcript (17,18). Conversely, Pab1 impedes shortening of the pA tail to an oligo(A) segment that would be too short to bind Pab1 but long enough to recruit the Lsm1-Lsm7/Pat1 complex to promote mRNA decapping and decay (1). Studies using reporter mRNAs suggested that eliminating Pab1 from yeast decreases the rate of pA tail shortening and allows decapping and 5’-3’ degradation by Xrn1 without prior deadenylation. These findings, and the fact that deleting *XRN1* suppresses the lethality of deleting *PAB1,* suggested that, despite the ability to accelerate deadenylation, Pab1 impedes decapping and degradation as an important component of its essential function in yeast (19). The genome-wide consequences of eliminating Pab1 on steady-state mRNA levels have not been determined for yeast; however, as mentioned above, depleting PABP from mammalian cells reduced global mRNA abundance (10,11). If removing Pab1 from yeast similarly confers widespread reductions in mRNA levels, this could indirectly alter the relative TEs of many mRNAs by changing the ratios of mRNAs to 43S PICs or RNA binding proteins (RBPs) that influence activation of mRNAs by eIF4F and Pab1.

To address the role of Pab1 in regulating translation in yeast cells, we examined the genome-wide consequences of depleting Pab1 on steady-state mRNA abundance, pA tail length, TEs, and steady-state protein abundance, both in the presence or absence of the decapping enzyme Dcp1:Dcp2. Our results indicate that Pab1 broadly influences translation in yeast and that most of the TE changes observed for individual transcripts at limiting Pab1 appear to be indirect consequences of widespread accelerated mRNA decapping and decay. These results support a limited role for closed-loop assembly via Pab1-eIF4G association in dictating the translation rates of most mRNAs in yeast cells under the conditions of our experiments.

## MATERIALS AND METHODS

### Yeast strains and plasmids

All yeast strains employed are listed in Table S1. All plasmids employed in this work are listed in Table S2, and all primers utilized in plasmid or strain constructions are given in Table S3. The *pab1-AID* strains were constructed in two steps. First, plasmid DNA of pHQ2122, an integrative plasmid containing the *OsTIR* gene encoding *Oryza sativa* F-box protein Tir1 expressed from the yeast *ADH1* promoter, was digested with NruI to direct integration to the *LYS2* gene in WT strain W303 and *dcp2*Δ mutant CFY1016 to obtain Ura^+^ transformants. Growth on 5-fluoro-orotic acid medium generated the Ura^-^ segregants PMY1 and PMY3, respectively, and colony PCR-amplification of chromosomal DNA confirmed loss of the plasmid sequences and replacement of *LYS2* with the *lys2::P_ADH1_-OsTIR-7myc* allele in both strains. In the second step, the *AID*-9myc-hphNT1* cassette on p6023 was PCR-amplified using primers encoding 60bp 5’ or 3’ of the *PAB1* stop codon and used to transform PMY1 and PMY3 to hygromycin B resistance, generating strains PMY2 and PRY1, respectively. Colony PCR-amplification of chromosomal DNA confirmed C-terminal tagging of *PAB1* with the *AID*-9myc* degron.

Plasmid pIS385-*OSTIR1* (a gift from Carl Wu) is an integrative *URA3* plasmid derived from the *LYS2* disintegrator plasmid pIS385 (20) containing the coding sequences for the *Oryza sativa TIR* gene under control of the yeast *ADH1* promoter. Plasmid pHQ2122 was generated by inserting between the BlpI-EagI sites of pIS385-*OSTIR1* a BlpI-PacI fragment encoding 383bp of 3’ coding sequence of *OsTIR1*, PCR-amplified from pIS385-*OSTIR1* using primers 2157/2158, and a PacI-EagI fragment encoding 7 tandem Myc epitopes and the *ADH1* transcription terminator (T*_ADH1_*) amplified from pHQ1435 using primers 688/2159. pHQ1435 was constructed by inserting a HincII fragment containing *SPT4-7myc-HIS* that was PCR-amplifed from genomic DNA of strain HQY974 (21) at the HincII site of pBS(+SK) (22).

### **C**ell spotting growth assays

Yeast strains and their respective transformants harboring plasmids containing *DCP2* or empty vector were grown to saturation in liquid synthetic complete medium (SC) without uracil, diluted to OD_600_ of 0.1 and 10-fold serial dilutions were spotted on YPD agar medium containing 1mM of 1-Napthaleneacetic acid (NAA) or 1mM of KOH and incubated at 30°C.

### Polysome profiling

Polysome profiling was conducted as described previously (23) with the following modifications. Strain PMY1 was pre-cultured in liquid YPD medium at 30°C for 3h and thereafter NAA or KOH was added to a final concentration of 1mM and growth was continued for 6h before harvesting. Strains PMY2, PMY3 and PRY1 were cultured in YPD at 30°C for one division and grown for 6h after NAA addition to 1mM. For strain PMY2 without NAA treatment, cells were cultured in YPD at 30°C for 4h, KOH was added to 1mM, and growth continued for 6h. All cultures reached mid-logarithmic phase growth (OD_600_ of ∼0.6-0.7) at the time of harvesting. Cells were harvested by centrifugation and whole cell extracts (WCEs) were prepared by vortexing the cell pellet with two volumes of glass beads in ice cold 1X breaking buffer (20 mM Tris-HCl (pH 7.5), 50 mM KCl, 10 mM MgCl2, 1 mM DTT, 200 μg/mL heparin, 50 μg/mL cycloheximide (CHX), and 1 Complete EDTA-free Protease Inhibitor cocktail Tablet (Roche)/50 mL buffer). Thirty OD_260_ units of cleared lysates were loaded onto 15-45% (w/w) sucrose gradients and centrifuged at 39,000 rpm for 2.5 h. Gradients were fractionated using the BioComp Gradient Station. Polysome to monosome (P/M) ratios were calculated using Fiji software.

### Polysome profiling to measure ribosome content

For measuring total ribosome content, cells were cultured as above for polysome profiling and harvested and quick-chilled by pouring into centrifuge tubes filled with ice and centrifuged for 10 min at 7000 × g. Cell pellets were resuspended in an equal volume of Buffer A (20 mM Tris–HCl [pH 7.5], 50 mM NaCl, 1 mM DTT, 200 μM PMSF and 1 Complete EDTA-free Protease Inhibitor cocktail Tablet (Roche)/50 mL buffer) and WCEs prepared by vortexing with glass beads in the cold, followed by two cycles of centrifugation for 10 min at 3000 rpm and 15,000 rpm at 4°C, respectively. (Cycloheximide and MgCl2 were omitted from the lysis buffer in order to separate 80S ribosomes into 40S and 60S subunits.) Equal volumes of cleared lysate were resolved on 5–47% (w/w) sucrose gradients by centrifugation at 39,000 rpm for 3 hr at 4°C in a Beckman SW41Ti rotor. Gradient fractions were scanned at 260 nm using a gradient fractionator (Bio-comp Instruments, Triax), and the area under the 40S and 60S peaks quantified using Fiji software. To estimate the ribosomal content per cell volume, the combined areas under the 40S and 60S peaks were normalized by the OD_600_ values of the starting cultures.

### Ribosome profiling (Ribo-Seq) and RNA-Seq

Ribo-Seq and RNA-seq were performed in parallel as described previously (24) employing the modified protocol based on McGlincy and Ingolia (25).

#### (i) Ribosome-protected fragment (RPF) library preparation and sequencing

Strains were cultured as above for polysome profiling, harvested by fast-filtration and frozen in liquid nitrogen. Cells were lysed in a freezer mill in the presence of lysis buffer (20 mM Tris (pH 8), 140 mM KCl, 1.5 mM MgCl_2_, 1%Triton X-100, 500 μg/mL CHX. Cell lysates were transferred to a 50 mL conical tube, thawed, and spun at 3000 g for 5 min at 4°C. The supernatant was clarified by centrifugation at full speed for 10 min in a refrigerated benchtop centrifuge at 4°C. The clarified supernatant was divided into aliquots before being snap-frozen in liquid N_2_ and stored at −80°C.

Cell lysates were digested with RNase I (Ambion; AM2294) at 15U per A_260_ unit for 1 hr at room temperature (25°C) on a Thermomixer at 700 rpm and resolved by sedimentation through a 10–50% sucrose gradient to isolate the 80S monosome fraction, which was snap-frozen in liquid N_2_ and stored at −80°C. RNA was extracted from the 80S fractions using Trizol LS reagent (Thermo Fisher, #10296010) and Zymo Research RNA clean and concentrator kit (Zymo R1018). The 25-34 nt ribosome footprints were purified by electrophoresis in a 15% TBE-Urea gel. RPF library preparation was performed as described previously (24) employing a modification of the protocol described by McGlincy and Ingolia (25). Quality of the libraries was assessed with a Bioanalyzer 2100 (Agilent Technologies) using the High Sensitivity DNA Kit (Agilent 5067–4626) and quantified by Qubit. Sequencing was done on an Illumina NovaSeq6000 platform (single-end 100 bp reads) at the NHLBI DNA Sequencing and Genomics Core at NIH (Bethesda, MD).

#### (ii) RNA-seq library preparation and sequencing

Total RNA was extracted and purified from aliquots of the same snap-frozen cells described above using Trizol LS reagent (Thermo Fisher, #10296010) and Qiagen miRNeasy kit (Qiagen 217004). For spike-in normalization, ERCC RNA Spike-In Mix 1 (Thermo Fisher, #4456740) was added in equal amounts (2.4 μl of 1:100-fold dilution) to 1.2 μg of total RNA of each sample before library preparation. RNA sequencing libraries were constructed by the NHLBI DNA sequencing Core at NIH (Bethesda, MD) using the Illumina TruSeq Stranded mRNA Library Prep Kit and sequenced using the Illumina NovaSeq6000 (single-end 100 bp reads) platform. Prior to library preparation, rRNA was depleted using the Qiagen FastSelect yeast rRNA depletion kit.

#### (ii) Data analysis

For spike-in normalized RNA-seq data, fastq files were trimmed using CUTADAPT (DOI: https://doi.org/10.14806/ej.17.1.200), the trimmed reads were aligned to a modification of the SacCer3 assembly version of the *S. cerevisiae* genome sequence containing the ERCC sequences using HISAT2 (26). The total number of reads obtained from ERCC alignment was used to calculate the size factor for each sample and reads corresponding to yeast genes were normalized by the size factors. Differential expression analysis between different strains or conditions was conducted using DESeq2 (27) by setting the size factor to unity.

For Ribo-Seq data analysis, RPF library sequencing Fastq files (de-barcoded at the NHLBI core facility according to their 6-nt Illumina barcodes) were trimmed of their linkers and separated according to their 5-nt internal sample barcode using CUTADAPT. Contaminating tRNA and rRNA were removed with a BOWTIE2 alignment (28) to the index of noncoding RNAs. The remaining reads were aligned to the SacCer3 genome sequence using HISAT2. Read counts were generated using a custom R-script (https://github.com/hzhanghenry/RiboProR). Similarly, RNA sequencing fastq files were trimmed using CUTADAPT. Contaminating tRNA and rRNA were removed with a BOWTIE2 alignment to the index of noncoding RNAs, and the remaining reads aligned to the SacCer3 genome sequence using HISAT2. Read counts were generated using the custom R-script described above. Statistical analysis of changes in mRNA, RPFs, or TE values between biological replicates from different genotypes or culture conditions was conducted using DESeq2.

### Single-molecule polyadenylated tail (SM-PAT) sequencing

#### (i) SM-PAT-seq library preparation and sequencing

SM-PAT sequencing was conducted on the same total RNA samples described above that were subjected to RNA-Seq and Ribo-Seq. RNA samples were quantified by using a Nanodrop ND-1000 spectrophotometer (Thermo Fisher) and evaluated for quality using the Bioanalyzer 2100 (Agilent Technologies). SM-PAT libraries were constructed from 5µg of total RNA for each sample as previously described (30) by the NICHD Molecular Genomics Core at NIH (Bethesda, MD). Alignment of mRNAs for tail length assignments was performed against cDNA sequences obtained from Ensembl *Saccharomyces_cerevisiae*.R64-1-1 release 109.

### CAGE sequencing and data analysis

#### (i) CAGE library preparation and sequencing

CAGE sequencing was conducted on the same total RNA samples subjected to RNA-Seq and Ribo-Seq described above for *PAB1* and *pab1-AID* strains PMY1 and PMY2 treated with auxin. For *spt6-1004* mutant FY2180 and isogenic WT strain FY2181, cells were cultured in SC medium at 30°C and shifted to 39°C for 90 min before harvesting. For *set1Δset2*Δ mutant H4293 and isogenic WT strain BY4741, cells were cultured in SC medium at 30°C. Total RNA was prepared by hot-phenol extraction (31) from two biological replicates of each strain. RNA samples were quantified using a Nanodrop ND-1000 spectrophotometer (Thermo Fisher) and evaluated for quality using the Bioanalyzer 2100. CAGE libraries were constructed from 5µg of total RNA for each sample using the no-amplification non-tagging CAGE (nAnT-iCAGE) protocol (32) by K.K. DNAFORM of Japan. Briefly, first strand cDNAs were synthesized by reverse transcription reaction with random primers to generate RNA/cDNA hybrids. The 5’ cap structures on RNA strands were biotinylated and enriched by streptavidin beads. Linkers containing a library-specific barcode were ligated to the cDNAs after removal of RNA strands and release from the streptavidin beads. After completion of the 2nd strand synthesis, each CAGE library was sequenced using Illumina NextSeq500 (single-end, 75-bp reads).

#### (ii) CAGE data processing, alignment, rRNA filtering and identification of TSSs and TCs

The sequenced CAGE reads of each sample were aligned to the *S. cerevisiae* reference genome (sacCer3) using HISAT2 (26) with “--no-softclip” option for disabling the soft clipping to reduce false-positive transcription start sites (TSSs). CAGE reads mapped to rRNA genes were identified using rRNAdust (http://fantom.gsc.riken.jp/5/sstar/Protocols:rRNAdust) and were excluded from subsequent TSS analyses. TSS identification, inference of TSS clusters (TC, representing putative core promoters), and assigning TCs to their downstream genes were carried out by using TSSr (33). CAGE reads with a mapping quality score (MAPQ) > 20 were considered uniquely mapped reads, which were used for subsequent analyses. TSS read counts of biological replicates were then merged as a single sample. The transcription abundance of each TSS was quantified as the numbers of CAGE tags/reads supporting the TSS per million mapped reads (TPM). Only TSSs with TPM ≥ 0.1 were used to infer TCs using the “peakclu” method (34), with the following options "peakDistance=50, extensionDistance=25, localThreshold = 0.01". A set of consensus TCs of all samples was generated by using the “consensusCluster” function in TSSr with an option of “dis = 100”. Consensus TCs were then assigned to their downstream genes if they are within 1000 bp upstream and 50 bp downstream of the start codon of annotated ORFs (File S7). The TPM value of a consensus TC in a sample is the sum of TPM values of all TSSs within its range. The TPM value of a gene was calculated as the sum of all consensus TCs assigned to the gene. Differential expression of capped mRNAs was assessed by DESeq2 based on raw read counts of genes from CAGE sequencing.

### TMT-MS of global protein abundance

Three replicate cultures of PMY1, PMY2, PMY3 and PRY1 strains were cultured as above for polysome profiling. Cells were harvested by centrifugation for 5 min at 3000 x *g,* resuspended in nuclease-free water, collected by centrifugation, and stored at -80°C. WCEs were prepared in freshly prepared extraction buffer (8M urea, 25 mM triethylammonium-bicarbonate (TEAB; Thermo Fisher, 90114)) by washing the cell pellets once and resuspending again in extraction buffer, then vortexing with glass beads in the cold room. Lysates were clarified by centrifugation at 13,000 x *g* for 30 min and the quality of extracted proteins was assessed following SDS-PAGE using GelCode™ Blue Stain (Thermo Fisher, #24592) and quantified using the Bradford reagent. Lysates were stored at -80°C. Sample preparation and TMT-MS/MS (35) was performed at the IDeA National Resource for Quantitative Proteomics. Briefly, total protein from each sample was reduced, alkylated, and digested using filter-aided sample preparation (36) with sequencing-grade modified porcine trypsin (Promega). Tryptic peptides were labeled using tandem mass tag isobaric labeling reagents (Thermo Fisher) following the manufacturer’s instructions and combined into one 16-plex TMT pro sample group. The labeled peptide multiplex was separated into 46 fractions on a 100 x 1.0 mm Acquity BEH C18 column (Waters) using an UltiMate 3000 UHPLC system (Thermo Fisher) with a 50 min gradient from 99:1 to 60:40 buffer A:B ratio under basic pH conditions, and then consolidated into 24 super-fractions. Each super-fraction was then further separated by reverse phase XSelect CSH C18 2.5 um resin (Waters) on an in-line 150 x 0.075 mm column using an UltiMate 3000 RSLCnano system (Thermo Fisher). Peptides were eluted using a 75 min gradient from 98:2 to 60:40 buffer A:B ratio (Buffer A = 0.1% formic acid, 0.5% acetonitrile and Buffer B = 0.1% formic acid, 99.9% acetonitrile). Eluted peptides were ionized by electrospray (2.4 kV) followed by mass spectrometric analysis on an Orbitrap Eclipse Tribrid mass spectrometer (Thermo Fisher) using multi-notch MS3 parameters. MS data were acquired using the FTMS analyzer in top-speed profile mode at a resolution of 120,000 over a range of 375 to 1500 m/z. Following CID activation with normalized collision energy of 35.0, MS/MS data were acquired using the ion trap analyzer in centroid mode and normal mass range. Using synchronous precursor selection, up to 10 MS/MS precursors were selected for HCD activation with normalized collision energy of 65.0, followed by acquisition of MS3 reporter ion data using the FTMS analyzer in profile mode at a resolution of 50,000 over a range of 100-500 m/z.

Proteins were identified and MS3 reporter ions quantified using MaxQuant (Version 1.6.12.0, Max Planck Institute) against the UniprotKB Saccharomyces cerevisiae database (UP000002311, 01/2022) with a parent ion tolerance of 3 ppm, a fragment ion tolerance of 0.5 Da, and a reporter ion tolerance of 0.003 Da. Scaffold Q+S software (Proteome Software) was used to verify MS/MS based peptide and protein identifications (protein identifications were accepted if they could be established with less than 1.0% false discovery and contained at least 2 identified peptides; protein probabilities were assigned by the Protein Prophet algorithm (37) and to perform reporter ion-based statistical analysis. Protein TMT MS3 reporter ion intensity values are assessed for quality and normalized using ProteiNorm (DOI: 10.1021/acsomega.0c02564). The data were normalized using VSN (38) and statistical analysis was performed using Linear Models for Microarray Data (limma) with empirical Bayes (eBayes) smoothing to the standard errors (39). Proteins with an FDR adjusted p-value < 0.05 and fold-change > 2 were considered to differ significantly between two conditions under comparison.

### Western blot analysis

WCEs were prepared by trichloroacetic acid (TCA) extraction as previously described (40) and immunoblot analysis was conducted as described previously (41). After electroblotting to PVDF membranes (Millipore IPFL00010), membranes were probed with antibodies against myc (Roche 11-667-203-001), Pab1 (a kind gift from Maurice Swanson of University of Florida) and Hcr1 (42). Secondary antibodies employed were HRP-conjugated anti-rabbit (Cytiva, NA9340V) and anti-mouse IgG (Cytiva, NA931V). Detection was performed using enhanced chemiluminescence (ECL) Western Blotting Detection Reagent (Cytiva, RPN2016) and the Azure 200 gel imaging biosystem or Amersham Hyperfilm MP x-ray films (28906845).

### Data visualization and statistical analysis

Notched boxplots were constructed using a web-based tool at http://shiny.chemgrid.org/boxplotr/.

In all such plots, the upper and lower boxes contain the 2^nd^ and 3^rd^ quartiles, and the band gives the median. If the notches in two plots do not overlap, there is roughly 95% confidence that their medians are different. Correlation matrices displaying Spearman correlations between sequencing read counts from biological replicates were created using the corr.test and corrplot function in R and density scatter plots were created using the ggplot2 function in R. Volcano plots were generated using a web-based tool (https://huygens.science.uva.nl/VolcaNoseR/). Venn diagrams were generated using a web-based tool (https://www.biovenn.nl/) and the significance of gene set overlaps in Venn diagrams was evaluated with the hypergeometric distribution using a web-based tool (https://systems.crump.ucla.edu/hypergeometric/index.php). Hierarchical clustering analysis was conducted with the R heatmap.2 function from the R ‘gplots’ library, using the default hclust hierarchical clustering algorithm. Gene ontology (GO) analysis was conducted using the web-based tool at http://funspec.med.utoronto.ca/. Gene browser images were generated using the Integrative Genomics

Viewer (IGV 2.4.14, http://software.broadinstitute.org/software/igv/) (43).

## RESULTS

### Depletion of Pab1 leads to widespread reductions in mRNA abundance

We depleted Pab1 from yeast cells using an auxin-inducible degron mutant, *pab1-AID* (Fig. S1A). A 6 hr auxin treatment reduced Pab1-AID to low levels and impaired cell growth (Fig. S1B-C), in agreement with previous findings that Pab1 is essential in yeast (44). To examine the impact of Pab1-AID depletion on mRNA abundance, we conducted RNA-seq on *pab1-AID* and isogenic WT *PAB1* cultures treated or untreated with auxin. The results were highly reproducible among all three biological replicates obtained for each strain/condition (Fig. S1G), allowing us to combine the data from replicates for all downstream analyses. By normalizing RNA-Seq reads for the recovery of ERCC spike-in transcripts, we determined that depletion of Pab1-AID reduced bulk mRNA abundance by a factor of ∼0.59, assessed either by comparing *pab1-AID* to WT cells both treated with auxin, or *pab1-AID* cells with and without auxin treatment (Fig. 1A, cols. 2-3), suggesting a critical role for Pab1 in stabilizing most mRNAs. (In this and all subsequent notched box plots, when notches of boxes in different columns do not overlap, their medians differ at ∼95% confidence level.) Indeed, the majority of individual mRNAs across the transcriptome showed reduced abundance on Pab1-AID depletion under these same two comparisons between normal and depleted Pab1 levels (Fig. 1B, cols. 1-2, red hues). Furthermore, DESeq2 analysis comparing *pab1-AID* to WT cells both treated with auxin revealed significant reductions in abundance of 3294 mRNAs, but increased levels of only 424 mRNAs on Pab1-AID depletion (Fig. 1C). DEseq2 analysis comparing *pab1-AID* cells with and without auxin treatment revealed similar numbers of mRNAs, 3195 and 366, displaying significant reductions or increases in abundance, respectively (Fig. S1D). As the differentially expressed mRNAs identified in the two comparisons showed highly significant overlaps (Fig. S1E), for all subsequent analyses we examined the mRNAs at the intersections of the two sets in Fig. S1E. These two core groups, designated mRNA_dn_*pab1* and mRNA_up*_pab1*, exhibit similar median changes in mRNA abundance of 0.38-fold and 2.97 or 2.69-fold, respectively in comparing either *pab1-AID* to WT cells both treated with auxin (Fig. 1D, cols. 4 & 6) or *pab1-AID* cells with and without auxin treatment (Fig. S1F, cols. 4 & 6). Thus, the mRNAs up- or down-regulated on Pab1 depletion are well-defined by these results.

**Fig. 1.**
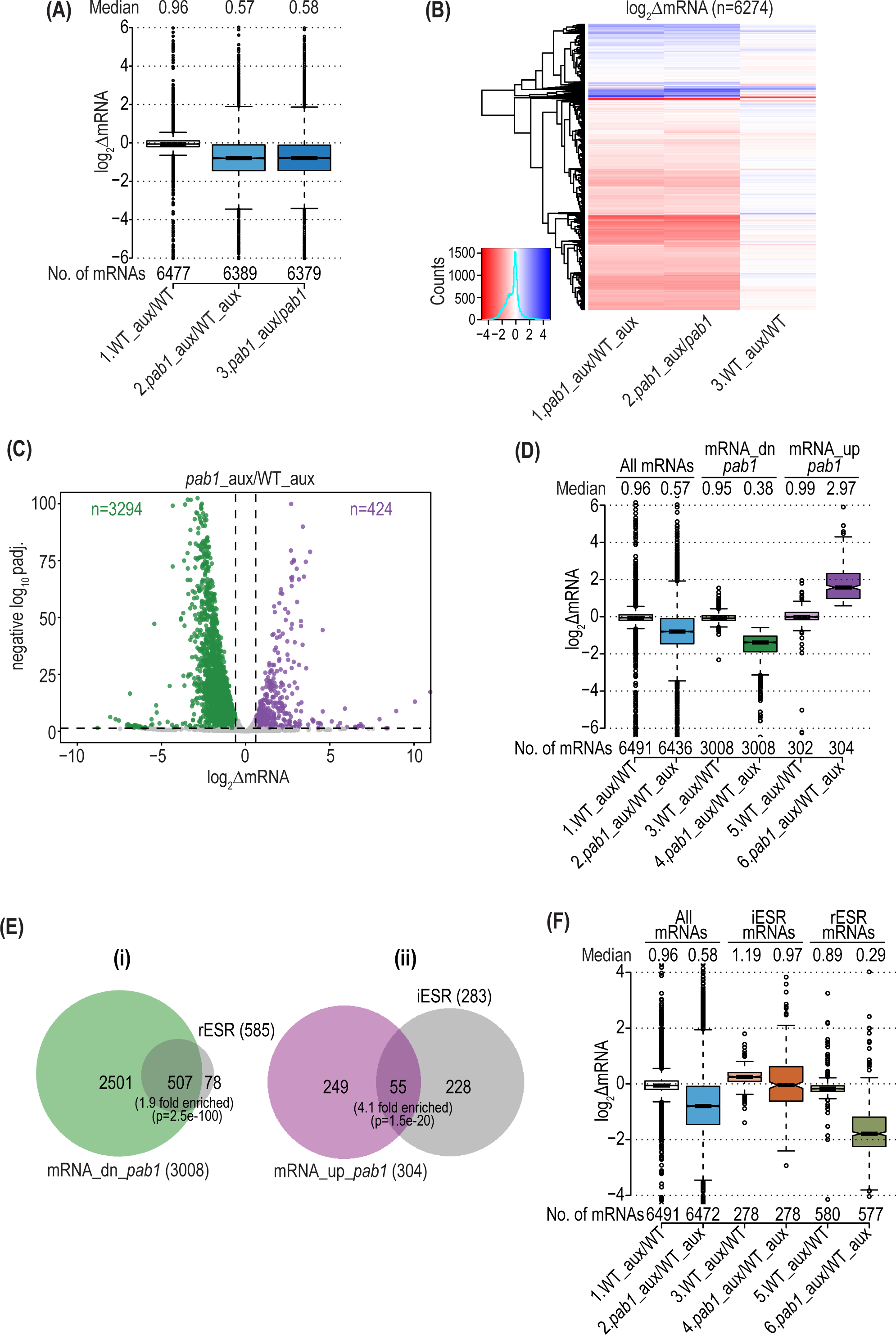
Pab1 depletion confers widespread changes in mRNA abundance that greatly exceed the ESR. **(A)**Notched boxplot showing log_2_ fold-changes in spike-in normalized total mRNA abundance (ΔmRNA) from DESeq2 analysis of the RNA-Seq data for the WT *PAB1* strain (PMY1) treated vs. untreated with 1 mM NAA for 6 hr in YPD medium (WT_aux/WT), the *pab1-AID* mutant (PMY2) similarly treated with NAA or untreated (*pab1-*AID_aux/*pab1-AID*), and for NAA-treated *pab1-AID* vs. NAA-treated WT cells (*pab1_*aux/WT_aux). Unlogged median values and numbers of mRNAs for each group are indicated on the top and bottom, respectively. Outlier mRNAs (between 0-57 for different columns) with log_2_ΔmRNA values of >6.0 or <-6.0 were removed to expand the y-axis scale. In these and all other notched boxplots, non-overlapping notches in adjacent columns indicate that the two medians differ significantly with ∼95% or higher confidence. **(B)** Hierarchical clustering analysis of the log_2_ΔmRNA values from (A) ranging from 4 (strong derepression, dark blue) to −4 (strong repression, dark red). Outliers (217 mRNAs) with values >5.0 or <-5.0 were excluded to enhance color differences. **(C)** Volcano-plot of log_2_ΔmRNA values from (A) for *pab1_*aux/WT_aux vs. negative log_10_ of adjusted p-values (padj.) for the fold-changes determined by DESeq2 analysis of the spike-in normalized RNA-Seq data (y-axis). The dotted line marks the 5% padj. threshold for mRNAs showing a significant increase (mRNA_up*_pab1,* n=424) or decrease mRNA_dn*_pab1* n=3294) in mRNA abundance at padj. < 0.05, in purple and green dots, respectively. Outlier mRNAs (between 2-67) with -log10 padj. >100 and log_2_ΔmRNA values >10 or < -10 were excluded to expand the axes. **(D)** Boxplot of log_2_ΔmRNA values of all expressed mRNAs or the 304 mRNA_up*_pab1* and 3008 mRNA_dn*_pab1* groups defined in Fig. S1E for the comparisons along the x-axis, presented as in (A), removing outlier mRNAs (2–51) with values >6.0 or <-6.0. **(E)** Proportional Venn diagrams of overlaps between the mRNA_up*_pab1* and mRNA_dn*_pab1* groups from Fig. S1E and the induced (iESR, n=283) or repressed (rESRs, n=585) ESR mRNAs, respectively. Hypergeometric distribution p-values are shown for overlaps. (**F)** Boxplot of log_2_ΔmRNA values for all mRNAs, the iESR, and rESR mRNAs from (E) presented as in (A), removing between 0-129 mRNAs with values >4.0 or <-4.0.

In agreement with the slow-growth of the Pab1-AID-depleted cells (Fig. S1B), the down-regulated mRNAs include 87% of the 585 mRNAs belonging to the Environmental Stress Response, which are repressed by various stresses or mutations that impair cell growth and primarily encode proteins required for ribosome biogenesis and translation (45) (Fig. 1E(i), rESR transcripts). Depletion of Pab1 confers a lesser, but significant, up-regulation of many of the 283 induced ESR mRNAs (Fig. 1E(ii), iESR). The subset of down-regulated mRNAs belonging to the rESR shows a greater decrease in median mRNA abundance compared to all mRNAs on depletion of Pab1-AID (Fig. 1F, cols. 6 vs. 2). The group of iESR mRNAs shows no decrease in median mRNA abundance; however, because the median abundance of all mRNAs is decreased, the iESR transcripts exhibit a relative increase compared to all other transcripts on Pab1-AID depletion (Fig. 1F, cols. 4 vs. 2), which is consistent with induction of iESR transcripts on Pab1 depletion. While these results indicate that the ESR is mobilized, it is also apparent that most mRNAs down- or up-regulated by Pab1-AID depletion are not ESR transcripts (Fig. 1E; 2501 transcripts, (i); 249 transcripts, (ii)). This finding suggests a broader and more direct role for Pab1 in controlling mRNA abundance.

### Reductions in mRNA abundance on Pab1-AID depletion result primarily from increased decapping and degradation that preferentially targets short pA-tailed isoforms

We explored next the impact of depleting Pab1-AID on lengths of pA tails by conducting single-molecule pA tail sequencing (SM-PAT-Seq) (46) on *pab1-AID* and WT cells, both treated with auxin. The distributions and median pA tail lengths were highly reproducible among 3 biological replicates obtained for each strain for ∼5600 expressed genes (Fig. S2A, cols. 1-8), allowing us to combine the data from replicates for all downstream analyses. The results revealed a substantial increase in pA tail lengths throughout the transcriptome on depletion of Pab1-AID (Fig. 2A). (Henceforth results for auxin-treated *pab1-AID* or treated WT cells will be abbreviated *pab1* and WT, respectively.) While this finding agrees with a published analysis of aggregate pA tail lengths in total yeast mRNA from cells depleted of Pab1 (44), these previous measurements were likely biased towards the most highly abundant transcripts. Our results indicate that lengthening of pA tails applies to the great majority of all expressed mRNAs, with a median increase of 20 nt (Fig. 2B). The shift to longer pA tails on Pab1-AID depletion could result from diminished recruitment of deadenylases to pA tails, reducing the rate of deadenylation. Alternatively, as suggested previously for PABP-depleted mammalian cells (10), it could arise from preferential degradation of the short-tailed isoforms of most mRNAs owing to preferential loss of Pab1 binding to shorter pA tails at reduced Pab1 levels. This would occur if mRNAs with short pA tails compete poorly for limiting Pab1, a possibility supported by previous RIP-Seq data indicating that mRNAs with relatively high Pab1 occupancies tend to have longer than average pA tails in yeast (16). The loss of Pab1 from short-tailed isoforms should result in loss of Pab1-eIF4G interaction, diminishing eIF4G-eIF4E binding to the cap and accelerating decapping and 5’-3’ degradation (Fig. S2B(ii)).

**Fig. 2.**
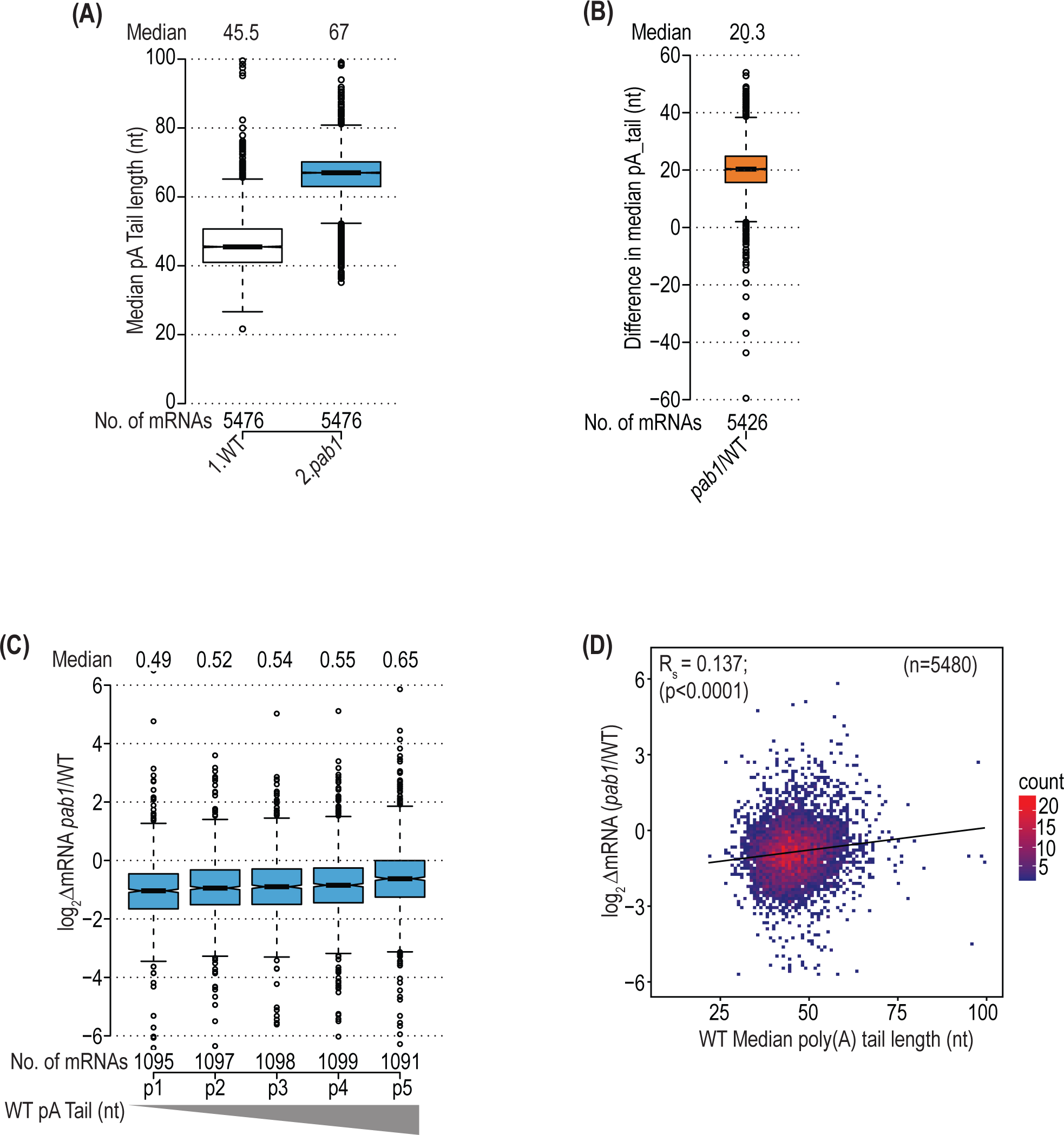
Pab1 depletion leads to pA tail shortening driven partly by preferential degradation of short-tailed transcripts. **(A)**Boxplot of median pA tail lengths (in nt) calculated for each of 5476 yeast genes in NAA-treated WT or treated *pab1-AID* cells, presented as in Fig. 1, with 4 or less outliers removed having median pA tail lengths >100 nt. **(B)** Boxplot of differences in median pA tail length for the genes in (A) in NAA-treated *pab1-AID*/WT cells, with 3 outliers removed having differences >60 nt. **(C)** Boxplot of spike-in normalized log_2_ΔmRNA changes in NAA-treated *pab1-AID*/WT cells for pentiles of all genes binned according to median pA tail lengths, progressing left to right from shortest to longest, in treated WT cells. Outliers (≤5) with values >6.0 or <-6.0 were removed. **(D)** Density scatterplot of spike-in normalized log_2_ΔmRNA changes versus median pA tail lengths in treated WT cells for 5480 genes, showing the Spearman correlation coefficient (R_s_) and its p-value. Outliers (0–18) with pA tail lengths >100 or log_2_ΔmRNA values >6.0 or <-6.0 were excluded to expand the axes.

Consistent with this explanation, we observed that mRNAs with shorter median pA tails in WT cells tend to show greater reductions in total mRNA abundance on Pab1-AID depletion (Fig. 2C). Thus, the pentile of ∼1100 transcripts of shortest pA tails in WT is reduced in median abundance by 0.49-fold, whereas the pentile with longest pA tails is reduced only 0.65-fold. This point was underscored by the fact that the median pA tail length of the mRNA_dn_*pab1* transcripts is smaller than that of the mRNA_up_*pab1* group in WT cells (Fig. S2C, cols. 2-3), and these differences are eliminated in *pab1* cells (Fig. S2C, cols. 5-6). Moreover, the change in mRNA abundance on depletion of Pab1-AID (log_2_ΔmRNA(*pab1/*WT)) shows a small but highly significant positive correlation with median pA tail length across the transcriptome (Spearman ρ = 0.137, P < 0.0001) (Fig. 2D). This trend would be expected if the short-tailed isoforms of most mRNAs undergo preferential decapping/decay on Pab1-AID depletion (Fig. S2B(ii)).

More compelling evidence supporting preferential degradation of short-tailed isoforms at limiting Pab1 came from investigating the consequences on transcript abundance and pA tail length of eliminating Dcp2, catalytic subunit of the decapping enzyme. Deletion of *DCP2* in the *pab1-AID* strain suppressed the effect on cell growth of auxin treatment (Fig. S3A, cf. *left* vs. *right* panels of rows 8 & 3), but *dcp2*Δ did not interfere with depletion of Pab1-AID protein (Fig. S3B, cf. +/- NAA in lanes 5-8 vs. 3-4). These results are consistent with previous findings that eliminating genes involved in decapping and 5’-3’ decay, including *DCP1* (encoding the regulatory subunit of decapping enzyme) and *XRN1*, suppresses the lethality of deleting *PAB1* (19,47). Accordingly, we conducted RNA-Seq on auxin-treated *dcp2Δpab1-AID* and *dcp2*Δ cells, obtaining highly reproducible results among biological replicates (Fig. S1G), and determined spike-in normalized changes in mRNA abundance for all expressed genes. Remarkably, the *dcp2*Δ mutation also rescued the reduction in bulk mRNA levels observed in *DCP2* cells on depletion of Pab1-AID (Fig. 3A) and dampened or eliminated the changes in abundance of most individual mRNAs across the transcriptome that occurred in auxin-treated *DCP2 pab1-AID* cells (Fig. 3B, cols 1-2). This effect reduced by ∼36-fold the number of mRNAs significantly down-regulated on Pab1-AID depletion in *dcp2*Δ versus *DCP2* cells (Fig. 3C(ii), 3294 vs. 92).

**Fig. 3.**
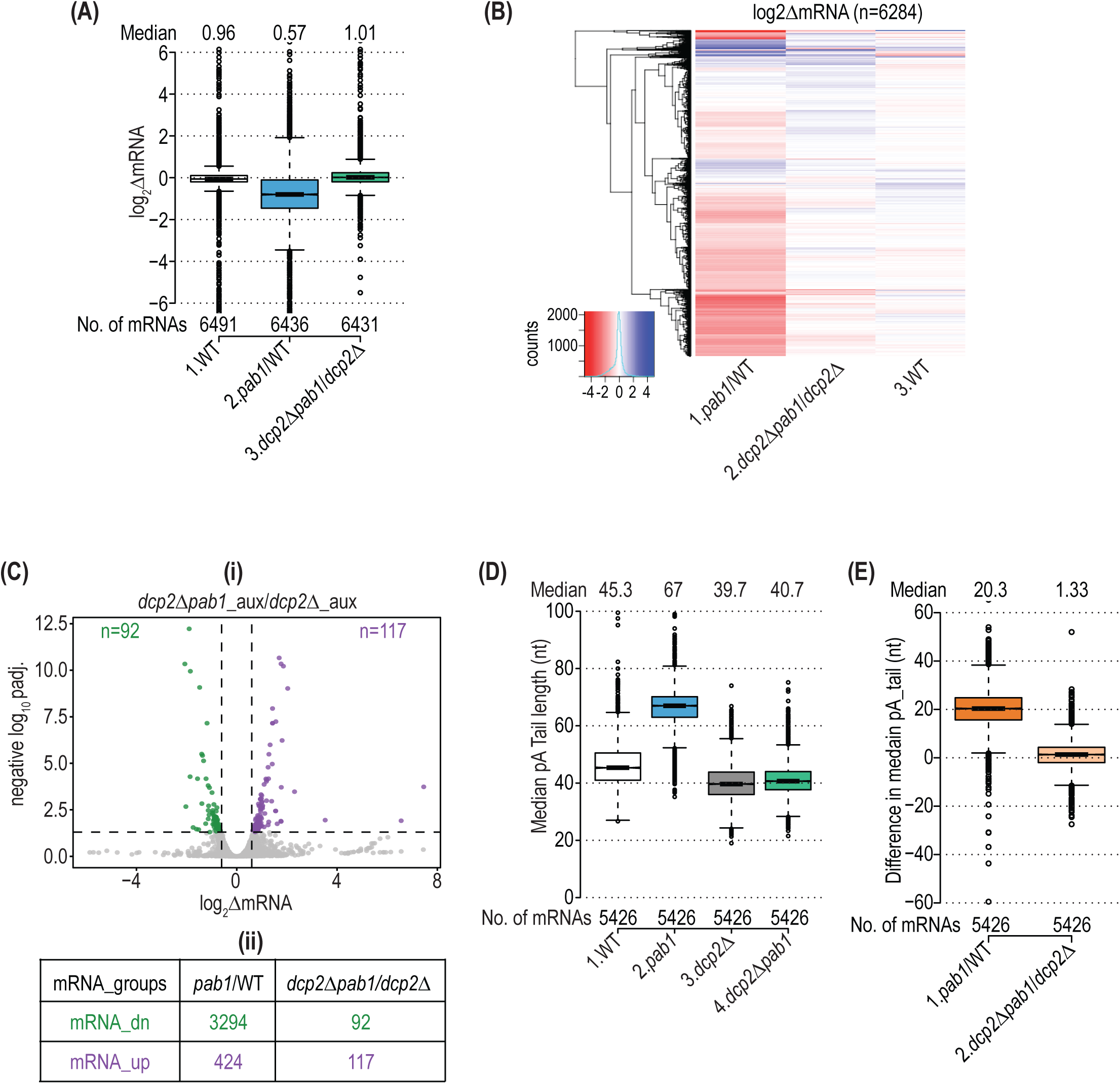
Eliminating *DCP2* suppresses reductions in mRNA abundance in Pab1-depleted cells. **(A)**Boxplot of spike-in normalized log_2_ΔmRNA changes in (1) NAA-treated vs. untreated WT, (2) treated *pab1-AID* versus treated WT cells, and (3) treated *dcp2Δpab1-AID* vs. treated *dcp2*Δ cells, with between 5-50 outliers with values >6.0 or < -6.0 removed. **(B)** Hierarchical clustering analysis of the data in (A) presented as in Fig. 1B, removing 100 mRNA outliers with values of >5.0 or < -5.0. **(C) (i)** Volcano-plot of spike-in normalized log_2_ΔmRNA changes in NAA-treated *dcp2Δpab1-AID* vs. treated *dcp2*Δ cells presented as in Fig. 1C. **(ii)** Numbers of mRNAs showing significant decreases (mRNA_dn) or increases (mRNA_up) in mRNA abundance in treated *pab1-AID/*WT (taken from Fig. 1C) or treated *dcp2Δpab1-AID*/*dcp2*Δ cells (from (i)). **(D)** Boxplot of median pA tail lengths for each of 5426 genes in (1) NAA-treated WT, (2) treated *pab1-AID* (3) treated *dcp2*Δ, and (4) treated *dcp2Δpab1-AID* cells, with 3-4 outliers removed with median pA tail lengths >100 nt. **(E)** Boxplot of differences in median pA tail length for the genes in (D) in (1) NAA-treated *pab1-AID*/WT cells and (2) treated *dcp2Δpab1-AID* vs. treated *dcp2*Δ cells, with 3 outliers removed having differences >60 nt.

Examining the smaller group of 92 mRNAs down-regulated by Pab1 depletion in *dcp2*Δ cells reveals that they are significantly enriched for the 3008 mRNA_dn_*pab1* transcripts down-regulated in *DCP2* cells (Fig. S3C). The 76 mRNAs common between these two groups (designated Group II in Fig. S3C) differ from the 2932 mRNAs down-regulated exclusively in *DCP2* cells (Group I) in showing relatively larger reductions in mRNA abundance on Pab1-AID depletion in *DCP2* cells (Fig. S3D, col. 5 vs. 3) and retaining marked reductions on Pab1-AID depletion in *dcp2*Δ cells (Fig. S3D, col. 6 vs. 4). One possibility is that the 76 Group II mRNAs are protected by Pab1 from degradation by both the cytoplasmic exosome and decapping enzyme in *DCP2* cells and thus undergo exosomal degradation even after eliminating Dcp2 in the *dcp2Δpab1-AID* double mutant. This would also explain why they show a relatively greater reduction on Pab1-AID depletion in *DCP2* cells where both degradation pathways are intact. The small group of 16 mRNAs in Group III of the Venn diagram in Fig. S3C are distinctive in being substantially down-regulated on Pab1-AID depletion only in the absence of Dcp2 in *dcp2*Δ cells (Fig. S3D, cols. 7-8). This unexpected behavior could be explained if these transcripts are targeted by the exosome at low levels of Pab1-AID only when the decapping machinery is absent. Despite this last suggestion that Pab1 might impede degradation by the exosome on a small fraction of transcripts, our results provide strong evidence that the widespread reductions in mRNA abundance conferred by depletion of Pab1-AID arise largely from decapping and attendant 5’-3’ degradation.

Having shown that eliminating *DCP2* strongly suppressed changes in mRNA abundance throughout the transcriptome, we went on to conduct SM-PAT-Seq on auxin-treated *dcp2Δpab1-AID* and *dcp2*Δ cells (Fig. S2C). Remarkably, *dcp2*Δ completely abrogated the lengthening of pA tails conferred by Pab1-AID depletion in *DCP2* cells (Fig. 3D, cols. 3-4 vs. 1-2; Fig. 3E). This result strongly supports the notion that the widespread shift to longer pA tails on Pab1-AID depletion results from preferential decapping and degradation of short-tailed transcript isoforms, which is eliminated in *dcp2*Δ cells, rather than from diminished recruitment of deadenylases to the pA tail at low Pab1 levels.

### Properties of mRNAs dysregulated in abundance by Pab1-AID depletion

The results in Figs. 2C-D imply that relatively short mean pA tail length in WT cells is a determinant of preferential mRNA degradation on Pab1 depletion. However, the correlations shown there are relatively small and mRNAs with the longest mean pA tails still undergo marked reductions in mRNA levels on Pab1-AID depletion (Fig. 2C, pentile 5), indicating that a short pA tail is not the only determinant of preferential decapping and turnover at limiting Pab1 levels. We considered whether the propensity of mRNAs to form the closed-loop intermediate might also be involved, such that mRNAs preferentially down-regulated at limiting Pab1 would assemble this intermediate very inefficiently. As noted above, mRNAs with higher-than-average occupancies of Pab1, eIF4G and eIF4E and lower than average occupancies of 4E-BPs, consistent with strong closed-loop (SCL) assembly, tend to be highly translated and have shorter than average CDS lengths (16,48) (Figs. S4B-C, col. 4 vs. 1), a property that appears to facilitate the closed-loop (49). As shown in Figs. S4A-C, the properties of the 3008 mRNAs down-regulated on Pab1-AID depletion in *DCP2* cells (mRNA_dn_*pab1*) seem incompatible with unusually inefficient closed-loop assembly, as they exhibit: (i) only ∼10% lower-than average median occupancies of eIF4G and eIF4E and essentially average occupancies of Pab1 and the inhibitory 4E-BPs (Fig. S4A, green or brown vs. grey boxes); (ii) only ∼10% lower than average median CDS lengths (Fig. S4B, cols. 1-2); and (iii) only ∼17% higher than average median TE in WT cells (Fig. S4C, cols. 1-2). (See summary in Table 1, rows 1, 3, 7-11). Furthermore, these mRNAs are moderately enriched for, rather than depleted of, the set of 395 SCL mRNAs identified by Costello et al. (Fig. S4D). Thus, a low propensity for forming the closed loop does not appear to be a key determinant of the preferential degradation of the mRNA_dn_*pab1* group of transcripts at limiting Pab1 levels.

**Table 1:**
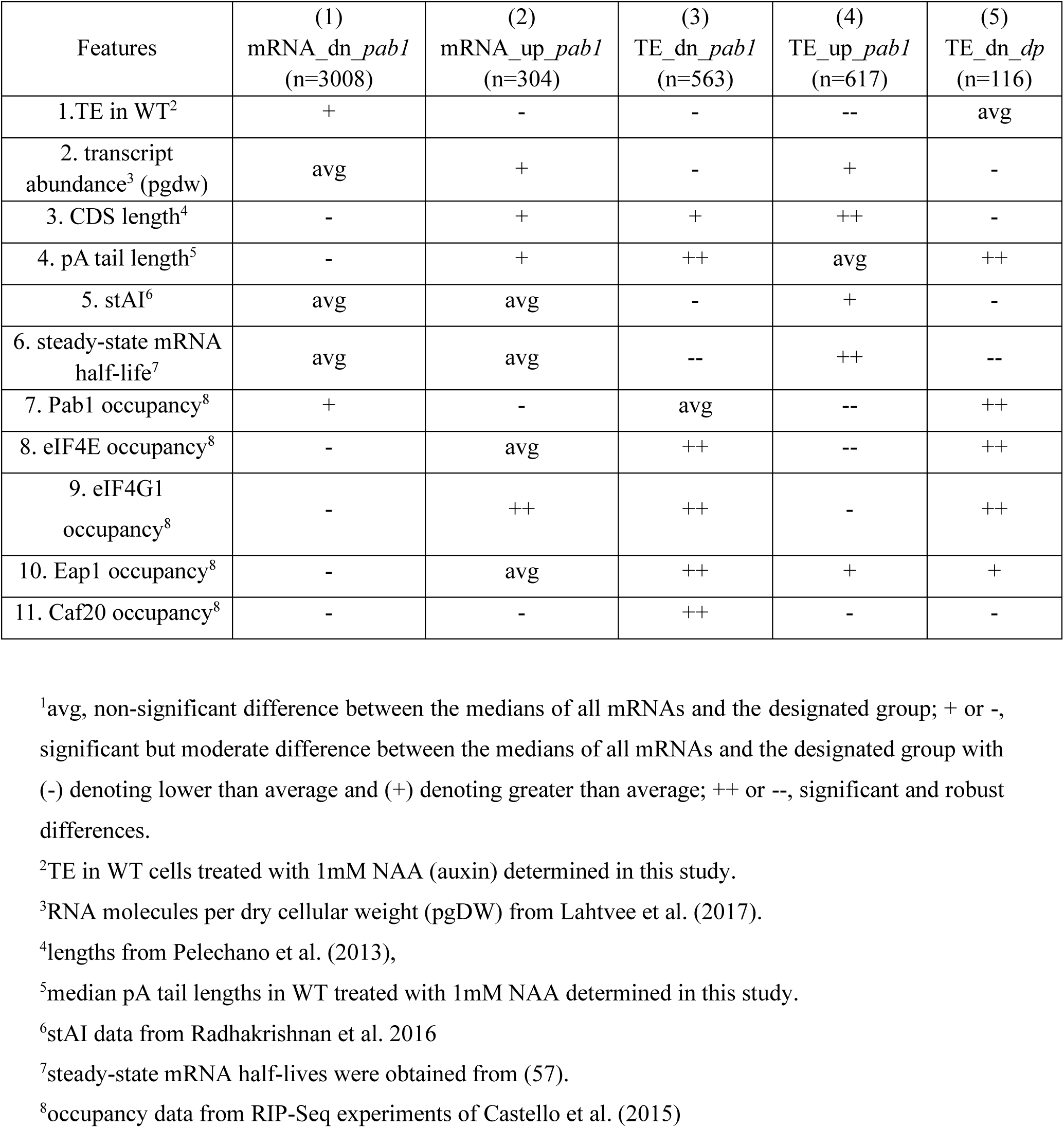
A comparison of mRNA features between different mRNA groups^1^.

It was shown recently for mammalian somatic cell lines (11) that the mRNAs whose abundance decreases on PABP depletion tend to be more stable and have shorter than average 5’UTRs, 3’UTRs, and pA tail lengths in non-depleted cells. In contrast, the mRNA_dn_*pab1* transcripts have slightly less than average stability (Fig. S4E) and shorter than average pA tails (Fig. S2B), but average 5’UTR and 3’UTR lengths in WT yeast (Fig. S4F-G); and they have essentially average codon optimality (Fig. S4H, cols. 1-2) and transcript abundance (S4I, cols.1-2). (See summary in Table 1, rows 2,4,5,6). Overall, the mRNAs down-regulated at limiting Pab1-AID levels do not have atypical features suggestive of a particular mechanism underlying their preferential targeting for degradation.

### Depletion of Pab1-AID preferentially diminishes both histone mRNAs and histone proteins and derepresses cryptic transcripts

Gene ontology analysis of the 3008 mRNA_dn_*pab1* transcripts revealed a significant enrichment for genes involved in regulation of transcription and chromatin modifications (P = 3.2 x 10^-14^ and 6.5 x 10^-7^, respectively). Interestingly, we observed that the mRNAs transcribed from the 11 histone genes showed a dramatic reduction on Pab1-AID depletion, much greater than that observed for all mRNAs (Fig. 4A, col. 5 vs. 2). To examine whether the reductions in histone mRNAs lead to reduced histone protein levels, we conducted TMT-mass spectrometry of all expressed proteins in auxin-treated *pab1-AID* vs. WT cells, obtaining highly reproducible results between biological replicates (Fig. S6E). Indeed. we observed reduced expression of histone proteins relative to all other cellular proteins detected in the experiment (Fig. 4A, cols. 9 vs. 7). Importantly, the decreased abundance of histone mRNA and protein conferred by *pab1-AID* was substantially suppressed by *dcp2*Δ (Fig. 4A, cols. 5-6 and 9-10), indicating that it results primarily from elevated decapping and degradation of these transcripts at low Pab1 levels.

**Fig. 4.**
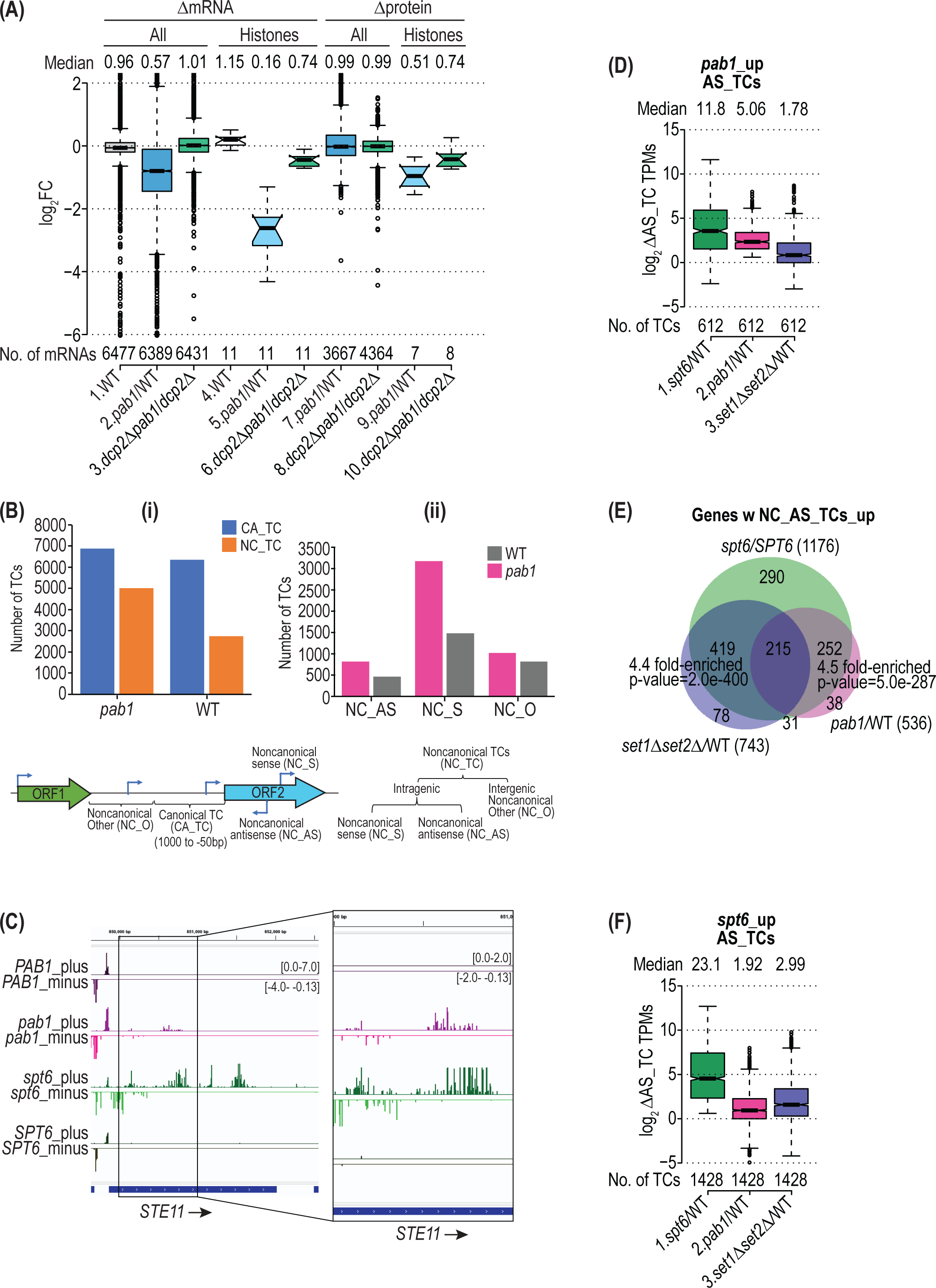
Pab1 depletion leads to preferential degradation of histone mRNAs, reduced histone proteins, and derepressed internal cryptic promoters. **(A)**Boxplot of spike-in normalized log_2_ΔmRNA changes (columns 1-6) and log_2_ relative fold-changes in protein abundance determined by TMT-MS (columns 7- 10) for all mRNAs or histone mRNAs/proteins in (1 and 4) NAA-treated vs. untreated WT cells, (2,5,7 and 9) treated *pab1-AID* versus treated WT cells, and (3, 6, 8 and 10) treated *dcp2Δpab1-AID* vs. treated *dcp2*Δ cells, with outliers of between 0-220 mRNAs with values >2.0 or < -6.0 removed. **(B)** Histograms of numbers of transcription start site clusters (TCs) from CAGE data (with >1 TPM) for (i) all canonical (CA) and all non-canonical (NC) TCs, and (ii) NC TCs that are intragenic antisense (NC_AS), intragenic sense (NC_S), or “other” intergenic transcripts (NC_O) based on their locations in NAA-treated WT or treated *pab1-AID* cells. Nomenclature and coordinates of TC classes are schematized below. **(C)** Screenshot of the IGV Genome Browser showing CAGE signals (TPM) for plus and minus strands for the *STE11* gene in NAA-treated *pab1-AID* or treated WT *PAB1* cells and for untreated *spt6-1004* and isogenic WT *SPT6* cells. The dotted rectangle encompasses NC_AS and NC_S TCs overlapping between *pab1-AID* mutant and *spt6-1004* cells. All rows are scaled equally with the ranges in brackets for the plus and minus strands. **(D)** Boxplots showing log_2_ fold-change in CAGE TPMs for the 612 NC_AS TCs upregulated by NAA-treatment of *pab1-AID* cells (mapping in 563 genes) in untreated *spt6-1004/SPT6,* NAA-treated *pab1-AID*/WT, or untreated *set1Δset2*Δ/WT cells determined by DESeq2 analysis of the CAGE data for each comparison. The *spt6-1004* mutant FY2180*, set1Δset2*Δ mutant H4293, and corresponding isogenic WT strains F2181 and BY4741, respectively, were cultured without NAA in SC medium at 30°C for *set1Δset2*Δ and WT control cells and at 30°C followed by a 90 min shift to 39° for *spt6-1004* and WT control cells. **(E)** Proportional Venn diagrams of overlaps between genes containing NC_AS TCs significantly upregulated ≥1.5-fold (determined by DESeq2 analysis of the CAGE data) in NAA-treated *pab1-AID/*WT cells (n=536), untreated *spt6-1004* vs WT cells (n=1176), or untreated *set1Δset2*Δ vs WT cells (n=743) among all 6028 genes containing TCs of any category. Hypergeometric distribution p-values are shown for overlaps between the genes containing AS_TCs derepressed by *spt6-1004* (n=1176) and either *set1Δset2*Δ (n=743) or NAA-treatment of *pab1-AID* cells (n=536). **(F)** Boxplots showing log_2_ fold-change in CAGE TPMs for the 1428 NC_AS TCs upregulated by *spt6-1004* (mapping in 1176 genes) in *spt6-1004/*WT, NAA-treated *pab1-AID/*WT, or *set1Δset2Δ/*WT cells determined by DESeq2 analysis of the CAGE data for each comparison.

A known cellular consequence of depleting histones is the activation of cryptic internal promoters owing to reduced nucleosome densities within the coding sequences of many genes (50,51). To determine whether the depletion of histones on Pab1-AID depletion is large enough to confer a significant reduction in nucleosome densities, we conducted CAGE analysis to identify the transcription start sites (TSSs) of all expressed mRNAs, including those mapping within CDSs. Transcription in yeast generally initiates at a cluster of nearby TSSs, which can be computationally grouped as a “TSS cluster” (TC), representing the location of a putative core promoter (52). Transcription initiation activity from a core promoter can be quantified as the number of CAGE tags mapped within a TC normalized by the sequencing depth of a sample (CAGE tags per million mapped tags, or TPMs). As only one CAGE read is generated for each mRNA transcript, the CAGE TPM is equivalent to transcripts per million and can be compared between different conditions or yeast strains (52). We conducted CAGE analysis (32) on the *pab1-AID* and WT strains both treated with auxin and obtained highly reproducible results between the biological replicates (Fig. S5A). Notably, depletion of Pab1-AID led to a marked increase in total number of TCs throughout the genome that was more pronounced for “non-canonical” TCs (NC_TC), those mapping within CDSs or >1000 nt upstream of CDSs, compared to canonical TCs mapping between 50 and 1000 nt upstream of CDSs (Fig. 4B(i)). Furthermore, among the NC_TCs, the increase is especially large for those mapping within CDSs, including both sense (NC_S) and antisense (NC_AS) orientations (Fig. 4B(ii)). Increased CAGE TPMs in *pab1-AID* cells were observed on both the sense and antisense strands within the CDS of *STE11* (Fig. 4C, *pab1* vs *PAB1*), a gene previously shown to exhibit activation of internal cryptic promoters in mutant cells lacking Spt6, a histone chaperone required for proper occupancies of genic nucleosomes (50). This phenomenon was widespread, as differential expression analysis of the CAGE data using DESeq2 identified 612 AS TCs in the CDSs of 536 genes whose CAGE TPMs were significantly derepressed by *pab1-AID* (Fig. 4D, col. 2), of which four representative genes (*CSG2, TCD1, NRP1,* and *BAT2*) are shown in Fig. S5B-E (*pab1* vs *PAB1*).

If the increase in intragenic AS transcripts on Pab1-AID depletion results from nucleosome depletion, it would be expected to occur at many of the same genes in cells with impaired Spt6 function. To test this prediction, we obtained CAGE data from a *spt6-1004* mutant and isogenic wild-type *SPT6* strain (Fig. S5A). A relatively greater number of AS TCs (1428 TCs) were significantly derepressed in *spt6-1004* vs. *SPT6* cells, which encompasses 85% of the 612 AS TCs described above derepressed on Pab1-AID depletion. Importantly, the 536 genes containing derepressed AS transcripts in *pab1-AID* cells were highly enriched for the 1176 genes containing AS TCs derepressed by *spt6-1004* (Fig. 4E, green vs. red gene sets, P = 5 x 10^-287^). This overlap is further illustrated by examining the locations of AS TCs derepressed by the two mutations at particular genes, wherein those up-regulated by *pab1-AID* represent a subset of the AS TCs up-regulated by *spt6-1004.* This pattern is evident at *STE11* and four other representative genes, which also exhibit derepression of subgenic transcripts from the sense strand (NC_S) that initiate at overlapping positions within the CDSs in these two mutants (Figs. 4C and S5B-E, cf. *pab1* vs. *spt6*). Consistent with these findings, both groups of the 612 AS TCs derepressed by *pab1-AID* and the 1428 TCs derepressed by *spt6-1004* show significantly increased median abundance in both mutants that is relatively greater in magnitude for *spt6-1004* (Figs. 4D & 4F, cols. 1-2).

In addition to mutants with reduced nucleosome density, cells lacking co-transcriptional histone H3 methylation catalyzed by Set1 or Set2 also exhibit activation of cryptic internal promoters within CDSs (53). Accordingly, we extended our analysis to include CAGE data obtained from a *set1Δset2*Δ double mutant and isogenic WT strain (Fig. S5A). We observed highly significant enrichment for the genes containing AS TCs derepressed by *set1Δset2*Δ and those containing AS TCs up-regulated by *spt6-1004* or *pab1-AID* (Fig. 4E) as well as significant increases in median AS TC abundance in all three mutants that is greatest in magnitude for *spt6-1004* (Figs. 4D & 4F). In summary, CAGE analysis of AS transcription provides evidence that depletion of Pab1-AID reduces genic nucleosome occupancies sufficiently to activate cryptic internal promoters at many genes where these promoters are normally repressed by Spt6, Set1, or Set2.

### Depletion of Pab1 leads to widespread changes in translational efficiencies

We turned next to consider the consequences of depleting Pab1 on the translatome by polysome analysis and Ribo-Seq of the *pab1-AID* mutant. Consistent with previous findings (44), depletion of Pab1-AID leads to a strong reduction in bulk translation as demonstrated by a depletion of total polysomes and accumulation of 80S monosomes to reduce the polysome:monosome (P/M) ratio by ∼3-fold in auxin-treated *pab1-AID* vs. WT cells (Fig. 5A). We next conducted ribosome profiling on auxin-treated and untreated *pab1-AID* and WT cells, obtaining highly correlated results for three biological replicates of each strain/condition (Fig. S6A). In contrast to observations in PABP-depleted mammalian cells (10,11), we observed a substantial reprogramming of translation on Pab1-AID depletion when comparing auxin-treated *pab1-AID* and WT cells, involving decreased relative TEs of 735 mRNAs and increased relative TEs of 864 transcripts (ΔTE ≥ 1.4-fold, FDR ≤ 0.1) (Fig. 5B). A similar number of transcripts, 700 and 777 respectively, showed decreased or increased relative TEs in comparing auxin-treated and untreated *pab1-AID* cells (Fig. S6B). As the two groups of similarly dysregulated transcripts overlapped substantially, we restricted our attention to the 563 and 617 transcripts identified in both comparisons, designated TE_dn_*pab1* and TE_up_*pab1*, respectively (Fig. S6C). These groups displayed median fold-changes in TE of 0.53-0.54 and 1.68-1.75, respectively, on comparing either *pab1-AID* and WT cells both treated with auxin (Fig. 5C) or *pab1-AID* cells with or without auxin treatment (Fig. S6D).

**Fig. 5.**
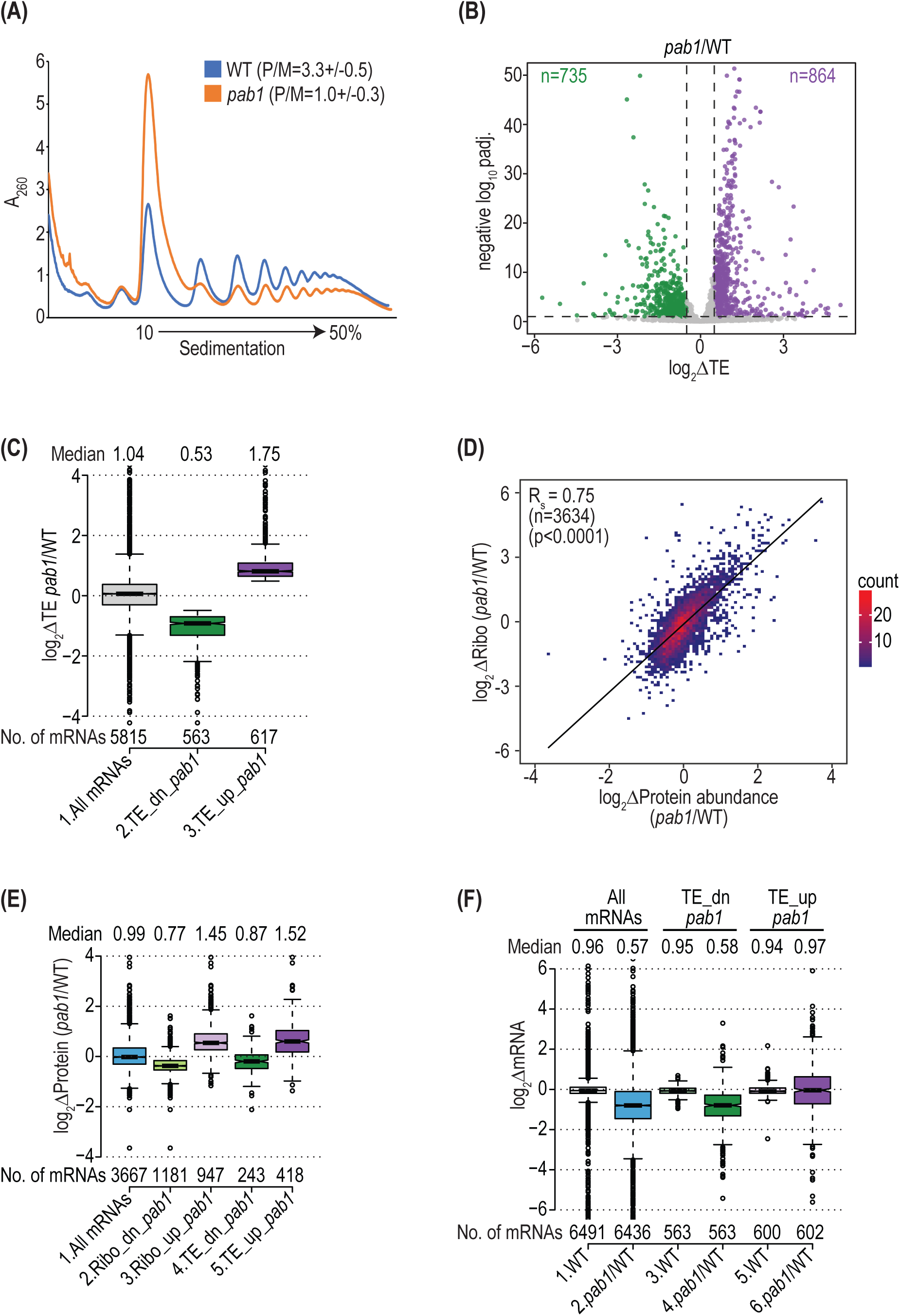
Pab1 depletion downregulates bulk translation and reprograms translational efficiencies of many genes. **(A)**Polysome profiles of WT and *pab1-AID* strains, both treated with NAA and cultured under the same conditions used for RNA-Seq and Ribo-Seq except that 50 mg/mL cycloheximide was added to the culture 5 min before harvesting at 4°C. Whole cell extracts were resolved by sedimentation through 10-50% sucrose gradients and scanned at 260 nm to visualize (from left to right) free 40S and 60S subunits, 80S monosomes, and polysomes. The polysome to monosome ratio was calculated from the corresponding areas under the tracing and mean ratios +/- SEM determined from 3 biological replicates are given above the representative plots shown for one replicate of each genotype. **(B)** Volcano-plot showing log_2_ fold-changes in relative translation efficiencies (ΔTE) for all 5816 yeast mRNAs in NAA-treated *pab1-AID* vs. treated WT cells vs. the -log_10_ padj. values for ΔTE changes determined by DESeq2 analysis of the profiling data (y-axis). The dotted line marks the 10% padj. threshold for mRNAs showing a significant increase (TE_up_*pab1,* n=864) or decrease of >1.4-fold (TE_dn_*pab1,* n=735) in ΔTE, plotted in purple and green, respectively. 12 outlier mRNAs with -log_10_ padj. >50 were excluded. **(C)** Boxplot of log_2_ΔTE values for NAA-treated *pab1-AID* vs. treated WT cells for all expressed mRNAs or the TE_up*_pab1* and TE_dn*_pab1* groups defined in Fig. S6C, removing 15, 4 and 5 outlier mRNAs respectively from col.1, 2 and 3 with log_2_ΔTE values >4.0 or < -4.0. **(D)** Density scatterplot of log_2_ fold-changes in relative numbers of ribosome protected fragments (RPFs) (log_2_ΔRibo) and log_2_ fold-changes in relative protein abundance measured by TMT-MS (log_2_ΔProtein) for 3634 yeast mRNAs in NAA-treated *pab1-AID* vs. treated WT cells, showing the Spearman correlation coefficient (R_s_) and its p-value. Two outliers with log_2_ΔmRNA values >6.0 were excluded. **(E)** Boxplot of log_2_ΔProtein values in NAA-treated *pab1-AID* vs. treated WT cells for (1) all mRNAs, (2-3) mRNAs showing ≥1.5-fold increases (Ribo_up_*pab1*) or decreases (Ribo_dn_*pab1*) at padj. <0.05 in NAA-treated *pab1-AID/*WT cells, and (3-4) the TE_up_*pab1* and TE_dn_*pab1* mRNAs defined in Fig. S6C. **(F)** Boxplot of spike-in normalized log_2_ΔmRNA changes for (1-2) all mRNAs, (3-4) the TE_up_*pab1* mRNAs, and (5-6) the TE_dn_*pab1* mRNAs defined in Fig. S6B. Outlier mRNAs (15 from col.1 and 51 from col.2) with values >6.0 or < -6.0 were removed.

To determine whether the changes in RPFs measured by ribosome profiling produce corresponding changes in rates of protein synthesis that alter the steady-state yeast proteome, we examined the results of TMT-mass spectrometry from auxin-treated *pab1-AID* vs. WT cells (Fig. S6E). Importantly, we found a strong positive correlation between relative changes in RPFs and relative changes in protein abundance determined by TMT-MS (Fig. 5D). Moreover, the groups of mRNAs displaying significantly up-regulated or down-regulated RPFs in auxin-treated *pab1-AID* vs WT cells, as well as the TE_up_*pab1* and TE_dn_*pab1* groups just described, exhibit significant changes in median protein abundance in the same directions (Fig. 5E). These last results indicate that changes in translation (RPFs) or TE on Pab1-AID depletion revealed by ribosome profiling generally lead to corresponding changes in steady-state protein abundance throughout the translatome. This correspondence implies in turn that increases or decreases in ribosome density on mRNAs (ΔTEs) reflect increases or decreases in rates of translation initiation on Pab1-AID depletion.

The TE_dn_*pab1* mRNAs exhibit a reduction in median absolute transcript levels of 0.58 on Pab1-AID depletion, similar in magnitude to that noted above for all transcripts (Fig. 5F col. 4 vs 2). In contrast, the TE_up_*pab1* transcripts show no change in median absolute mRNA abundance (Fig. 5F col. 6), which implies that they exhibit increased transcript levels relative to the average transcript in response to Pab1-AID depletion. We suggest below that the increased relative concentrations of the TE_up_*pab1* transcripts contributes to their increased relative TEs on Pab1 depletion.

### Reductions in mRNA abundance drive TE changes on Pab1-AID depletion

As described above, deletion of *DCP2* strongly suppressed reductions in transcript levels conferred by depletion of Pab1-AID. Remarkably, *dcp2*Δ also reversed the effects of Pab1-AID depletion on polysome assembly, restoring the P/M ratio to essentially WT levels (Fig. 6A). This finding suggests that the reduced polysome assembly conferred by Pab1-AID depletion results largely from diminished mRNA levels rather than loss of Pab1 function in stimulating translation initiation. Accordingly, we next conducted ribosome profiling on auxin-treated *dcp2Δpab1-AID* and *dcp2*Δ cells, obtaining highly reproducible results among biological replicates (Fig. S7A). Importantly, the *dcp2*Δ mutation also diminished the translational reprogramming observed on Pab1-AID depletion in *DCP2* cells, reducing the numbers of translationally down-regulated and up-regulated mRNAs by ∼4.8-fold and ∼2.5-fold, respectively (Fig. 6B(i)-(ii)). Furthermore, clustering analysis indicates that most of the transcripts who’s relative TEs are significantly up- or down-regulated on Pab1-AID depletion in *DCP2* cells show dampened TE changes in cells lacking Dcp2 (Fig. 6C, generally lighter red or blue hues in *dcp2*Δ vs. *DCP2* cells). Consistent with this, the median TE changes on Pab1-AID depletion are reduced by *dcp2*Δ from 0.53- to 0.88-fold for the TE_dn_*pab1* mRNAs and from 1.75- to 1.23-fold for the TE_up_*pab1* transcripts (Fig. 6D). Because *dcp2*Δ attenuates changes in both TEs and mRNA abundance elicited by Pab1-AID depletion, it seems likely that a large component of the translational reprogramming observed at limiting Pab1 in *DCP2* cells is an indirect consequence of concurrent changes in mRNA abundance.

**Fig. 6.**
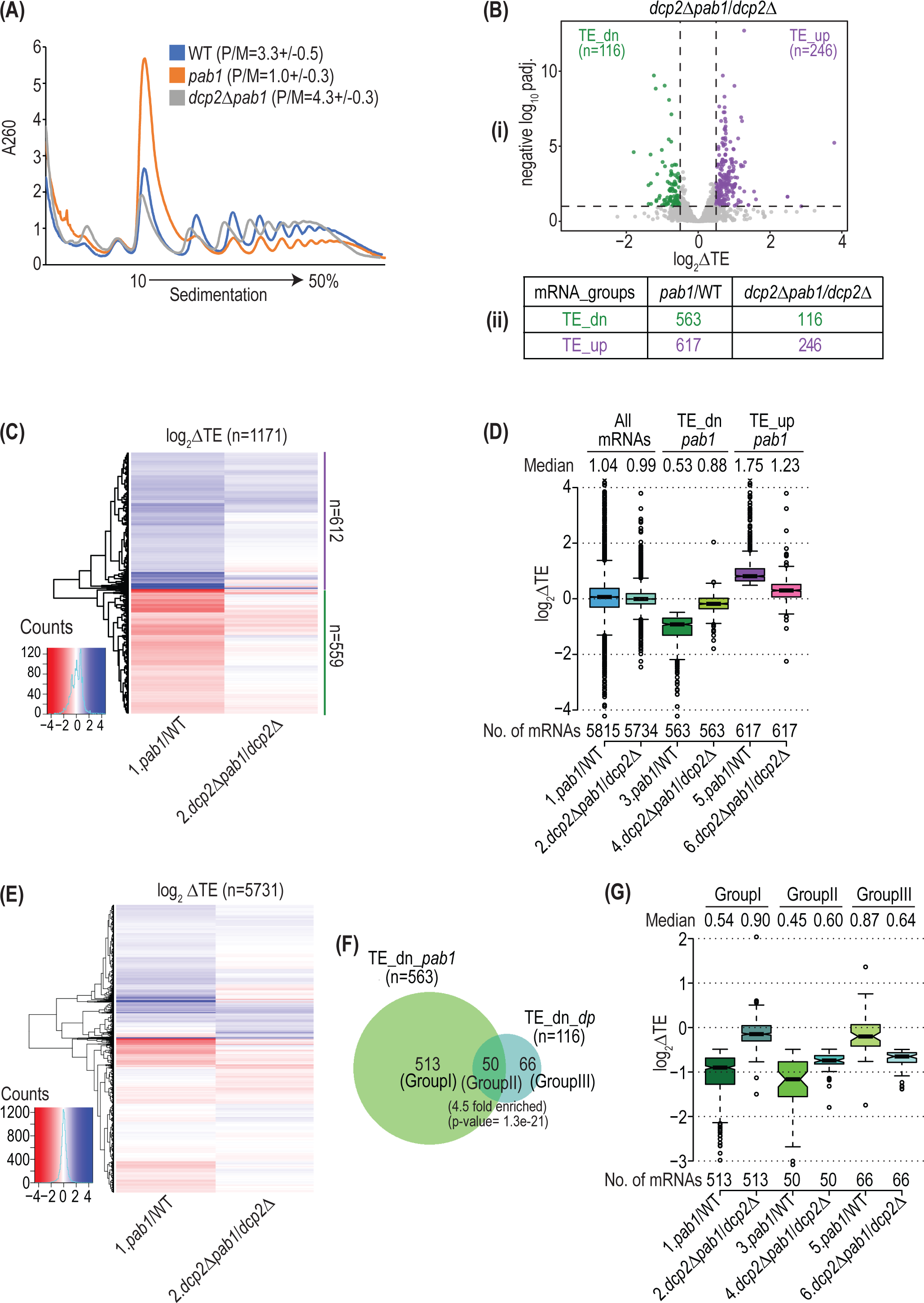
Eliminating *DCP2* suppresses reductions in bulk translation and TE changes in Pab1-depleted cells. **(A)**Polysome profiles of WT, *pab1-AID* mutant and *dcp2Δpab1-AID* mutant strains all treated with NAA conducted and analyzed as in Fig. 5A. **(B)** Volcano-plot of log_2_ΔTE values for all 5836 yeast mRNAs in NAA-treated *dcp2Δpab1-AID* vs. treated *dcp2*Δ cells presented as in Fig. 5B. **(C)** Hierarchical clustering analysis of log_2_ΔTE values for 1171 of the 1180 TE_dn_*pab1* or TE_up_*pab1* mRNAs analyzed in Fig. 5C observed in (1) NAA-treated *pab1-AID/*WT or (2) NAA-treated *dcp2Δpab1-AID/dcp2*Δ cells, presented as in Fig. 1B. Nine Outlier mRNAs with log2ΔTE >4.0 or < -4.0 were excluded. The positions of the TE_up_Pab1 (n=612) and TE_dn_Pab1 (n=559) mRNAs analyzed in Fig. 5C are indicated with purple and green lines, respectively, to the right of the plot. **(D)** Boxplot of log_2_ΔTE values for (1-2) all mRNAs, (3-4) the TE_up_Pab1 (n=617) and (5-6) TE_dn_Pab1 (n=563) mRNAs, analyzed in Fig. 5C, in NAA-treated *pab1-AID* vs. treated WT cells (1, 3, and 5) and in NAA-treated *dcp2Δpab1-AID* vs. treated *dcp2*Δ cells (2, 4, and 6). Outlier mRNAs (0-15) with log2ΔTE values of >4.0 or < -4.0 were removed. **(E)** Hierarchical clustering analysis of log_2_ΔTE values for all 5732 mRNAs observed in (1) NAA-treated *pab1-AID/*WT or (2) NAA-treated *dcp2Δpab1-AID/dcp2*Δ cells, presented as in Fig. 1B. Outlier mRNAs (85) with log2ΔTE >5.0 or <-5.0 were excluded. **(F)** Proportional Venn diagrams showing overlaps between the 563 TE_dn_*pab1* mRNAs from Fig. 5C and the 116 TE_dn *dcp2Δpab1* mRNAs from (B). Hypergeometric distribution p-value is shown for the overlap yielding transcripts designated Group II. **(G)** Boxplot of log_2_ΔTE values for the three mRNA groups defined in (F) in NAA-treated *pab1-AID* vs. treated WT cells (1, 3, and 5) and NAA-treated *dcp2Δpab1-AID* vs. treated *dcp2*Δ cells (2, 4, and 6). Outlier mRNAs (9 for col.1) with log2ΔTE >2.0 or <-3.0 were removed.

Interestingly, the hierarchical clustering analyses of fold-changes in TE in Fig. 6C revealed subsets of mRNAs that did not exhibit a marked dampening of TE changes on Pab1-AID depletion in *dcp2*Δ cells. Furthermore, a similar clustering analysis for all mRNAs revealed additional small groups of transcripts that were down- or up-regulated in TE only in *dcp2*Δ cells (Fig. 6E), suggesting that they are regulated more directly by Pab1. About 43% of the 116 mRNAs translationally down-regulated on Pab1-AID in *dcp2*Δ cells belong to the larger group of 563 mRNAs translationally down-regulated in *DCP2* cells (Group II of Fig. 6F). These 50 transcripts differ from the 513 mRNAs down-regulated exclusively in *DCP2* cells (Group I of Fig. 6F) in showing a much smaller dampening of TE reductions on Pab1-AID depletion conferred by *dcp2*Δ and, hence, substantial TE reductions in both *dcp2*Δ and *DCP2* cells (Fig. 6G, cols. 3-4 vs. 1-2). This weaker dampening for the Group II mRNAs is not due to lesser stabilization of their mRNA levels by *dcp2*Δ compared to Group I mRNAs (Fig. S7B, cols. 5-6 vs. 3-4). Thus, it appears that the Group II mRNAs are translationally down-regulated on Pab1-AID depletion in *DCP2* cells owing to loss of a direct stimulatory function of Pab1 combined with an indirect TE reduction arising from decreased mRNA abundance at limiting Pab1 levels. The remaining 66 mRNAs in Group III of Fig. 6F are distinctive in being translationally down-regulated by Pab1-AID depletion to a greater extent in *dcp2*Δ vs. *DCP2* cells (Fig. 6G, cols. 5-6). Thus, translation of these mRNAs appears to be dependent on a direct function of Pab1 in a manner exacerbated by eliminating the Dcp2-dependent changes in mRNA abundance that accompany Pab1-AID depletion. By conducting TMT-MS analysis (Fig. S7D), we determined that the changes in translation revealed by altered RPFs in *dcp2Δpab1-AID* versus *dcp2*Δ cells were highly correlated with changes in protein expression for the corresponding mRNAs measured by TMT-MS (Fig. S7C), supporting a role for Pab1 in translational reprogramming independent of its function in controlling mRNA abundance.

## DISCUSSION

In this study, we used a multi-omics approach to reveal the roles of yeast Pab1 in controlling the abundance and translation of individual mRNAs throughout the transcriptome and translatome and to evaluate the contribution of mRNA decapping to the influences of Pab1 on mRNA turnover and translation. Depletion of AID-tagged Pab1 by auxin addition to cells conferred a marked reduction in absolute levels of most mRNAs, decreasing median transcript abundance by a factor of ∼0.59. Yeast Pab1 is essential for viability, and in keeping with the impaired growth of Pab1-depleted cells, the stereotypical ESR was mobilized (54); however, most (∼84%) of the mRNAs dysregulated by Pab1-AID depletion are not ESR transcripts and likely controlled more directly by Pab1. Supporting this inference, a striking result was that changes in mRNA abundance conferred by depleting Pab1-AID were essentially abolished in cells lacking Dcp2, catalytic subunit of the decapping enzyme. This finding implies that low Pab1 abundance destabilizes most mRNAs by increasing susceptibility to decapping and subsequent 5’-3’ degradation by Xrn1. These findings support the notion that protecting mRNAs from degradation represents at least one essential function of Pab1 (19,47).

Single-molecule pA tail sequencing revealed a substantial increase in pA tail lengths throughout the transcriptome on Pab1-AID depletion, with a median increase of ∼20 nt. Strikingly, this effect was also eliminated by deleting *DCP2,* suggesting that it results from preferential degradation of short pA-tailed isoforms of most transcripts (Fig. S2B). The same conclusion was reached from analyzing consequences of PABP depletion on pA tail lengths in mammalian cells and attributed to preferential dissociation of PABP from short-tailed isoforms, leading to terminal uridylation and subsequent decay (10). Our finding that changes in mRNA levels on Pab1-AID depletion correlate with median pA tail length in WT cells (Fig. 2C-D), and more importantly that lengthening of pA tails on Pab1-AID depletion is suppressed by *dcp2Δ,* provide compelling support for this proposed mechanism. As yeast lacks terminal uridylation, the inability of short-tailed isoforms to bind Pab1 may lead to impaired eIF4F binding and increased accessibility of decapping enzyme at limiting Pab1 (Fig. S2B(ii)). This interpretation departs from a previous suggestion that increased lengths of bulk pA tails in the absence of Pab1 involves reduced rates of deadenylation (44). However, examining reporter mRNAs further suggested that, in addition to slower deadenylation, nascent mRNAs had longer pA tails and the long-tailed isoforms persisted longer in cells lacking Pab1 versus WT (19), in accordance with our conclusion. Although there is strong evidence that Pab1 recruits deadenylases to stimulate shortening of pA tails, there is also evidence that Pab1 blocks pA tail shortening beyond the size required to retain Pab1 (1). Under the conditions of our experiments, the latter function appears to predominate for most mRNAs to yield a net protection from deadenylation/degradation by Pab1 of short-tailed isoforms.

While having short median pA tails appears to be a determinant of accelerated mRNA decay on Pab1-AID depletion in otherwise WT cells, mRNAs with the longest pA tails are still destabilized on Pab1 depletion (Fig. 2C), implying that other features dictate preferential degradation at limiting Pab1. While it seems logical that inefficient assembly of the closed-loop would make transcripts hypersensitive to degradation at limiting Pab1 by exposing the cap to decapping enzyme, we did not find strong support for this prediction by examining the CDS lengths and association with Pab1, eIF4E/eIF4G (subunits of eIF4F) and inhibitory eIF4E-binding proteins in published RIP-Seq data (16) for the mRNAs down-regulated in abundance on Pab1 depletion (Table 1, col. 1). Thus, other sequence or structural features of mRNAs, or RBPs that recruit the degradation machinery, presumably render mRNAs more or less susceptible to decapping/decay at limiting Pab1.

Studies on PABP-depleted mammalian cells failed to uncover evidence implicating PABP in controlling the translational efficiencies of mRNAs (10,11). Bartel and colleagues attributed this finding partly to the observation that TEs are uncoupled from pA tail lengths in mammalian somatic cells, unlike in oocytes, presumably because PABP is not limiting in somatic cells and mRNAs with shorter pA tails are not at a competitive disadvantage for binding PABP. A second factor was the preferential degradation of mRNAs with shorter pA tails in somatic cells in the absence of PABP, eliminating them from the pool of transcripts from which TEs could be measured, whereas the mRNA decay machinery is present at low levels in oocytes (10). It has been concluded that budding yeast is also an uncoupled system, with no correlation between pA tail lengths and TE in WT cells (12). As noted above, we also found evidence in yeast for preferential degradation of transcripts with short pA tails. Nevertheless, our ribosome profiling data revealed a marked reprogramming of TEs on Pab1-AID depletion involving ∼1180 mRNAs who’s relative TEs were significantly decreased or increased by limiting Pab1. These changes in RPFs and TEs were highly correlated with changes in relative protein abundance determined by TMT-MS analysis, indicating that the alterations in translation initiation generally conferred corresponding changes in steady-state protein levels in Pab1-AID depleted cells.

Closed-loop assembly promotes translational efficiency by stabilizing eIF4F binding to the cap, which should enhance 43S PIC recruitment and scanning of the 5’UTR, and possibly recycling of ribosomes from the stop codon to the start codon on the same mRNAs (1). Thus, mRNAs inherently inefficient in closed-loop assembly might exhibit greater than average TE reductions at limiting Pab1. As summarized in Table 1 (col. 3), we found some support for this model, as the mRNAs showing TE reductions on Pab1-AID depletion in *DCP2* cells (TE_dn_*pab1*) tend to have longer than average CDSs and lower than average TEs in WT cells and are nearly devoid of the SCL mRNAs (Fig. S8A-C, green vs. grey). They also tend to be less stable, have low transcript abundance, and low codon optimality (Fig. S8D-F), properties of poorly translated mRNAs (55). Although they have greater than average pA tail lengths and eIF4E/eIF4G occupancies, they also have greater occupancies of the inhibitory 4E-BPs compared to all mRNAs (Fig. S8G-H). It is possible that other mRNA features or RBPs capable of repressing translation also contribute to rendering their TEs hypersensitive to limiting Pab1 when bulk mRNAs are reduced in *DCP2* cells.

A striking observation was that many of the TE changes, as well as the reduction in bulk polysome assembly evoked by Pab1-AID depletion, were suppressed by *dcp2Δ.* This result implies that the major effects of Pab1 depletion on both bulk translation and relative TEs of individual transcripts are indirect consequences of accelerated mRNA decapping/degradation and attendant global reductions in mRNA levels. One way to account for the marked dampening of TE changes would be to propose that reducing bulk mRNA levels by depleting Pab1 alters the mRNA binding distribution for one or more RBPs that interfere with assembly of the eIF4F-mRNP complex competent for translation initiation. Indeed, it was shown previously that translational stimulation by mammalian PABP in cell extracts was intensified by the presence of the RBP YB-1 and was attributed to the ability of PABP to overcome competition between YB-1 and eIF4G for association with the mRNA by preferentially occupying the pA tail and enabling cooperative binding of eIF4G near the mRNA cap (5). Our finding above that the TE_dn_*pab1* mRNAs generally have higher than average occupancies of inhibitory 4E-binding proteins Eap1 and Caf20 (Fig. S8H) is consistent with this explanation. Based on this, it is possible that the inefficiency of the TE_dn_*pab1* group of mRNAs in forming the closed-loop makes them particularly sensitive to competition with eIF4G for mRNA binding by an inhibitory RBP at their reduced mRNA levels on Pab1-AID depletion, leading to a lower proportion of transcripts that assemble eIF4F-mRNPs and, hence, to lower TEs. This hypothetical scenario is depicted for a 4E-BP in Fig. S9(i)-(ii) but could also apply to other RBPs that compete with eIF4G for mRNA binding. By restoring WT mRNA abundance, *dcp2*Δ would diminish the proposed competition, enabling eIF4G to occupy a greater fraction of these transcripts without the assistance of Pab1, and thereby restore their normal TEs (Fig. S9(iii)). An important step in testing this model would be to identify one or more RBPs required for the TE reductions conferred by Pab1-AID depletion in *DCP2* cells.

More than 600 mRNAs showed an increase in relative TE on Pab1 depletion in *DCP2* cells (TE_up_*pab1,* Fig. S6D). This behavior might be explained if these mRNAs have a greater than average propensity to form the closed-loop, making them more resistant to the competition with eIF4G by RBPs envisioned above (Fig. S9). However, this possibility is not supported by their properties (summarized in Table 1, col. 4), as the TE_up_*pab1* transcripts tend to have much longer than average CDS lengths and low TEs in WT (Fig. S8A-B), low occupancies of Pab1 and eIF4E and greater than average occupancies of Eap1 (Fig. S10A); and they are underrepresented among the SCL transcripts (Fig. S10B). On the other hand, these mRNAs tend to be more stable, more abundant, and to have higher-than-average codon optimalities (Fig. S8D-F). They are further distinctive in not exhibiting reduced absolute mRNA levels on Pab1-AID depletion in *DCP2* cells (Fig. 5F, col. 6 vs. 2). This increased abundance relative to other mRNAs might allow the TE_up_*pab1* transcripts to escape the proposed increased competition with eIF4G exerted by RBPs (Fig. S9(ii)) and achieve a relative increase in TE compared to other mRNAs whose abundance declines at limiting Pab1. The increase in relative mRNA abundance of the TE_up_group might also allow them to compete better for limiting PICs on Pab1-AID depletion compared to WT cells. In fact, we observed a reduction in total ribosome abundance in the *pab1-AID* mutant (Fig. S11A-B), which likely results from the strong down-regulation of rESR transcripts (Fig. 1E-F) encoding ribosomal proteins and ribosome biogenesis factors (45). We have shown previously that changes in 40S ribosome:mRNA ratios can be instrumental in translational reprogramming in yeast (48,56). Thus, even though the TE_up_*pab1* mRNAs will remain inefficiently translated on Pab1-AID depletion, they will exhibit increased relative TEs because their resistance to enhanced decay makes them less sensitive to inhibitory RBPs and reduced levels of 43S PICs compared to other mRNAs. As this advantage will be nullified by *dcp2*Δ with the restoration of normal transcript levels, this proposed explanation also accounts for the dampening of TE increases conferred by *dcp2*Δ for the TE_up_*pab1* group (Figs. 6C-D). The postulated alternative fates of the TE_dn_*pab1* and TE_up_*pab1* mRNAs on Pab1-AID depletion in the presence or absence of *DCP2* are summarized schematically in Fig. S12.

While the great majority of TE changes conferred by Pab1-AID depletion were dampened or eliminated by *dcp2Δ,* a small group of 116 mRNAs still showed significant TE reductions in *dcp2Δ pab1-AID* versus *dcp2*Δ cells (the TE_dn_*dp* group), implying a more direct role for Pab1 in stimulating their translation. The fact that ∼60% of these mRNAs showed TE reductions only in *dcp2*Δ cells (GroupIII in Fig. 6F) suggests that Pab1’s direct contribution to their translation was obscured in *DCP2* cells by changes in PIC:mRNA or RBP:mRNA ratios conferred by the bulk mRNA reductions produced by Pab1-AID depletion in the presence of Dcp2. As summarized in Table 1 (cf. cols. 5 & 3), the entire group of 116 TE_dn_*dp* mRNAs have properties more consistent with closed-loop assembly compared to the larger group of 513 mRNAs translationally down-regulated in *DCP2* cells (TE_dn_*pab1*), exhibiting greater Pab1 but lower Caf20 occupancies, shorter CDS lengths and higher TEs in WT (Fig. S13A-C). Both groups however have relatively low transcript abundance, low mRNA half-lives and codon-optimality (Fig. S13D-F, Table 1, cols. 5 & 3) and are depleted of SCL mRNAs (Fig. S8C & S13G). Perhaps the 116 TE_dn_*dp* mRNAs achieve their average TEs in WT cells in a manner strongly dependent on closed-loop assembly and are out-competed by other mRNAs for binding the low-level Pab1 remaining after Pab1-AID depletion. This competition might be relatively more intense for the GroupIII vs. GroupII subset of these transcripts (Fig. 6F) and increased further by deleting *DCP2* and restoring bulk mRNA levels, possibly accounting for GroupIII’s strong TE reductions on Pab1-AID depletion only in *dcp2*Δ cells (Fig. 6F, cols. 5-6). The dampening of TE reductions conferred by *dcp2*Δ for the GroupII subset (Fig. 6G, GroupII) could follow the hypothetical mechanism in Fig. S9 involving inhibitory RBPs. Further experiments will be required to test these proposals.

The final important observation of this study is that transcripts preferentially targeted for degradation on Pab1-AID depletion include the mRNAs encoding all four canonical histones, leading to reduced expression of the histone proteins. Low-level histone expression is known to activate cryptic promoters located within CDSs, which normally have high nucleosome densities compared to 5’ and 3’ non-coding regions of genes. The closely packed genic nucleosomes normally impede assembly of functional transcription initiation complexes within CDSs. As such, cryptic internal promoters become activated in mutants with abnormally low genic nucleosome densities, which include those lacking a full complement of histone genes or deficient for a histone chaperone, like Spt6, required to assemble genic nucleosomes and maintain their WT densities (51). Interestingly, our CAGE analysis of capped mRNA 5’ ends revealed that depletion of Pab1-AID confers a genome-wide increase in the number of transcription start site clusters (TCs) driven largely by TCs mapping within gene CDSs. The majority of TCs derepressed by Pab1-AID were found to be derepressed in a *spt6-1004* mutant and were also enriched for those up-regulated by eliminating histone H3 methyltransferases Set1 and Set2, which mediate co-transcriptional suppression of internal cryptic promoters directed by transcription from upstream canonical promoters (51). These findings suggest that Pab1-AID depletion reduces genic nucleosome densities sufficiently to derepress internal cryptic promoters.

In summary, our results provide valuable new insights into the molecular functions of Pab1 in cells, showing that it protects most mRNAs from decapping by Dcp1:Dcp2 and subsequent decay in a manner accentuated for transcripts/isoforms with short pA tails but not those with a low propensity to form the closed-loop intermediate. Despite the uncoupling of pA tail lengths and TEs described for yeast, we found that depleting Pab1 conferred extensive reprogramming of TEs, but in a manner heavily influenced by the decreased mRNA levels produced at limiting Pab1 through enhanced decapping. The fact that only ∼120 mRNAs displayed TE reductions on Pab1-AID depletion in *dcp2*Δ cells, where the decline in mRNA levels was suppressed, implies that only a small fraction of mRNAs rely heavily on closed-loop formation via Pab1-eIF4G interaction in WT cells—at least under the optimum growth conditions of our experiments. Finally, histone mRNAs are preferentially targeted for decapping/decay in cells depleted of Pab1, leading to reduced histone protein expression and the derepression of internal cryptic promoters expected from reduced genic nucleosome densities, identifying a new layer of post-transcriptional control over histone homeostasis involving Pab1.

## DATA AVAILABILITY

All primary data obtained from RNA-Seq, Ribo-Seq, SM-PAT-Seq, CAGE sequencing, and TMT-MS are deposited at the Gene Expression Omnibus data repository under pending accession numbers.

## ACKNOWLEDGEMENTS

We thank members of our laboratories and those of the Dever, Lorsch, and Guydosh groups for helpful comments and suggestions. We thank Fabio Rueda Faucz of the NICHD Molecular Genomics core for advice and assistance, and Fred Winston and Maurice Swanson for generous gifts of yeast strains and Pab1 antibodies. We thank Dr. Anil Thakur with his help in generating Pab1-AID strains. This work utilized the computational resources of the NIH HPC Biowulf cluster. (http://hpc.nih.gov). It was supported in part by the Intramural Research Program of the National Institutes of Health and of X.N. and Z.L. by NSF grant 1951332 and the Saint Louis University 2022 President’s Research Fund.

## Notes

### Competing Interest Statement

The authors have declared no competing interest.

## REFERENCES

1. Passmore, L.A. and Coller, J. (2022) Roles of mRNA poly(A) tails in regulation of eukaryotic gene expression. Nat Rev Mol Cell Biol, 23, 93–106.

2. Tarun, S.Z. and Sachs, A.B. (1995) A common function for mRNA 5’ and 3’ ends in translation initiation in yeast. Genes & Dev, 9, 2997–3007.

3. Tarun, S.Z., Wells, S.E., Deardorff, J.A. and Sachs, A.B. (1997) Translation initiation factor eIF4G mediates in vitro poly (A) tail-dependent translation. Proceedings of the National Academy of Sciences of the United States of America, 94, 9046–9051.

4. Kahvejian, A., Svitkin, Y.V., Sukarieh, R., M’Boutchou, M.N. and Sonenberg, N. (2005) Mammalian poly(A)-binding protein is a eukaryotic translation initiation factor, which acts via multiple mechanisms. Genes Dev, 19, 104–113.

5. Svitkin, Y.V., Evdokimova, V.M., Brasey, A., Pestova, T.V., Fantus, D., Yanagiya, A., Imataka, H., Skabkin, M.A., Ovchinnikov, L.P., Merrick, W.C. et al. (2009) General RNA-binding proteins have a function in poly(A)-binding protein-dependent translation. Embo J, 28, 58–68.

6. Hinnebusch, A.G. (2011) Molecular mechanism of scanning and start codon selection in eukaryotes. Microbiol Mol Biol Rev, 75, 434–467.

7. Gulay, S., Gupta, N., Lorsch, J.R. and Hinnebusch, A.G. (2020) Distinct interactions of eIF4A and eIF4E with RNA helicase Ded1 stimulate translation in vivo. Elife, 9.

8. Cosson, B., Couturier, A., Chabelskaya, S., Kiktev, D., Inge-Vechtomov, S., Philippe, M. and Zhouravleva, G. (2002) Poly(A)-binding protein acts in translation termination via eukaryotic release factor 3 interaction and does not influence [PSI(+)] propagation. Mol Cell Biol, 22, 3301–3315.

9. Uchida, N., Hoshino, S., Imataka, H., Sonenberg, N. and Katada, T. (2002) A Novel Role of the Mammalian GSPT/eRF3 Associating with Poly(A)-binding Protein in Cap/Poly(A)-dependent Translation. J Biol Chem, 277, 50286–50292.

10. Xiang, K. and Bartel, D.P. (2021) The molecular basis of coupling between poly(A)-tail length and translational efficiency. Elife, 10.

11. Kajjo, S., Sharma, S., Chen, S., Brothers, W.R., Cott, M., Hasaj, B., Jovanovic, P., Larsson, O. and Fabian, M.R. (2022) PABP prevents the untimely decay of select mRNA populations in human cells. EMBO J, 41, e108650.

12. Subtelny, A.O., Eichhorn, S.W., Chen, G.R., Sive, H. and Bartel, D.P. (2014) Poly(A)-tail profiling reveals an embryonic switch in translational control. Nature, 508, 66–71.

13. Eichhorn, S.W., Subtelny, A.O., Kronja, I., Kwasnieski, J.C., Orr-Weaver, T.L. and Bartel, D.P. (2016) mRNA poly(A)-tail changes specified by deadenylation broadly reshape translation in Drosophila oocytes and early embryos. Elife, 5.

14. Park, J.E., Yi, H., Kim, Y., Chang, H. and Kim, V.N. (2016) Regulation of Poly(A) Tail and Translation during the Somatic Cell Cycle. Mol Cell, 62, 462–471.

15. Park, E., Walker, S., Lee, J., Rothenburg, S., Lorsch, J. and Hinnebusch, A. (2011) Multiple elements in the eIF4G1 N-terminus promote assembly of eIF4G1 PABP mRNPs in vivo. EMBO J, 30, 302–316.

16. Costello, J., Castelli, L.M., Rowe, W., Kershaw, C.J., Talavera, D., Mohammad-Qureshi, S.S., Sims, P.F., Grant, C.M., Pavitt, G.D., Hubbard, S.J. et al. (2015) Global mRNA selection mechanisms for translation initiation. Genome Biol, 16, 10.

17. Muhlrad, D., Decker, C.J. and Parker, R. (1994) Deadenylation of the unstable mRNA encoded by the yeast MFA2 gene leads to decapping followed by 5’-->3’ digestion of the transcript. Genes Dev, 8, 855–866.

18. Decker, C.J. and Parker, R. (1993) A turnover pathway for both stable and unstable mRNAs in yeast: evidence for a requirement for deadenylation. Genes Dev, 7, 1632–1643.

19. Caponigro, G. and Parker, R. (1995) Multiple functions for the poly(A)-binding protein in mRNA decapping and deadenylation in yeast. Genes Dev, 9, 2421–2432.

20. Sadowski, I., Su, T.C. and Parent, J. (2007) Disintegrator vectors for single-copy yeast chromosomal integration. Yeast, 24, 447–455.

21. Qiu, H., Hu, C., Wong, C.M. and Hinnebusch, A.G. (2006) The Spt4p subunit of yeast DSIF stimulates association of the Paf1 complex with elongating RNA polymerase II. Mol Cell Biol, 26, 3135–3148.

22. Alting-Mees, M.A. and Short, J.M. (1989) pBluescript II: gene mapping vectors. Nucleic Acids Res, 17, 9494.

23. 23. Zeidan, Q., He, F., Zhang, F., Zhang, H., Jacobson, A. and Hinnebusch, A.G. (2018) Conserved mRNA-granule component Scd6 targets Dhh1 to repress translation initiation and activates Dcp2-mediated mRNA decay in vivo. PLoS Genet, 14, e1007806.

24. Young, D.J. and Guydosh, N.R. (2019) Hcr1/eIF3j Is a 60S Ribosomal Subunit Recycling Accessory Factor In Vivo. Cell Rep, 28, 39–50 e34.

25. McGlincy, N.J. and Ingolia, N.T. (2017) Transcriptome-wide measurement of translation by ribosome profiling. Methods, 126, 112–129.

26. Kim, D., Paggi, J.M., Park, C., Bennett, C. and Salzberg, S.L. (2019) Graph-based genome alignment and genotyping with HISAT2 and HISAT-genotype. Nat Biotechnol, 37, 907–915.

27. Love, M.I., Huber, W. and Anders, S. (2014) Moderated estimation of fold change and dispersion for RNA-seq data with DESeq2. Genome Biol, 15, 550.

28. Langmead, B. and Salzberg, S.L. (2012) Fast gapped-read alignment with Bowtie 2. Nat Methods, 9, 357–359.

29. Langmead, B., Trapnell, C., Pop, M. and Salzberg, S.L. (2009) Ultrafast and memory-efficient alignment of short DNA sequences to the human genome. Genome Biol, 10, R25.

30. Iben, J.R., Li, T., Mattijssen, S. and Maraia, R.J. (2024) Single-Molecule Poly(A) Tail Sequencing (SM-PATseq) Using the PacBio Platform. Methods Mol Biol, 2723, 285–301.

31. Schmitt, M.E., Brown, T.A. and Trumpower, B.L. (1990) A rapid and simple method for preparation of RNA from Saccharomyces cerevisiae. Nucleic Acids Res, 18, 3091–3092.

32. Murata, M., Nishiyori-Sueki, H., Kojima-Ishiyama, M., Carninci, P., Hayashizaki, Y. and Itoh, M. (2014) Detecting expressed genes using CAGE. Methods Mol Biol, 1164, 67–85.

33. Lu, Z., Berry, K., Hu, Z., Zhan, Y., Ahn, T.H. and Lin, Z. (2021) TSSr: an R package for comprehensive analyses of TSS sequencing data. NAR Genom Bioinform, 3, lqab108.

34. Lu, Z. and Lin, Z. (2021) The origin and evolution of a distinct mechanism of transcription initiation in yeasts. Genome research, 31, 51–63.

35. Zecha, J., Satpathy, S., Kanashova, T., Avanessian, S.C., Kane, M.H., Clauser, K.R., Mertins, P., Carr, S.A. and Kuster, B. (2019) TMT Labeling for the Masses: A Robust and Cost-efficient, In-solution Labeling Approach. Mol Cell Proteomics, 18, 1468–1478.

36. Wisniewski, J.R., Zougman, A., Nagaraj, N. and Mann, M. (2009) Universal sample preparation method for proteome analysis. Nat Methods, 6, 359–362.

37. Nesvizhskii, A.I., Keller, A., Kolker, E. and Aebersold, R. (2003) A statistical model for identifying proteins by tandem mass spectrometry. Anal Chem, 75, 4646–4658.

38. 38. Huber, W., von Heydebreck, A., Sultmann, H., Poustka, A. and Vingron, M. (2002) Variance stabilization applied to microarray data calibration and to the quantification of differential expression. Bioinformatics, 18 **Suppl 1**, S96–104.

39. Ritchie, M.E., Phipson, B., Wu, D., Hu, Y., Law, C.W., Shi, W. and Smyth, G.K. (2015) limma powers differential expression analyses for RNA-sequencing and microarray studies. Nucleic Acids Res, 43, e47.

40. Reid, G.A. and Schatz, G. (1982) Import of proteins into mitochondria. Yeast cells grown in the presence of carbonyl cyanide m-chlorophenylhydrazone accumulate massive amounts of some mitochondrial precursor polypeptides. J Biol Chem, 257, 13056–13061.

41. Nanda, J.S., Cheung, Y.N., Takacs, J.E., Martin-Marcos, P., Saini, A.K., Hinnebusch, A.G. and Lorsch, J.R. (2009) eIF1 controls multiple steps in start codon recognition during eukaryotic translation initiation. J Mol Biol, 394, 268–285.

42. Olsen, D.S., E.M., S., Mathew, A., Zhang, F., Krishnamoorthy, T., Phan, L. and Hinnebusch, A.G. (2003) Domains of eIF1A that mediate binding to eIF2, eIF3 and eIF5B and promote ternary complex recruitment in vivo. EMBO J, 22, 193–204.

43. Robinson, J.T., Thorvaldsdottir, H., Winckler, W., Guttman, M., Lander, E.S., Getz, G. and Mesirov, J.P. (2011) Integrative genomics viewer. Nat Biotechnol, 29, 24–26.

44. Sachs, A.B. and Davis, R.W. (1989) The Poly(A) binding protein is requried for Poly(A) shortening and 60S ribosomal subunit-dependent translation initiation. Cell, 58, 857–867.

45. Gasch, A.P., Spellman, P.T., Kao, C.M., Carmel-Harel, O., Eisen, M.B., Storz, G., Botstein, D. and Brown, P.O. (2000) Genomic expression programs in the response of yeast cells to environmental changes. Mol Biol Cell, 11, 4241–4257.

46. Coon, S.L., Li, T., Iben, J.R., Mattijssen, S. and Maraia, R.J. (2021) Single-molecule polyadenylated tail sequencing (SM-PAT-Seq) to measure polyA tail lengths transcriptome-wide. Methods Enzymol, 655, 119–137.

47. Brambilla, M., Martani, F., Bertacchi, S., Vitangeli, I. and Branduardi, P. (2019) The Saccharomyces cerevisiae poly (A) binding protein (Pab1): Master regulator of mRNA metabolism and cell physiology. Yeast, 36, 23–34.

48. Gaikwad, S., Ghobakhlou, F., Young, D.J., Visweswaraiah, J., Zhang, H. and Hinnebusch, A.G. (2021) Reprogramming of translation in yeast cells impaired for ribosome recycling favors short, efficiently translated mRNAs. Elife, 10.

49. Amrani, N., Ghosh, S., Mangus, D.A. and Jacobson, A. (2008) Translation factors promote the formation of two states of the closed-loop mRNP. Nature, 453, 1276–1280.

50. Cheung, V., Chua, G., Batada, N.N., Landry, C.R., Michnick, S.W., Hughes, T.R. and Winston, F. (2008) Chromatin- and transcription-related factors repress transcription from within coding regions throughout the Saccharomyces cerevisiae genome. PLoS Biol, 6, e277.

51. Rando, O.J. and Winston, F. (2012) Chromatin and transcription in yeast. Genetics, 190, 351–387.

52. Lu, Z. and Lin, Z. (2019) Pervasive and dynamic transcription initiation in Saccharomyces cerevisiae. Genome Res, 29, 1198–1210.

53. Smolle, M. and Workman, J.L. (2013) Transcription-associated histone modifications and cryptic transcription. Biochim Biophys Acta, 1829, 84–97.

54. 54. O’Duibhir, E., Lijnzaad, P., Benschop, J.J., Lenstra, T.L., van Leenen, D., Groot Koerkamp, M.J., Margaritis, T., Brok, M.O., Kemmeren, P. and Holstege, F.C. (2014) Cell cycle population effects in perturbation studies. Mol Syst Biol, 10, 732.

55. Weinberg, D.E., Shah, P., Eichhorn, S.W., Hussmann, J.A., Plotkin, J.B. and Bartel, D.P. (2016) Improved Ribosome-Footprint and mRNA Measurements Provide Insights into Dynamics and Regulation of Yeast Translation. Cell Rep, 14, 1787–1799.

56. Vijjamarri, A.K., Niu, X., Vandermeulen, M.D., Onu, C., Zhang, F., Qiu, H., Gupta, N., Gaikwad, S., Greenberg, M.L., Cullen, P.J. et al. (2023) Decapping factor Dcp2 controls mRNA abundance and translation to adjust metabolism and filamentation to nutrient availability. Elife, 12.

57. Chan, L.Y., Mugler, C.F., Heinrich, S., Vallotton, P. and Weis, K. (2018) Non-invasive measurement of mRNA decay reveals translation initiation as the major determinant of mRNA stability. Elife, 7.

58. Nishimura, K., Fukagawa, T., Takisawa, H., Kakimoto, T. and Kanemaki, M. (2009) An auxin-based degron system for the rapid depletion of proteins in nonplant cells. Nat Methods, 6, 917–922.

59. Pelechano, V., Wei, W. and Steinmetz, L.M. (2013) Extensive transcriptional heterogeneity revealed by isoform profiling. Nature, 497, 127–131.

60. Neymotin, B., Athanasiadou, R. and Gresham, D. (2014) Determination of in vivo RNA kinetics using RATE-seq. RNA, 20, 1645–1652.

61. Lahtvee, P.J., Sanchez, B.J., Smialowska, A., Kasvandik, S., Elsemman, I.E., Gatto, F. and Nielsen, J. (2017) Absolute Quantification of Protein and mRNA Abundances Demonstrate Variability in Gene-Specific Translation Efficiency in Yeast. Cell Syst, 4, 495–504 e495.

62. He, F., Li, X., Spatrick, P., Casillo, R., Dong, S. and Jacobson, A. (2003) Genome-wide analysis of mRNAs regulated by the nonsense-mediated and 5’ to 3’ mRNA decay pathways in yeast. Mol Cell, 12, 1439–1452.

63. Ginsburg, D.S., Govind, C.K. and Hinnebusch, A.G. (2009) The NuA4 lysine acetyltransferase Esa1 is targeted to coding regions and stimulates transcription elongation with Gcn5. Mol Cell Biol.

64. Winzeler, E.A., Shoemaker, D.D., Astromoff, A., Liang, H., Anderson, K., Andre, B., Bangham, R., Benito, R., Boeke, J.D., Bussey, H. et al. (1999) Functional Characterization of the *S. cerevisiae* Genome by Gene deletion and Parallel Analysis. Science, 285, 901–906.

65. Sikorski, R.S. and Hieter, P. (1989) A system of shuttle vectors and yeast host strains designed for efficient manipulation of DNA in *Saccharomyces cerevisiae*. Genetics, 122, 19–27.

66. Morawska, M. and Ulrich, H.D. (2013) An expanded tool kit for the auxin-inducible degron system in budding yeast. Yeast, 30, 341–351.

