## Supplementary Material for "Yeast poly(A)-binding protein (Pab1) controls translation initiation in vivo primarily by blocking mRNA decapping and decay"

### SUPPLEMENTARY FIGURE LEGENDS

**Fig. S1. Pab1 depletion in an auxin-inducible degron mutant confers widespread changes in mRNA abundance. (A)** Schematic representation of auxin-induced degradation of AID-tagged Pab1 adapted from (58). The chromosomal *PAB1* gene is tagged by inserting sequences encoding the 71-114 amino-acids of an auxin-inducible degron and 9 tandem Myc epitopes ($AID^{71\text{-}114}$-9myc) immediately before the stop codon in a strain harboring the *OsTIR* gene, encoding rice *Oryza sativa* F-box protein Tir1 expressed from the yeast *ADH1* promoter, inserted at the *LYS2* locus. In auxin-treated cells the $SCF^{TIR}$ E3 ubiquitin ligase comprised of OsTir, Skp1, Rbx1, Cdc53, an E2 ubiquitin ligase and ubiquitin attaches a ubiquitin chain to Pab1-AID $AID^{71\text{-}114}$-9myc, targeting it for proteasomal degradation. **(B)** Serial 10-fold dilutions of WT strain W303 and strains containing *OsTIR* integrated at *LYS2* containing WT *PAB1* (PMY1) or *pab1-AID* (PMY2) were spotted on YPD agar supplemented with either 1mM of KOH or 1mM of synthetic auxin 1-naphthalene acetic acid (NAA) and incubated at 30°C. **(C)** Western blot analysis of strains PMY1 (WT) and PMY2 (*pab1-AID*) cultured in YPD medium to log phase and treated with 1 mM NAA for 6 hr (+) or 1mM KOH (-) as indicated at the top. Whole cell extracts prepared under denaturing conditions were resolved by 4-20% TGX SDS-PAGE, blotted to a PVDF membrane, and probed with antibodies against myc (detects both Pab1-AID-9myc and OsTIR-7myc), Pab1, or Hcr1 (examined as a loading control), as indicated on the right along with positions of molecular weight markers resolved on the same gel. Below the blot are listed the $\log_2$ fold-change in Pab1-AID vs Pab1 relative protein abundance ($\log_2\Delta$protein *pab1*/WT) and corresponding padj. value for the change determined by TMT-MS analysis of NAA-treated PMY2 (*pab1-AID*) vs NAA-treated PMY1 (WT) cells, as described below in RESULTS. (**D)** Volcano-plot of $\log_2\Delta$mRNA values determined for NAA-treated vs. untreated PMY2 (*pab1-AID*) plotted against the negative $\log_{10}$ adjusted p-values (padj.) for the fold-changes determined by DESeq2 analysis of the spike-in normalized RNA-Seq data (y-axis). The dotted line marks the 5% padj. threshold for mRNAs showing a significant increase (n=367) or decrease (n=3195) in mRNA abundance at padj. $< 0.05$, shown in purple and green dots, respectively. Outlier mRNAs (2-67) with -log10 padj. >150 and $\log_2\Delta$mRNA values >10 were excluded to expand the y and x-axis scales. **(E)** Proportional Venn diagrams of overlaps between the mRNA_up and mRNA_dn groups for *pab1*_aux/WT_aux (from Fig.1C) and *pab1*_aux/*pab1* (from panel D). Hypergeometric distribution p-values are shown for overlaps. **(F)** Boxplot of $\log_2\Delta$mRNA values of all expressed mRNAs (1-2) and the mRNA_dn_*pab1* (3-4) and mRNA_up_*pab1* (5-6) groups defined in panel D, for the WT_aux/WT or *pab1*_aux/*pab1* comparisons as indicated on the x-axis, removing 0-57 outlier mRNAs with values >6.0 or <-6.0. **(G)** Correlation analysis of RNA-Seq results for biological replicates. Correlation matrix showing Spearman correlation coefficients calculated for pair-wise comparisons of numbers of RPKM-normalized RNA-Seq reads for all expressed genes among all 18 RNA-Seq libraries generated for 3 biological replicates (rep1-rep3) of untreated or NAA-treated *OsTIR* strains PMY1 (WT), PMY2 (*pab1AID*), PMY3 (*dcp2Δ,*) and PRY1 (*dcp2Δpab1AID*). The correlation coefficients for all comparisons are between 1.0-0.88 with a P-value ≈ 0. For this and all similar plots below, the eccentricity of the ellipses are scaled parametrically to the correlation value between the two samples.

**Fig. S2. Depletion of Pab1-AID leads to pA tail shortening driven partly by preferential degradation of short-tailed transcripts. (A)** Reproducibility of pA tail length distributions and median values among biological replicates. Boxplot of pA tail lengths (in nt) calculated for each gene in the three biological replicates (rep1-rep3) and average of all three replicates (avg) obtained for NAA-treated (aux) WT, *pab1-AID*, *dcp2Δ* and *dcp2Δpab1-AID* cells. Outliers (0-4) with median pA tail lengths >150 nt were removed to expand the y-axis. **(B)** Schematic model to explain preferential decapping/decay of short-tailed transcript isoforms at limiting Pab1. In WT (*top*), ample levels of Pab1 allow its binding to isoforms with both long (i) or short (ii) pA tails and closed-loop assembly via Pab1-eIF4G interaction stabilizes cap-binding by eIF4E and blocks access of Dcp2:Dcp1 to prevent decapping. Limiting Pab1 in *pab1-AID* cells (*bottom*) preferentially reduces Pab1 binding to short-tailed isoforms, eliminating closed-loop assembly, and increasing decapping/decay. **(C)** Boxplot of median pA tail lengths (in nt) calculated for each of 5476 yeast genes and for mRNA_dn_*pab1* and mRNA_up_*pab1* groups identified by overlaps in Fig S1E, in NAA-treated WT or treated *pab1-AID* cells. Three to four outliers with median pA tail lengths >100 nt were removed to expand the y-axis.

**Fig. S3. Removing *DCP2* suppresses reductions in mRNA abundance in Pab1-depleted cells. (A)** Serial 10-fold dilutions of strains of the indicated relevant genotypes were spotted on YPD agar supplemented with either 1mM KOH or 1mM NAA and incubated at 30°C: (1) W303, (2) PMY1, (3) PMY2, (4) CFY1016, (5) CFY1016 containing *DCP2* on plasmid pQZ145, (6) PMY3, (7) PMY3 containing *DCP2* on *URA3* plasmid pRS316, (8) PRY1, and (9) PRY1 containing pQZ145. Results in rows 1-3 are identical to those in Fig. S1B and depicted here only for comparison. **(B)** Results of Western blot (*upper*) and TMT-MS (*lower*) analyses conducted exactly as in Fig. S1C for *OsTIR* strains PMY1 (*PAB1*, 1-2), PMY2 (*pab1-AID*, 3-4), PMY3 (*dcp2Δ*, 5-8) and PRY1 (*dcp2Δ pab1-AID*, 9-12) cultured in YPD medium to log phase and treated with 1 mM NAA for 6 hr (+) or 1mM KOH (-) as indicated. **(C)** Proportional Venn diagrams showing overlaps between the 3008 mRNA_dn_*pab1* mRNAs from Fig. S1E and the 92 mRNA_dn_ *dp* mRNAs from Fig3C(i). Hypergeometric distribution p-value is shown for the overlap yielding transcripts designated as GroupII. **(D)** Boxplot of spike-in normalized $\log_2\Delta$mRNA values for the mRNA_dn_*pab1* and three mRNA_dn_*dp* groups defined in (D) in NAA-treated *pab1-AID* vs. treated WT cells (1, 3, 5 and 7) and NAA-treated *dcp2Δpab1-AID* vs. treated *dcp2Δ* cells (2, 4, 6 and 8). Outlier mRNAs (0-24) with log2ΔmRNA >2.0 or < -4.0 were removed.

**Fig. S4. mRNAs down-regulated at limiting Pab1-AID levels do not conform to strong-closed loop (SCL) mRNAs. (A)** Boxplot showing log2 fold-changes in IP/Total RNA immunoprecipitated with TAP-tagged versions of the indicated proteins in the RIP-Seq data of Costello et al. (2015) for all expressed mRNAs and the mRNA_dn_*pab1* mRNAs defined by the overlaps in Fig. S1E. Outlier mRNAs (0-28) with log2ΔFC values of >4.0 or <-4.0 were removed. **(B)** Boxplot showing coding sequence (CDS) lengths in nucleotides (from (59)) of all expressed mRNAs, mRNA_dn_*pab1* (this study), mRNA_up_*pab1* (this study) and strong closed loop (SCL) mRNAs (16). Outliers (0-19) with CDS lengths >10000 and <100 nts were removed. **(C)** Boxplot showing log2 translational efficiency (TE) in WT cells strain (this study) of all expressed mRNAs, mRNA_dn_*pab1*, mRNA_up_*pab1* and the SCL mRNAs. Outliers (0-79) with log2TE values >6.0 and <-6.0 were removed. **(D)** Proportional Venn diagrams showing overlaps between the 3008 mRNA_dn_*pab1* mRNAs and SCL mRNAs. Hypergeometric distribution p-value is shown for the overlap. **(E)** Boxplot showing log2 WT mRNA half-life in min (from (60)) for all expressed mRNAs and mRNA_dn_*pab1* and mRNA_up_*pab1* mRNAs. **(F)** Boxplot showing log2 5'UTR lengths in nucleotides (from Pelechano et al. (2013)) for all expressed mRNAs and mRNA_dn_*pab1* and mRNA_up_*pab1* mRNAs. **(G)** Boxplot showing log2 3'UTR lengths in nucleotides (from Pelechano et al. (2013) for all expressed mRNAs and mRNA_dn_*pab1* and mRNA_up_*pab1* mRNAs. Outlier mRNAs (0-5) with log2 lengths <2.0 were removed. **(H)** Boxplot showing stAI values (data from Radhakrishnan et al., 2016) for all expressed mRNAs and mRNA_dn_*pab1* and mRNA_up_*pab1* mRNAs. **(I)** Boxplot showing log2 steady state transcript abundance in molecules per dry cellular weight (pgDW) (from (61)) for all expressed mRNAs and mRNA_dn_*pab1* and mRNA_up_*pab1* mRNAs. Outlier mRNAs (0-14) with log2 transcript abundance >6.0 and <-6.0 were removed.

**Fig. S5. Pab1 depletion leads to derepressed internal cryptic promoters. (A)** Correlation analysis of raw counts of all TSSs between biological replicates by CAGE libraries. Correlation matrix showing Pearson correlation coefficients calculated for pair-wise comparisons of raw counts of all TSSs between all 14 CAGE libraries generated for 2 biological replicates (rep1-rep2) or for 3 biological replicates (rep1-rep3) respectively of untreated *spt6-1006*, isogenic *SPT6* WT, *set1Δset2Δ*, isogenic *SET1SET2* WT, NAA-treated *OsTIR* strains PMY1 (WT) and PMY2 (*pab1AID*) cells. The correlation coefficients for all comparisons are between 1.0-0.49 with a P-value ≈ 0. **(B-E)** Screenshot of the IGV Genome Browser showing CAGE signals (TPM) for plus and minus strands for genes **(A)** *CSG2*, **(B)** *TCD1*, **(C)** *NRP1* and **(D)** *BAT2* in NAA-treated *pab1-AID* or WT *PAB1* cells and untreated *spt6-1004* and WT *SPT6* cells. All rows are scaled equally with the ranges in brackets for the plus and minus strands.

**Fig. S6. Pab1 depletion down-regulates bulk translation and reprograms translational efficiencies of many genes. (A)** Correlation analysis of RPFs determined for biological replicates by Ribo-Seq.

Correlation matrix showing Spearman correlation coefficients calculated for pair-wise comparisons of numbers of RPKM-normalized RPF reads for all expressed genes among all 12 Ribo-Seq libraries generated for 3 biological replicates (rep1-rep3) of untreated or NAA-treated *OsTIR* strains PMY1 (WT) and PMY2 (*pab1AID*). The correlation coefficients for all comparisons are between 1.0-0.92 with a P-value ≈ 0. **(B)** Volcano-plot showing $\log_2$ fold-changes in relative translation efficiencies (ΔTE) for all 5816 yeast mRNAs in NAA-treated vs. untreated *pab1-AID* cells vs. the $-\log_{10}$ padj. values for ΔTE changes determined by DESeq2 analysis of the profiling data (y-axis). The dotted line marks the 10% padj. threshold for mRNAs showing a significant increase (n=777, >1.4-fold) or decrease (n=700, <1.4-fold) in ΔTE, plotted in purple and green, respectively. 7 outlier mRNAs with $-\log_{10}$ padj. >50 were excluded. **(C)** Proportional Venn diagrams of overlaps between the TE_up and TE_dn groups for *pab1*_aux/WT_aux (from Fig.5B) and for *pab1*_aux/*pab* (from panel B). Hypergeometric distribution p-values are shown for overlaps. **(D)** Boxplot of $\log_2$ΔTE values for NAA-treated vs. untreated *pab1-AID* cells for all expressed mRNAs or the TE_up_*pab1* and TE_dn_*pab1* groups defined by the overlaps in panel C, removing 4-15, outlier mRNAs with $\log_2$ΔTE values >4.0 or <-4.0. **(E)** Correlation analysis of normalized peptide abundances between biological replicates determined by TMT-MS analysis. Correlation matrix showing Spearman correlation coefficients calculated for pair-wise comparisons of log2 Cyclic Loess Normalized Exclusive MS1 intensities for all expressed proteins among 6 TMT-MS analyses conducted for 3 biological replicates (rep1-rep3) of NAA-treated *OsTIR* strains PMY1 (WT) and PMY2 (*pab1AID*). The correlation coefficients for all comparisons are >0.98 with a P-value ≈ 0.

**Fig. S7. Eliminating *DCP2* suppresses reductions in bulk translation and TE changes in Pab1-depleted cells. (A)** Correlation analysis of RPFs determined for biological replicates by Ribo-Seq. Correlation matrix showing Spearman correlation coefficients calculated for pair-wise comparisons of numbers of RPKM-normalized RPF reads for all expressed genes among 6 Ribo-Seq libraries generated for 3 biological replicates (rep1-rep3) of NAA-treated PMY3 (*dcp2Δ*,) and PRY1 (*dcp2Δpab1AID*). The correlation coefficients for all comparisons are between 1.0-0.98 with a P-value ≈ 0. **(B)** Boxplot of spike-in normalized $log_2$ΔmRNA values for the TE_dn_*pab1*, TE_up_*pab1* and three mRNA groups defined in Fig. 6F in NAA-treated *pab1-AID* vs. treated WT cells (1, 3, 5, 7) and NAA-treated *dcp2Δpab1-AID* vs. treated *dcp2Δ* cells (2, 4, 6, 8). Outlier mRNAs (0-6) with log2ΔmRNA for >4.0 or <-4.0 were removed. **(C)** Density scatterplot of $log_2$ fold-changes in relative numbers of ribosome protected fragments (RPFs) ($log_2$ΔRibo) and $log_2$ fold-changes in relative protein abundance measured by TMT-MS ($log_2$ΔProtein) for 4327 yeast mRNAs in NAA-treated *dcp2Δpab1-AID* vs. treated *dcp2Δ* cells, showing the Spearman correlation coefficient ($R_s$) and its p-value. **(D)** Correlation analysis of normalized peptide abundances between biological replicates determined by TMT-MS analysis. Correlation matrix showing Spearman correlation coefficients calculated for pair-wise comparisons of log2 Cyclic Loess Normalized Exclusive MS1 intensities for all expressed proteins among 6 TMT-MS analyses conducted for 3 biological replicates (rep1-rep3) of NAA-treated PMY3 (*dcp2Δ,*) and PRY1 (*dcp2Δpab1AID*). The correlation coefficients for all comparisons are >0.98 with a P-value ≈ 0.

**Fig. S8. Properties of mRNAs preferentially up-regulated or down-regulated in relative TE on Pab1-AID depletion. (A)** Boxplot showing CDS lengths of all expressed mRNAs, TE_dn_*pab1* and TE_up_*pab1* groups identified by overlaps in Fig S6C, and SCL mRNAs. Outliers (0-19) with CDS lengths >10000 and <100 nts were removed. **(B)** Boxplot showing log2 TE in WT (this study) of all expressed mRNAs, TE_dn_*pab1*, TE_up_*pab1* and SCL mRNAs. Outliers (0-79) with log2TE values >6.0 and <-6.0 were removed. **(C)** Proportional Venn diagrams showing overlaps between the 563 TE_dn_*pab1* mRNAs from Fig. S6C and SCL mRNAs. Hypergeometric distribution p-value is shown for the overlap. **(D)** Boxplot showing log2 WT steady-state mRNA half-life in min (from Neymotin et al., 2014) for all expressed mRNAs, TE_dn_*pab1,* TE_up_*pab1* mRNAs and SCL mRNAs (Costello et al., 2015). **(E)** Boxplot showing log2 steady state transcript abundance in molecules per dry cellular weight (pgDW) (from Lahtvee et al. 2017) for all expressed mRNAs and TE_dn_*pab1*, TE_up_*pab1* and SCL mRNAs. Outlier mRNAs (0-14) with log2 transcript abundance >6.0 and <-6.0 were removed. **(F)** Boxplot showing stAI values (data from Radhakrishnan et al., 2016) for all expressed mRNAs, TE_dn_*pab1*, TE_up_*pab1* mRNAs and SCL mRNAs. **(G)** Boxplot of median pA tail lengths (from this study) calculated for each of 5476 yeast genes and for the TE_dn_*pab1* and TE_up_*pab1* groups in NAA-treated WT or treated *pab1-AID* cells. Outliers with median pA tail lengths >100 nt (0-4) were removed to expand y-axis. **(H)** Boxplot showing log2 fold-changes in IP/Total RNA immunoprecipitated with TAP-tagged versions of the indicated proteins in the RIP-Seq data of Costello et al. (2015) for all expressed mRNAs and TE_dn_*pab1* mRNAs. Outliers (0-28) with log2FC values of >4.0 or <-4.0 were removed.

**Fig. S9. Hypothetical competition model to account for TE reductions at limiting Pab1-AID that are suppressed by *dcp2Δ*.** We assume that binding of eIF4E/eIF4G to mRNA is sufficient for efficient mRNA translation regardless of Pab1 binding to the pA tail, with Pab1 serving only to stabilize eIF4G binding to eIF4E and impede formation of inactive eIF4E/4E-BP mRNPs instead. **(i)** In WT cells Pab1 stabilizes eIF4G binding to mRNA, enabling eIF4G to outcompete a 4E-BP, and form eIF4F-mRNA complexes activated for translation initiation (*right*) versus inactive mRNPs lacking eIF4G (*left*). **(ii)** Depletion of Pab1-AID confers a 40% reduction in mRNA (mRNAs lost from the cell are depicted as partially transparent), increasing competition between eIF4G and 4E-BP for mRNA. At limiting Pab1, the proportion of the remaining mRNAs that bind eIF4G to form activated mRNPs is reduced with commensurate increase in the fraction binding 4E-BP instead, leading to a decrease in TE, equated with the fraction of total mRNA present in the activated complexes. **(iii)** Deletion of *DCP2* suppresses the loss in mRNA and enables eIF4G to bind a higher proportion of mRNA despite the absence of Pab1, restoring WT TE.

**Fig. S10. Additional properties of mRNAs preferentially up-regulated or down-regulated in relative TE on Pab1-AID depletion. (A)** Boxplot showing log2 fold-changes in IP/Total RNA immunoprecipitated with TAP-tagged versions of the indicated proteins in the RIP-Seq data of Costello et al. (2015) for all expressed mRNAs and TE_up_*pab1* mRNAs. Outliers (0-28) with log2FC values of >4.0 or <-4.0 were removed. **(B)** Proportional Venn diagrams showing overlaps between the 617 TE_up_*pab1* mRNAs from Fig. S6C and SCL mRNAs. Hypergeometric distribution p-value is shown for the overlap.

**Fig. S11. Depletion of Pab1-AID leads to reduced levels of total 40S and 60S ribosomal subunits. (A)** Quantification of total 40S and 60S ribosomal subunits in *pab1-AID* vs. WT cells both treated with auxin. Representative $A_{260}$ profiles of equal proportions of cell extracts obtained from WT (blue) and *pab1-AID* (orange) cultures are shown. **(B)** Histogram showing the combined areas under the 40S and 60S peaks normalized to the $OD_{600}$ of the cell cultures calculated from four biological replicates of auxin-treated *pab1-AID* and WT cells, setting the mean WT value to unity. An unpaired Student's t-test indicates a highly significant difference in the means calculated from the biological replicates of the two different strains (**p < 0.001).

**Fig. S12. Schematic summary of hypothetical models proposed to account for TE changes at limiting Pab1 that are suppressed by *dcp2Δ*.** Changes in mRNA levels and TE are depicted schematically for the average mRNA (col. 1), TE_dn_*pab1* mRNAs hyperdependent on Pab1 for TE in *DCP2* cells (col. 2) and TE_up_*pab1* mRNAs hypodependent on Pab1 for TE in *DCP2* cells in either auxin-treated WT (*upper*), *pab1-AID* (*middle*), or *dcp2Δ pab1-AID* cells (*lower*). The increased inhibition by RBPs proposed for the TE_dn_*pab1* group in *pab1-AID* cells, which is dampened in *dcp2Δ pab1-AID* cells is depicted explicitly in Fig. S9. See Discussion for further details.

**Fig. S13. Properties of mRNAs preferentially up-regulated or down-regulated in relative TE on Pab1-AID depletion in *dcp2Δ*. (A)** Boxplot showing log2 fold-changes in IP/Total RNA immunoprecipitated with TAP-tagged versions of the indicated proteins in the RIP-Seq data of Costello et al. (2015) for all mRNAs and TE_dn_*dp* mRNAs defined in Fig. 6B. Outliers (0-28) with log2FC values of >4.0 or <-4.0 were removed. **(B)** Boxplot showing CDS lengths of all mRNAs, TE_dn_*dp* and TE_up_*dp* groups identified in Fig. 6B. Outliers (0-19) with CDS lengths >10000 and <100 nts were removed. **(C)** Boxplot showing log2 TE in WT (this study) of all mRNAs, TE_dn_*dp* and TE_up_*dp* groups. Outliers (0-79) with log2TE values >6.0 and <-6.0 were removed. **(D)** Boxplot showing log2 steady state transcript abundance in molecules per dry cellular weight (pgDW) (from Lahtvee et al. 2017) for all expressed mRNAs and TE_dn_*dp* and TE_up_*dp* mRNAs. Outlier mRNAs (0-14) with log2 transcript abundance >6.0 and <-6.0

were removed. **(E)** Boxplot showing log2 WT steady-state mRNA half-life in min (from Neymotin et al., 2014) for all expressed mRNAs, TE_dn_*dp* and TE_up_*dp* groups. **(F)** Boxplot showing stAI values (data from Radhakrishnan et al., 2016) for all expressed mRNAs, TE_dn_*dp*, TE_up_*dp* mRNAs. **(G)** Proportional Venn diagram showing overlaps between the 116 TE_dn_*dp* mRNAs and SCL mRNAs. Hypergeometric distribution p-value is shown for the overlap.

### SUPPLEMENTARY TABLES

**Table S1: Yeast strains used in this study.**

| Strain name | Abbreviated genotype | Complete genotype | Source |
|---|---|---|---|
| F2147 | W303 | *MATa ade2-1 ura3-1 his3-11,15 trp1-1 leu2-3 112 can1-100* | (62) |
| CFY1016 | *dcp2Δ* | *MATa ade2-1 ura3-1 his3-11,15 trp1-1 leu2-3,112 can1-100 dcp2Δ::HIS3* | (62) |
| PMY1 | *OsTIR-myc* | *MATa ade2-1 ura3-1 his3-11,15 trp1-1 leu2-3,112 can1-100 lys2::*$P_{ADH1}$*-OsTIR1-7myc-*$T_{ADH1}$ | This study |
| PMY2 | *OsTIR-7myc pab1-AID*-9myc* | *MATa ade2-1 ura3-1 his3-11,15 trp1-1 leu2-3,112 can1-100 lys2::*$P_{ADH1}$*-OsTIR1-7myc-*$T_{ADH1}$ *pab1-AID* $^{(71\text{-}114)}$*-9myc::hphNT1* | This study |
| PMY3 | *dcp2Δ OsTIR-7myc* | *MATa ade2-1 ura3-1 his3-11,15 trp1-1 leu2-3,112 can1-100 dcp2Δ::HIS3 lys2::*$P_{ADH1}$*-OsTIR1-7myc-*$T_{ADH1}$ | This study |
| PRY1 | *dcp2Δ OsTIR-7myc pab1-AID*-9myc* | *MATa ade2-1 ura3-1 his3-11,15 trp1-1 leu2-3,112 can1-100 dcp2Δ::HIS3 lys2::* $P_{ADH1}$*-OsTIR1-7myc-*$T_{ADH1}$ *pab1-AID* $^{(71\text{-}114)}$*-9myc::hphNT1* | This study |
| FY2180 | *spt6-1004* | *MATα leu2-Δ1 his4-912δ lys2-128δ FLAG-spt6-1004* | Fred Winston |
| FY2181 | *SPT6* | *MATα leu2-Δ1 his4-912δ lys2-128δ* | Fred Winston |
| H4293/ DG184 | *set1Δ set1Δ* | *MATa his3-Δ1 leu2-Δ0 ura3-Δ0 met15-Δ0 set2Δ::KanMX4 set1Δ::hphMX4* | (63) |
| BY4741 | *SET1 SET2* | *MATa his3-Δ1 leu2-Δ0 ura3-Δ0 met15-Δ0* | (64) |

**Table S2: Plasmids used in this study.**

| Plasmid | Relevant description | Source or reference |
|---|---|---|
| pQZ145 | *DCP2* in pRS316 | (23) |
| pRS316 | CEN6, *URA3* vector | (65) |
| p6022 | pIS385-*OsTIR1* | Carl Wu |
| pHQ2122 | pIS385-*OsTIR1-7myc* | This study |
| pIS385 | *LYS2* disintegrator plasmid | (20) |
| p6023 | *pHyg-AID*-9Myc* | (66) |

**Table S3: Oligonucleotide primers used in this study.**

| Primer Name | Sequence | Notes |
|---|---|---|
| PYON1 | 5'-AGATTGCCACCGACGAAAAC-3' | Forward primer within *PAB1* ORF |
| PYON2 | 5'-TCTGGCTTGGATTTGTTGAGC-3' | Reverse primer within *PAB1* ORF |
| PYON3 | 5'-TTCCCAAGAAATGCCAACGA-3' | Forward primer within *PAB1* ORF to verify *pab1-AID* tagging |
| PYON4 | 5'-GCGGTGATGAATAGCAAGGT-3' | Reverse primer in 3' UTR of *PAB1* to verify *pab1-AID* tagging |
| PYON42 | 5'-GATTAAATCATTTGCTCAAT-3' | Forward primer within *DCP2* ORF |
| PYON43 | 5'-GCAACTCTCTACCGGGATCA -3' | Reverse primer within *DCP2* ORF |
| Pab1_deg_F | 5'-CAAAGAAGCTTCTGCTGCCTATGAGTCTTTCAAAAAGGAGCAAGAACAACAAACTGAGCAAGCTCCTAAAGATCCAGCCAAACCT-3' | Forward primer for *AID-9myc* tagging of *PAB1* containing sequences 64bp 5' to STOP codon |
| Pab1_deg_R | 5'-GAAGGGAGAAAAAAAAGATGATAAGTTTGTTGAGTAGGGAAGTAGGTGATTACATAGAGCAGTTAAAGCCTTCGAGCGTCCC-3' | Reverse primer for *AID-9myc* tagging of *PAB1* containing sequences 61bp 3' to STOP codon |
| Primer 2157 | 5'-CATCGGTAAATACGCTAAGCA-3' | Forward primer with BlpI site for cloning *OsTIR* |
| Primer 2158 | 5'-CACCGTTAATTAAAATCTTGACGAAGTTAGG-3' | Reverse primer with PacI site for cloning *OsTIR* |
| Primer 688 | 5'-CGGATCCCCGGGTTAATTAA-3' | Forward primer with PacI site for cloning *7myc*-$T_{ADH1}$ |
| Primer 2159 | 5'-ATGCTGGCGGCCGCATATTACCCTGTTATCCCTAGC-3' | Reverse primer with EagI site for cloning *7myc*-$T_{ADH1}$ |
| Primer 2180 | 5'-CGATCCATCCAATTGCATAAC-3' | Reverse primer for verifying integration of *OsTIR-7-myc* at *LYS2* locus |
| Primer 2181 | 5'-AGGAAGTCCATATTGTACACC-3' | Forward primer located in 5' end of *ADH1* promoter region for checking integration of *OsTIR-7-myc* at *LYS2* locus |
| Primer 2182 | 5'-GCCAGCTGATTTACAGTTCTT-3' | Reverse primer located in 3' region of *LYS2* gene for checking integration of *OsTIR*-7myc at *LYS2* locus |

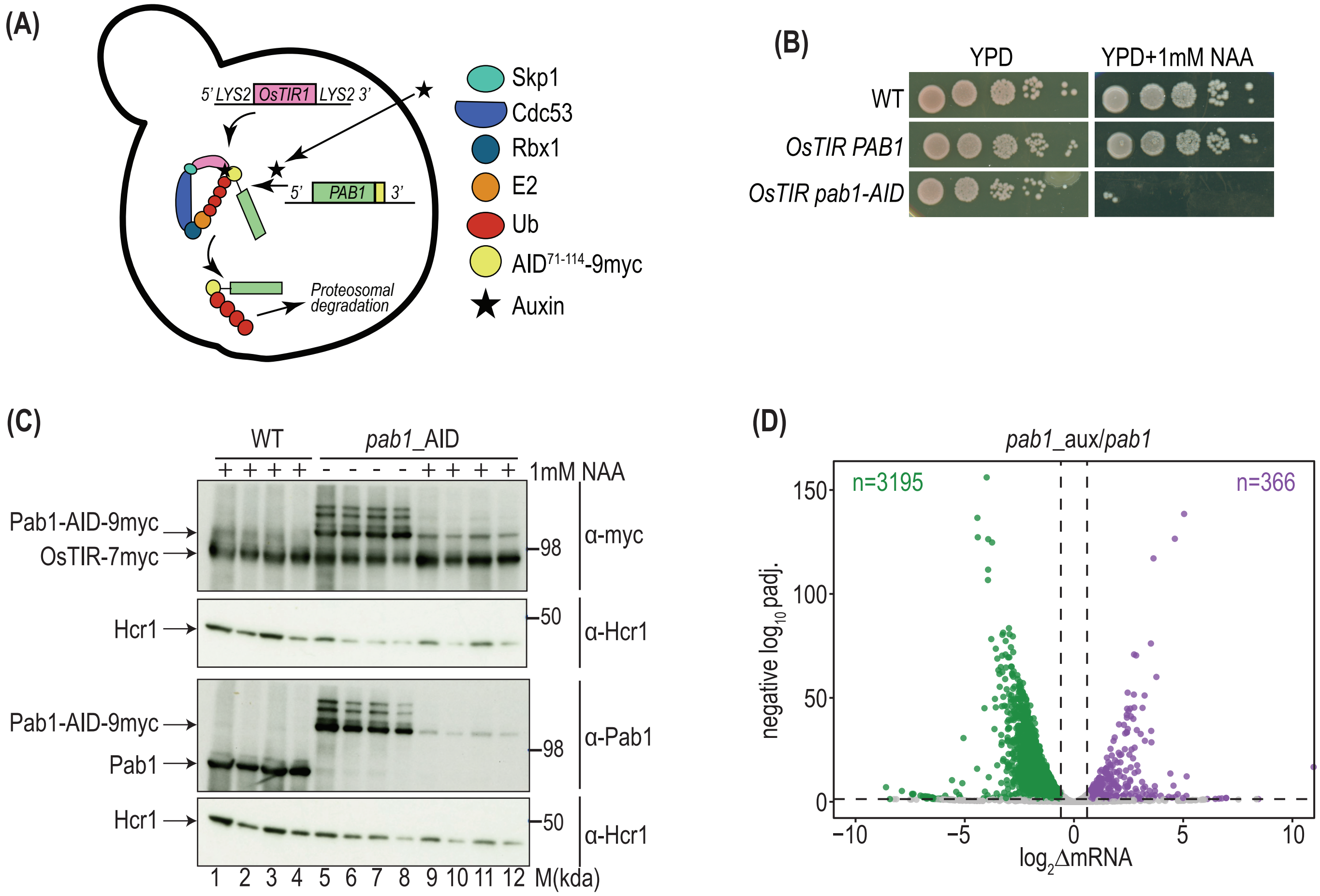


| Protein | $log_2$Δprotein *pab1*/WT (TMT MS) | padj. |
|---|---|---|
| Pab1 | -3.647 | 1.3e-08 |

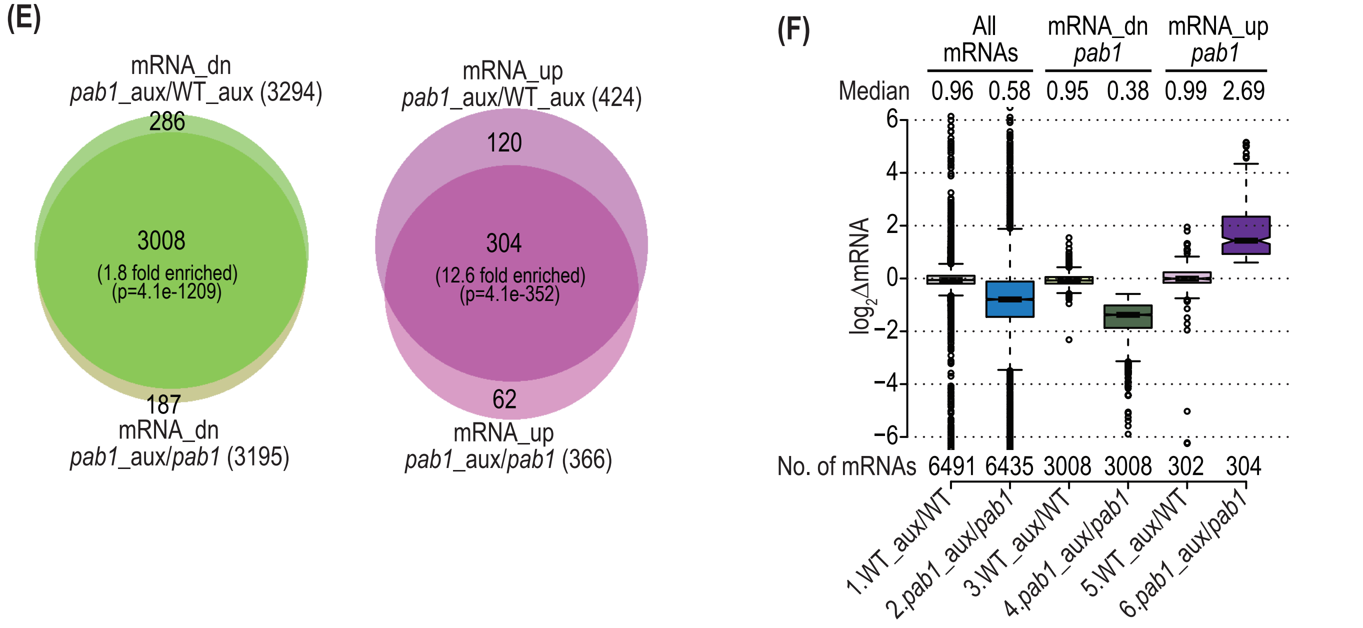


**Fig.S1**

**(G)**

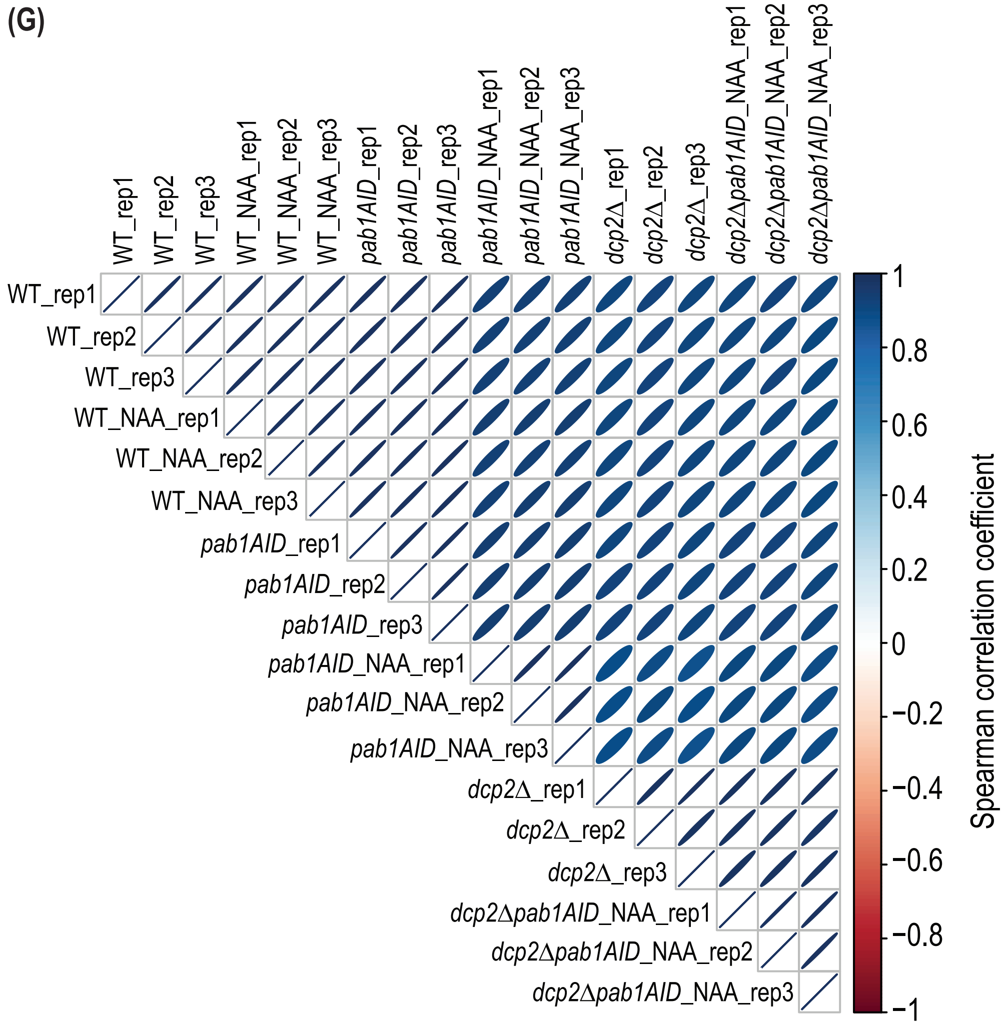
WT_rep1
WT_rep2
WT_rep3
WT_NAA_rep1
WT_NAA_rep2
WT_NAA_rep3
pab1AID_rep1
pab1AID_rep2
pab1AID_rep3
pab1AID_NAA_rep1
pab1AID_NAA_rep2
pab1AID_NAA_rep3
dcp2Δ_rep1
dcp2Δ_rep2
dcp2Δ_rep3
dcp2Δpab1AID_NAA_rep1
dcp2Δpab1AID_NAA_rep2
dcp2Δpab1AID_NAA_rep3
1
0.8
0.6
0.4
0.2
0
−0.2
−0.4
−0.6
−0.8
−1
Spearman correlation coefficient

**Fig.S1**

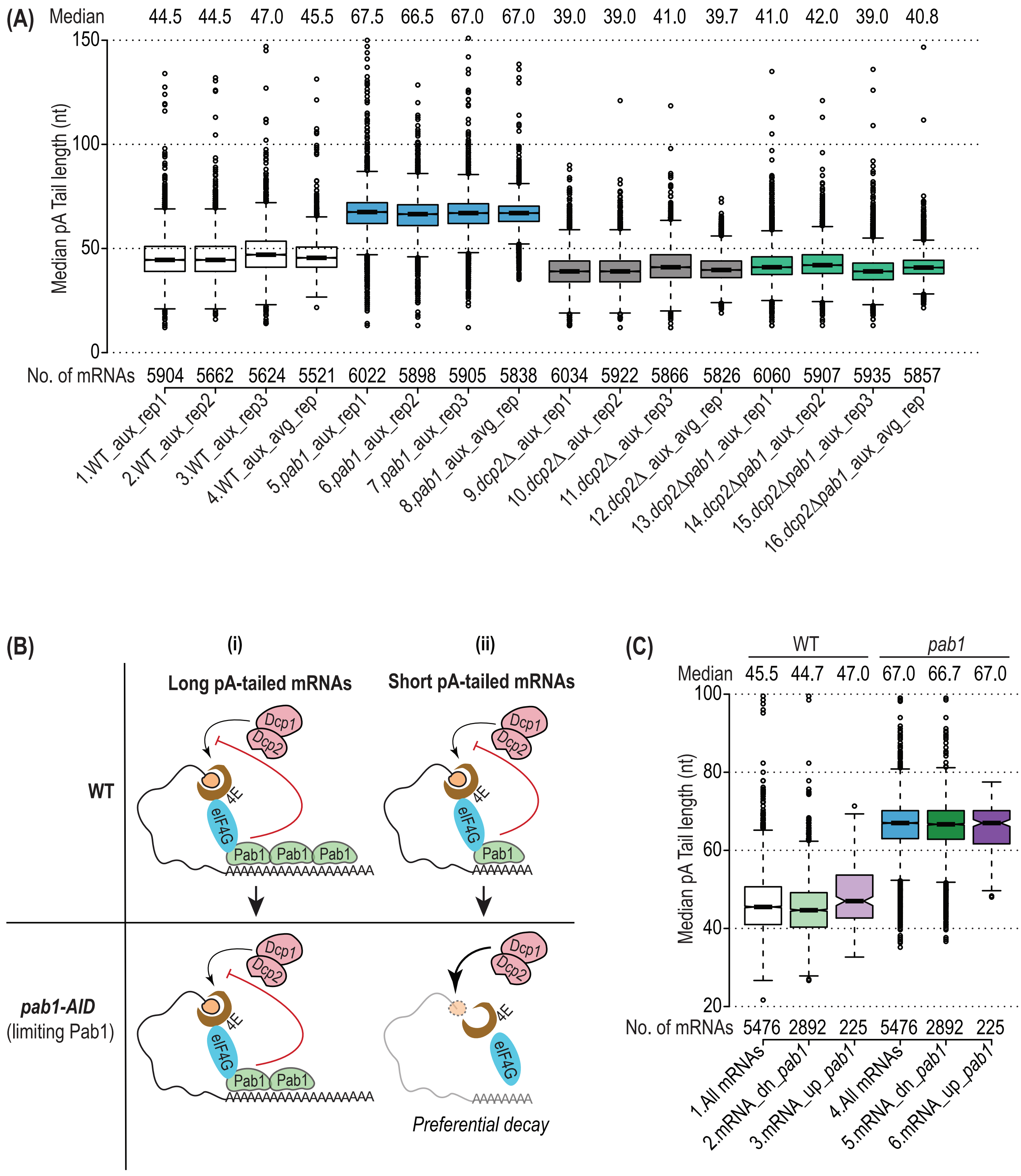

(A)
Median
44.5 44.5 47.0 45.5 67.5 66.5 67.0 67.0 39.0 39.0 41.0 39.7 41.0 42.0 39.0 40.8
Median pA Tail length (nt)
150
100
50
0
No. of mRNAs
5904 5662 5624 5521 6022 5898 5905 5838 6034 5922 5866 5826 6060 5907 5935 5857
1.WT_aux_rep1
2.WT_aux_rep2
3.WT_aux_rep3
4.WT_aux_avg_rep
5.pab1_aux_rep1
6.pab1_aux_rep2
7.pab1_aux_rep3
8.pab1_aux_avg_rep
9.dcp2Δ_aux_rep1
10.dcp2Δ_aux_rep2
11.dcp2Δ_aux_rep3
12.dcp2Δ_aux_avg_rep
13.dcp2Δpab1_aux_rep1
14.dcp2Δpab1_aux_rep2
15.dcp2Δpab1_aux_rep3
16.dcp2Δpab1_aux_avg_rep
(B)
(i)
(ii)
Long pA-tailed mRNAs
Short pA-tailed mRNAs
WT
Dcp1
Dcp2
4E
eIF4G
Pab1 Pab1 Pab1
AAAAAAAAAAAAAAAAAA
Pab1
AAAAAAA
pab1-AID
(limiting Pab1)
Pab1 Pab1
Preferential decay
(C)
WT
pab1
Median
45.5 44.7 47.0 67.0 66.7 67.0
Median pA Tail length (nt)
100
80
60
40
20
No. of mRNAs
5476 2892 225 5476 2892 225
1.All mRNAs
2.mRNA_dn_pab1
3.mRNA_up_pab1
4.All mRNAs
5.mRNA_dn_pab1
6.mRNA_up_pab1


**Fig.S2**

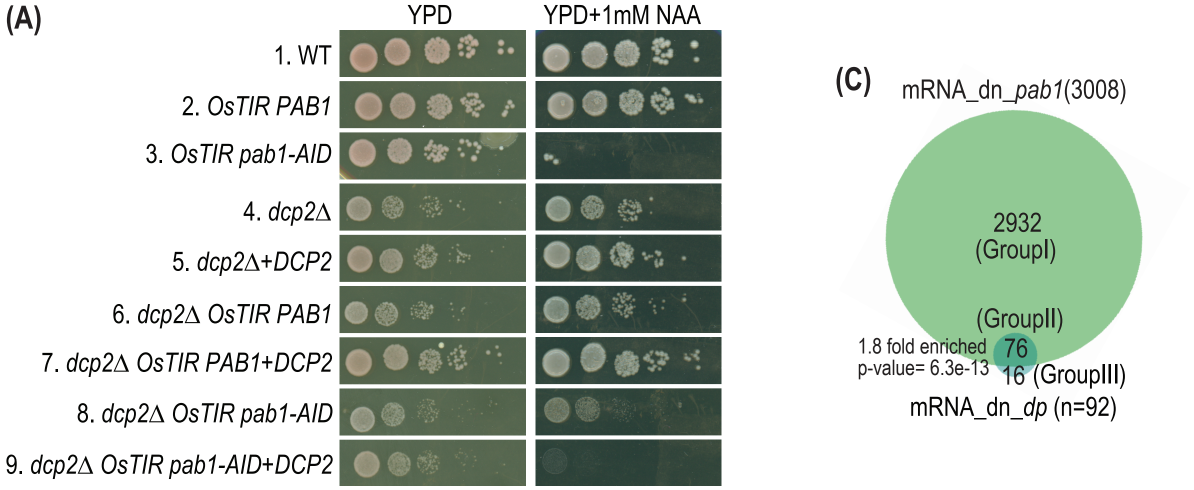


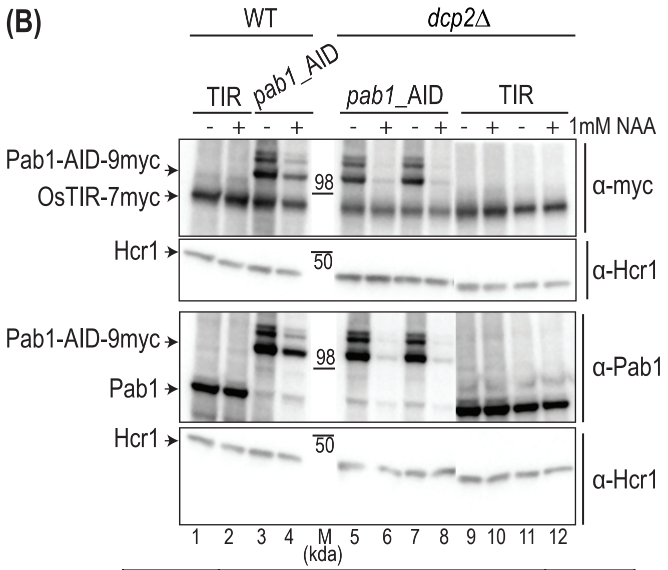


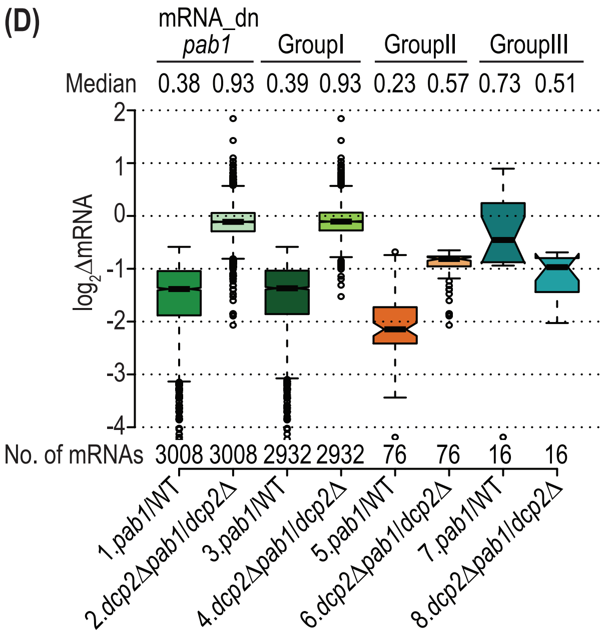


| Protein | $log_2$Δprotein *dcp2Δpab1/dcp2Δ* (TMT-MS) | padj. |
|---|---|---|
| Pab1 | -4.434 | 1.3e-07 |

**Fig.S3**

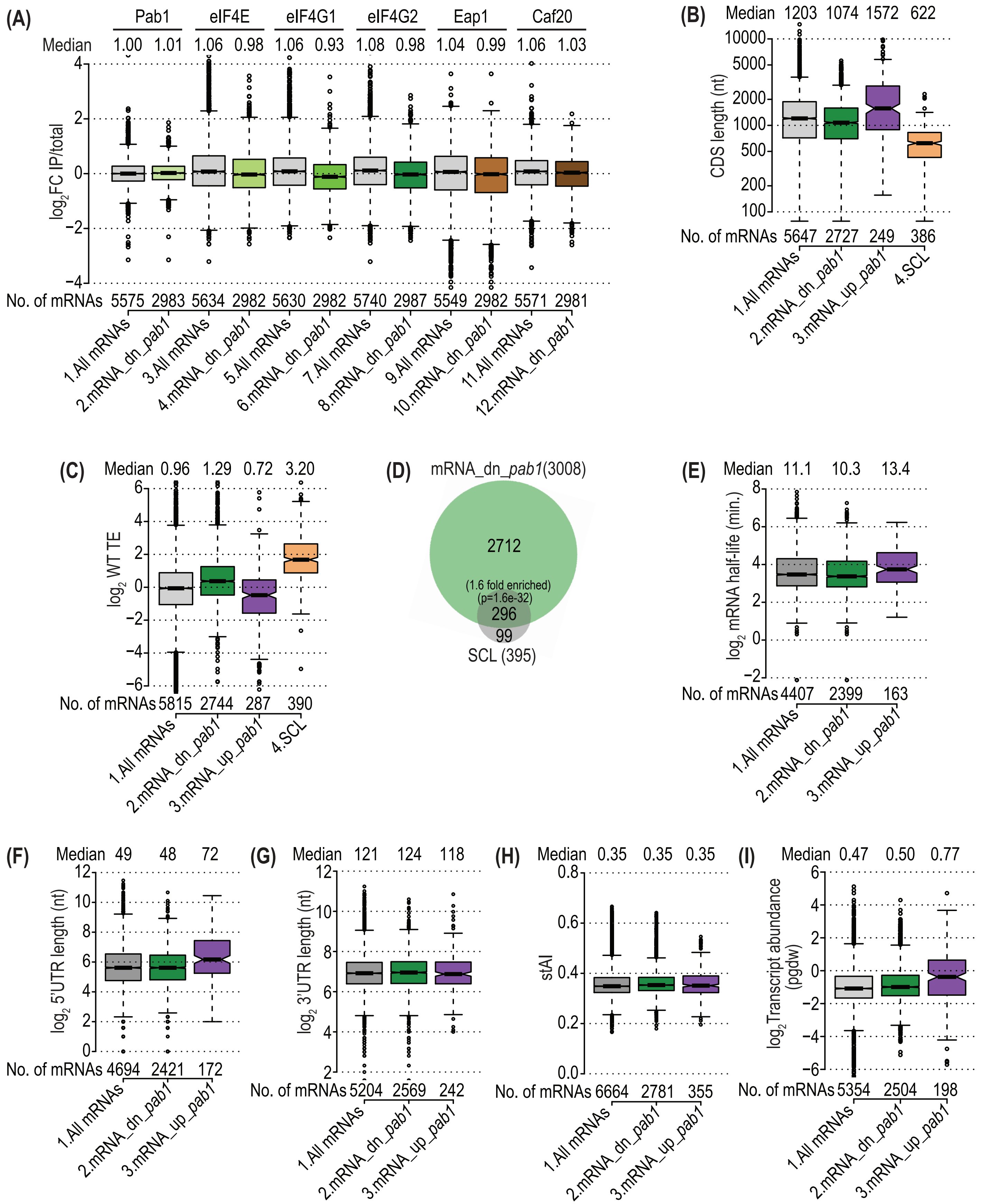
(A)
Pab1
eIF4E
eIF4G1
eIF4G2
Eap1
Caf20
Median 1.00 1.01 1.06 0.98 1.06 0.93 1.08 0.98 1.04 0.99 1.06 1.03
log2FC IP/total
No. of mRNAs 5575 2983 5634 2982 5630 2982 5740 2987 5549 2982 5571 2981
1.All mRNAs
2.mRNA_dn_pab1
3.All mRNAs
4.mRNA_dn_pab1
5.All mRNAs
6.mRNA_dn_pab1
7.All mRNAs
8.mRNA_dn_pab1
9.All mRNAs
10.mRNA_dn_pab1
11.All mRNAs
12.mRNA_dn_pab1
(B)
Median 1203 1074 1572 622
CDS length (nt)
No. of mRNAs 5647 2727 249 386
1.All mRNAs
2.mRNA_dn_pab1
3.mRNA_up_pab1
4.SCL
(C)
Median 0.96 1.29 0.72 3.20
log2 WT TE
No. of mRNAs 5815 2744 287 390
1.All mRNAs
2.mRNA_dn_pab1
3.mRNA_up_pab1
4.SCL
(D)
mRNA_dn_pab1(3008)
2712
(1.6 fold enriched)
(p=1.6e-32)
296
99
SCL (395)
(E)
Median 11.1 10.3 13.4
log2 mRNA half-life (min.)
No. of mRNAs 4407 2399 163
1.All mRNAs
2.mRNA_dn_pab1
3.mRNA_up_pab1
(F)
Median 49 48 72
log2 5'UTR length (nt)
No. of mRNAs 4694 2421 172
1.All mRNAs
2.mRNA_dn_pab1
3.mRNA_up_pab1
(G)
Median 121 124 118
log2 3'UTR length (nt)
No. of mRNAs 5204 2569 242
1.All mRNAs
2.mRNA_dn_pab1
3.mRNA_up_pab1
(H)
Median 0.35 0.35 0.35
stAI
No. of mRNAs 6664 2781 355
1.All mRNAs
2.mRNA_dn_pab1
3.mRNA_up_pab1
(I)
Median 0.47 0.50 0.77
log2 Transcript abundance (pgdw)
No. of mRNAs 5354 2504 198
1.All mRNAs
2.mRNA_dn_pab1
3.mRNA_up_pab1

**Fig.S4**

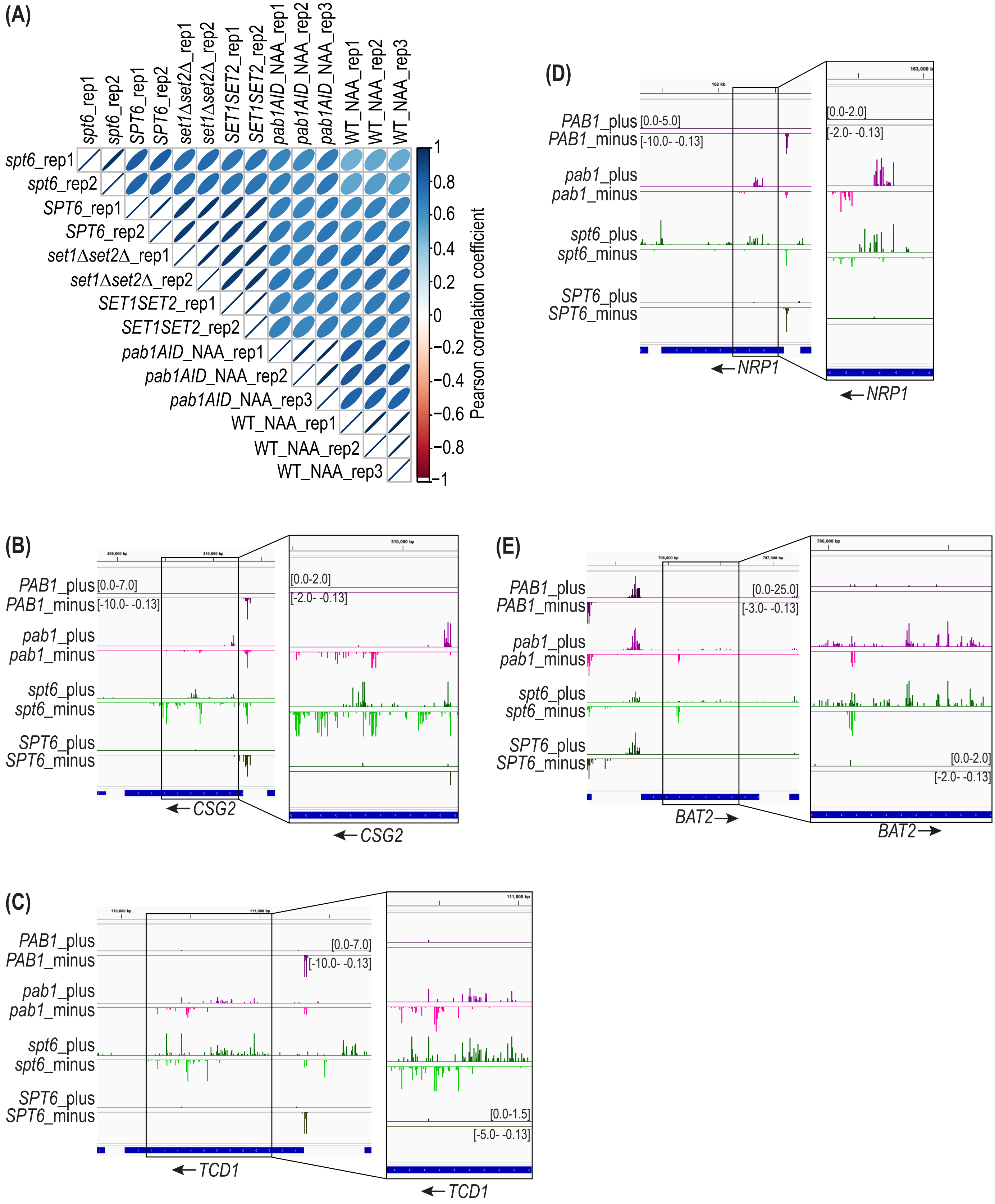
(A)
spt6_rep1
spt6_rep2
SPT6_rep1
SPT6_rep2
set1Δset2Δ_rep1
set1Δset2Δ_rep2
SET1SET2_rep1
SET1SET2_rep2
pab1AID_NAA_rep1
pab1AID_NAA_rep2
pab1AID_NAA_rep3
WT_NAA_rep1
WT_NAA_rep2
WT_NAA_rep3
Pearson correlation coefficient
1
0.8
0.6
0.4
0.2
0
−0.2
−0.4
−0.6
−0.8
−1
(B)
PAB1_plus [0.0-7.0]
PAB1_minus [-10.0- -0.13]
pab1_plus
pab1_minus
spt6_plus
spt6_minus
SPT6_plus
SPT6_minus
[0.0-2.0]
[-2.0- -0.13]
CSG2
(C)
PAB1_plus
PAB1_minus
[0.0-7.0]
[-10.0- -0.13]
pab1_plus
pab1_minus
spt6_plus
spt6_minus
SPT6_plus
SPT6_minus
[0.0-1.5]
[-5.0- -0.13]
TCD1
(D)
PAB1_plus [0.0-5.0]
PAB1_minus [-10.0- -0.13]
[0.0-2.0]
[-2.0- -0.13]
pab1_plus
pab1_minus
spt6_plus
spt6_minus
SPT6_plus
SPT6_minus
NRP1
(E)
PAB1_plus
PAB1_minus
[0.0-25.0]
[-3.0- -0.13]
pab1_plus
pab1_minus
spt6_plus
spt6_minus
SPT6_plus
SPT6_minus
[0.0-2.0]
[-2.0- -0.13]
BAT2

**Fig.S5**

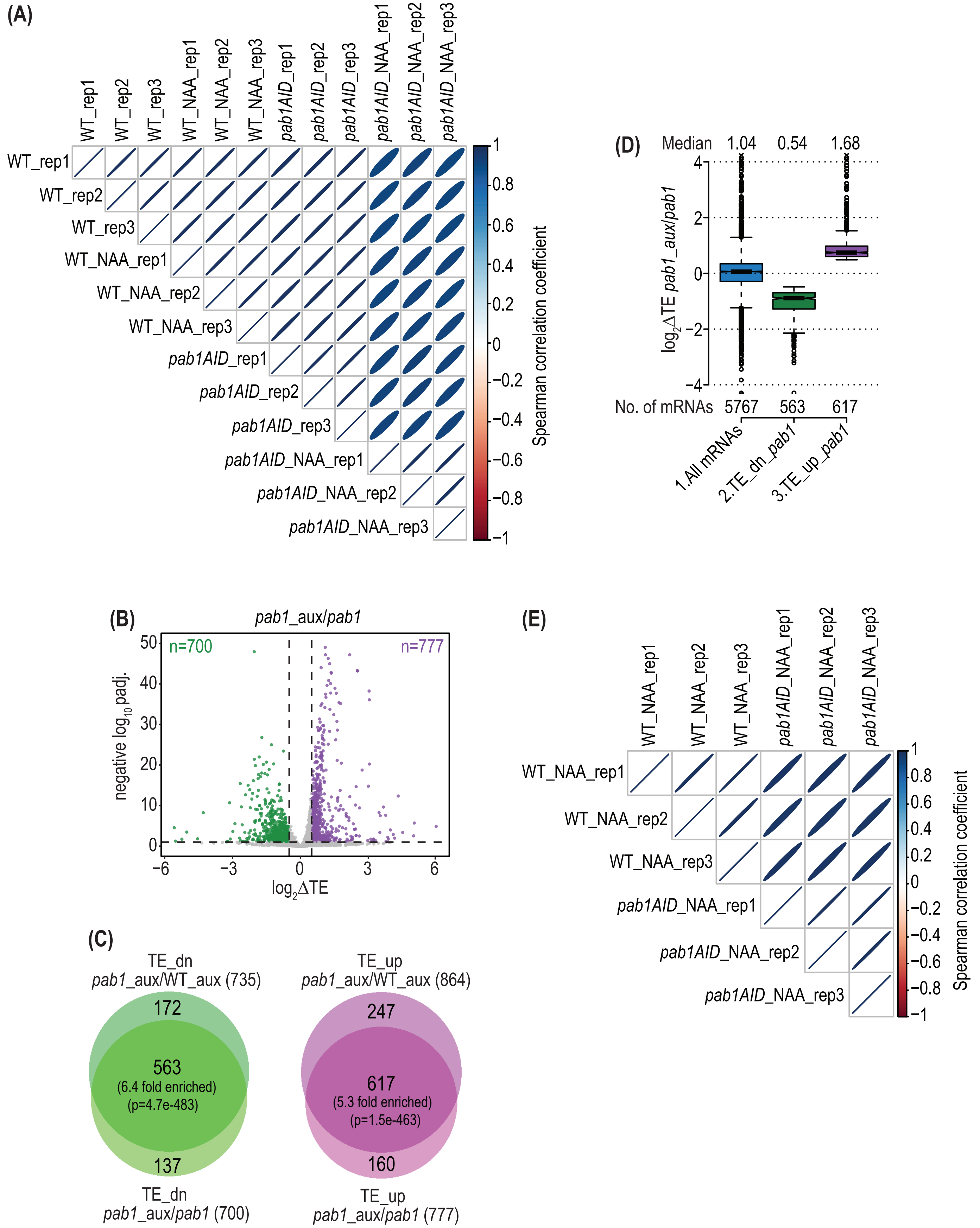
(A)
WT_rep1
WT_rep2
WT_rep3
WT_NAA_rep1
WT_NAA_rep2
WT_NAA_rep3
pab1AID_rep1
pab1AID_rep2
pab1AID_rep3
pab1AID_NAA_rep1
pab1AID_NAA_rep2
pab1AID_NAA_rep3
Spearman correlation coefficient
(D)
Median 1.04 0.54 1.68
log2ΔTE pab1_aux/pab1
No. of mRNAs 5767 563 617
1.All mRNAs
2.TE_dn_pab1
3.TE_up_pab1
(B)
pab1_aux/pab1
n=700
n=777
negative log10 padj.
log2ΔTE
(E)
WT_NAA_rep1
WT_NAA_rep2
WT_NAA_rep3
pab1AID_NAA_rep1
pab1AID_NAA_rep2
pab1AID_NAA_rep3
Spearman correlation coefficient
(C)
TE_dn
pab1_aux/WT_aux (735)
172
563
(6.4 fold enriched)
(p=4.7e-483)
137
TE_dn
pab1_aux/pab1 (700)
TE_up
pab1_aux/WT_aux (864)
247
617
(5.3 fold enriched)
(p=1.5e-463)
160
TE_up
pab1_aux/pab1 (777)

**Fig.S6**

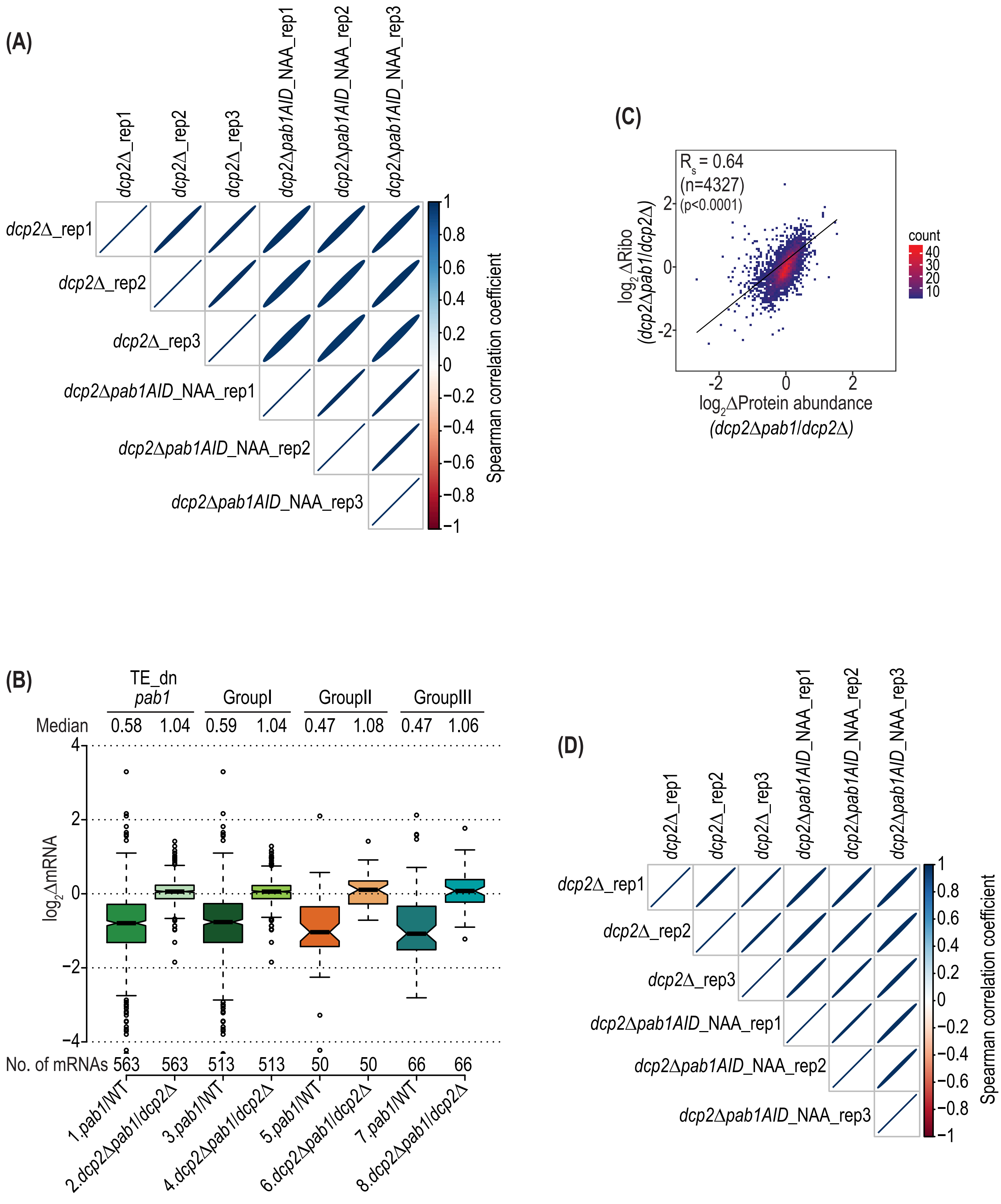

(A)
dcp2Δ_rep1
dcp2Δ_rep2
dcp2Δ_rep3
dcp2Δpab1AID_NAA_rep1
dcp2Δpab1AID_NAA_rep2
dcp2Δpab1AID_NAA_rep3
Spearman correlation coefficient
1
0.8
0.6
0.4
0.2
0
−0.2
−0.4
−0.6
−0.8
−1
(C)
Rs = 0.64
(n=4327)
(p<0.0001)
log2ΔRibo
(dcp2Δpab1/dcp2Δ)
log2ΔProtein abundance
(dcp2Δpab1/dcp2Δ)
count
40
30
20
10
(B)
TE_dn
pab1
GroupI
GroupII
GroupIII
Median
0.58
1.04
0.59
1.04
0.47
1.08
0.47
1.06
log2ΔmRNA
No. of mRNAs
563
563
513
513
50
50
66
66
1.pab1/WT
2.dcp2Δpab1/dcp2Δ
3.pab1/WT
4.dcp2Δpab1/dcp2Δ
5.pab1/WT
6.dcp2Δpab1/dcp2Δ
7.pab1/WT
8.dcp2Δpab1/dcp2Δ
(D)
dcp2Δ_rep1
dcp2Δ_rep2
dcp2Δ_rep3
dcp2Δpab1AID_NAA_rep1
dcp2Δpab1AID_NAA_rep2
dcp2Δpab1AID_NAA_rep3
Spearman correlation coefficient


**Fig.S7**

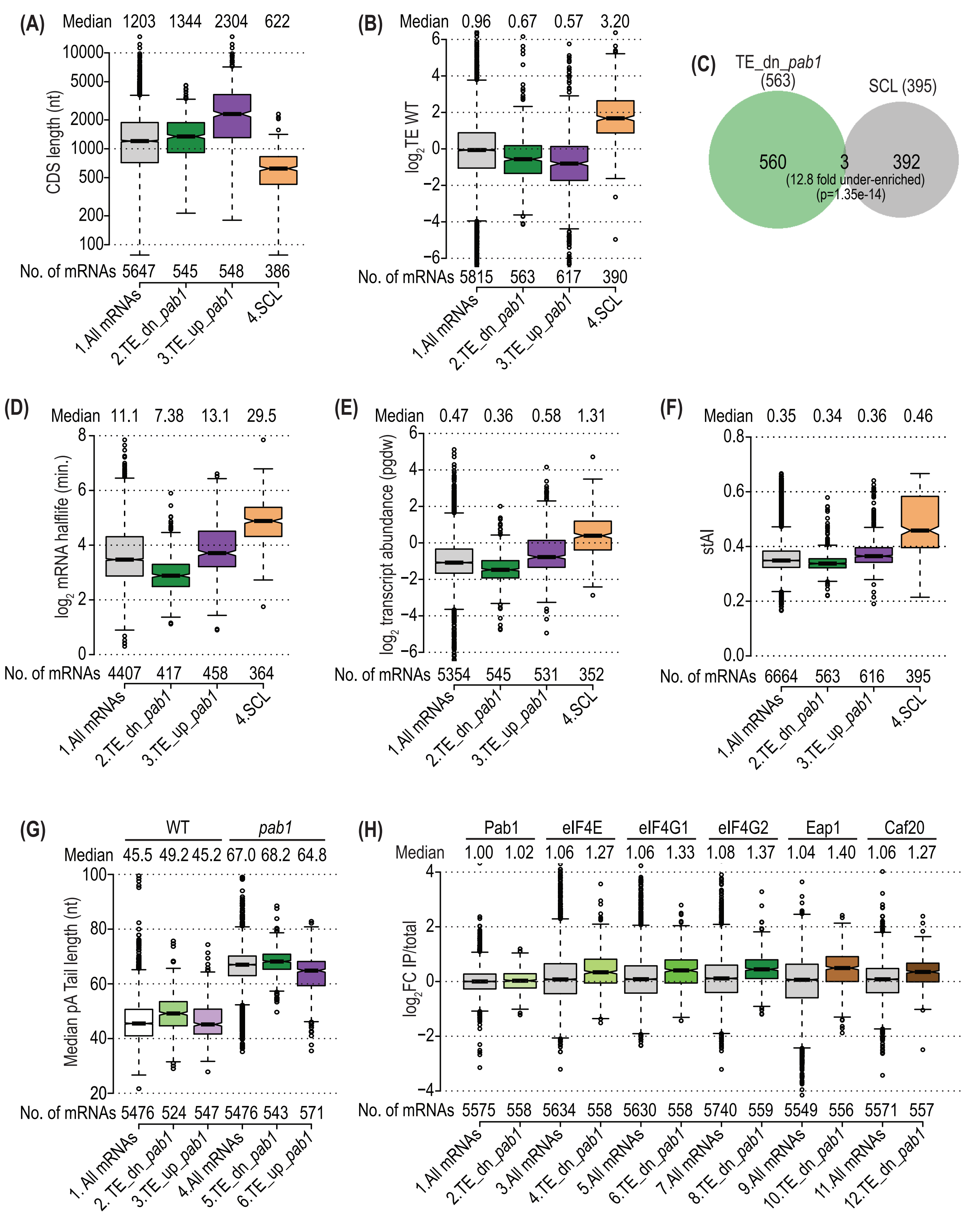
(A)
Median 1203 1344 2304 622
CDS length (nt)
10000
5000
2000
1000
500
200
100
No. of mRNAs 5647 545 548 386
1.All mRNAs
2.TE_dn_pab1
3.TE_up_pab1
4.SCL
(B)
Median 0.96 0.67 0.57 3.20
log2TE WT
6
4
2
0
−2
−4
−6
No. of mRNAs 5815 563 617 390
1.All mRNAs
2.TE_dn_pab1
3.TE_up_pab1
4.SCL
(C)
TE_dn_pab1 (563)
SCL (395)
560
3
392
(12.8 fold under-enriched)
(p=1.35e-14)
(D)
Median 11.1 7.38 13.1 29.5
log2 mRNA halflife (min.)
8
6
4
2
0
No. of mRNAs 4407 417 458 364
1.All mRNAs
2.TE_dn_pab1
3.TE_up_pab1
4.SCL
(E)
Median 0.47 0.36 0.58 1.31
log2 transcript abundance (pgdw)
6
4
2
0
−2
−4
−6
No. of mRNAs 5354 545 531 352
1.All mRNAs
2.TE_dn_pab1
3.TE_up_pab1
4.SCL
(F)
Median 0.35 0.34 0.36 0.46
stAI
0.8
0.6
0.4
0.2
0.0
No. of mRNAs 6664 563 616 395
1.All mRNAs
2.TE_dn_pab1
3.TE_up_pab1
4.SCL
(G)
WT
pab1
Median 45.5 49.2 45.2 67.0 68.2 64.8
Median pA Tail length (nt)
100
80
60
40
20
No. of mRNAs 5476 524 547 5476 543 571
1. All mRNAs
2. TE_dn_pab1
3.TE_up_pab1
4.All mRNAs
5.TE_dn_pab1
6.TE_up_pab1
(H)
Pab1
eIF4E
eIF4G1
eIF4G2
Eap1
Caf20
Median 1.00 1.02 1.06 1.27 1.06 1.33 1.08 1.37 1.04 1.40 1.06 1.27
log2FC IP/total
4
2
0
−2
−4
No. of mRNAs 5575 558 5634 558 5630 558 5740 559 5549 556 5571 557
1.All mRNAs
2.TE_dn_pab1
3.All mRNAs
4.TE_dn_pab1
5.All mRNAs
6.TE_dn_pab1
7.All mRNAs
8.TE_dn_pab1
9.All mRNAs
10.TE_dn_pab1
11.All mRNAs
12.TE_dn_pab1

**Fig.S8**

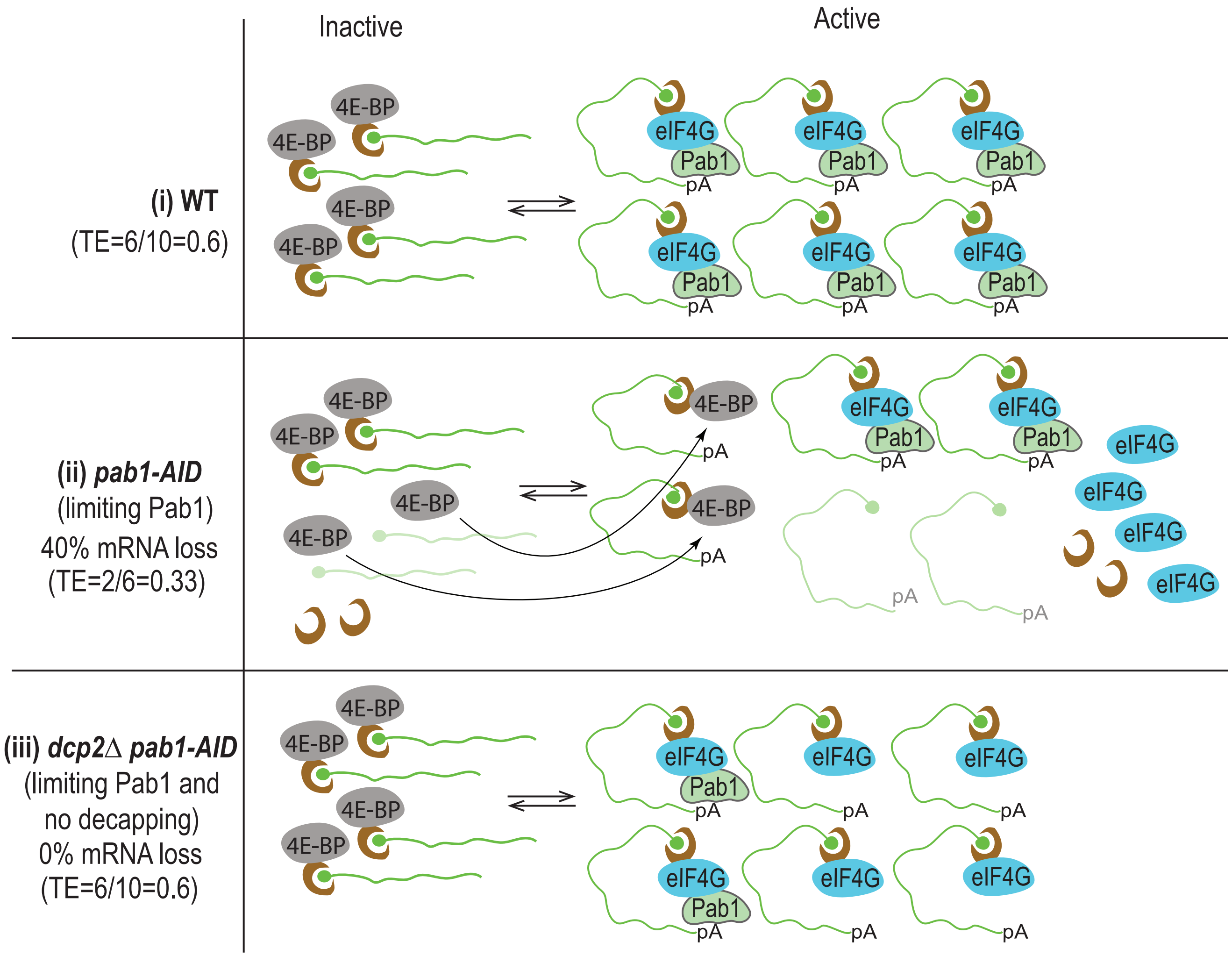

Inactive
Active
(i) WT
(TE=6/10=0.6)
4E-BP
eIF4G
Pab1
pA
(ii) pab1-AID
(limiting Pab1)
40% mRNA loss
(TE=2/6=0.33)
(iii) dcp2Δ pab1-AID
(limiting Pab1 and
no decapping)
0% mRNA loss
(TE=6/10=0.6)


Fig.S9

Fig.S9

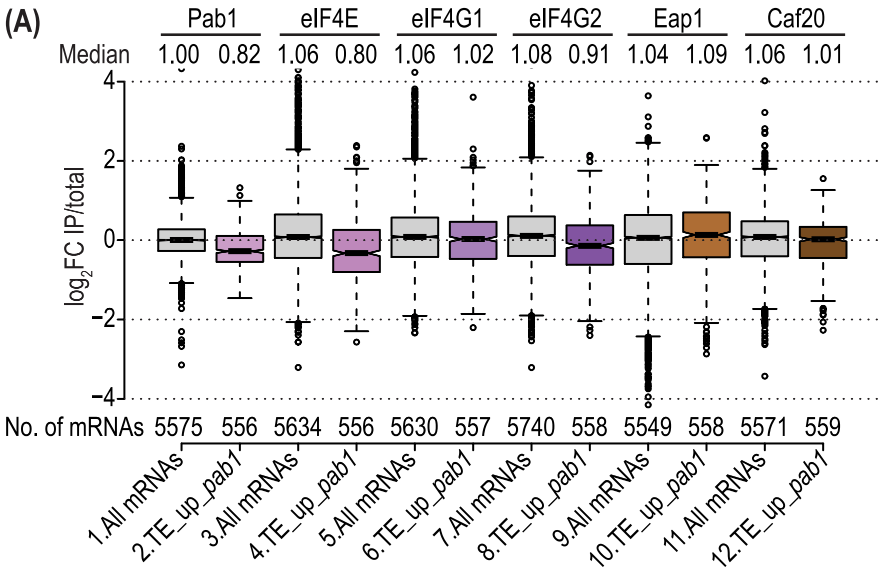
(A)
Pab1
eIF4E
eIF4G1
eIF4G2
Eap1
Caf20
Median 1.00 0.82 1.06 0.80 1.06 1.02 1.08 0.91 1.04 1.09 1.06 1.01
log2FC IP/total
4
2
0
−2
−4
No. of mRNAs 5575 556 5634 556 5630 557 5740 558 5549 558 5571 559
1.All mRNAs
2.TE_up_pab1
3.All mRNAs
4.TE_up_pab1
5.All mRNAs
6.TE_up_pab1
7.All mRNAs
8.TE_up_pab1
9.All mRNAs
10.TE_up_pab1
11.All mRNAs
12.TE_up_pab1

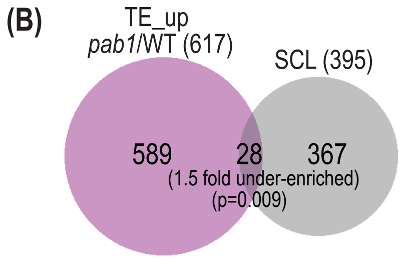
(B)
TE_up
pab1/WT (617)
SCL (395)
589
28
367
(1.5 fold under-enriched)
(p=0.009)

**Fig.S10**

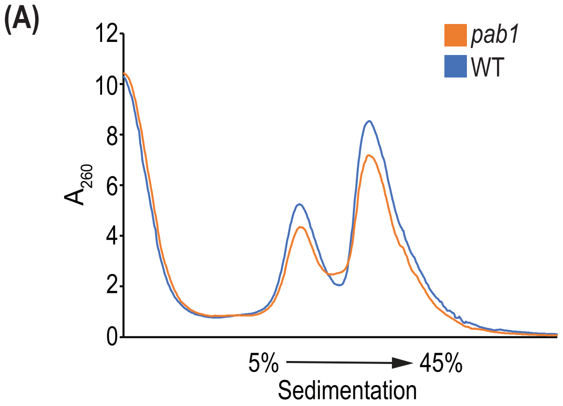
(A)
12
10
8
6
4
2
0
$A_{260}$
pab1
WT
5%
45%
Sedimentation

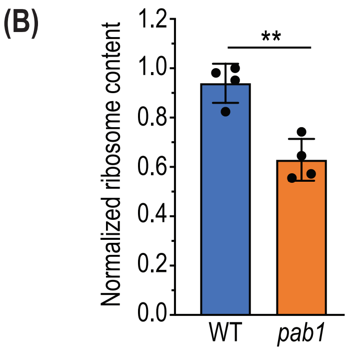
(B)
1.2
1.0
0.8
0.6
0.4
0.2
0.0
Normalized ribosome content
**
WT
pab1

**Fig.S11**

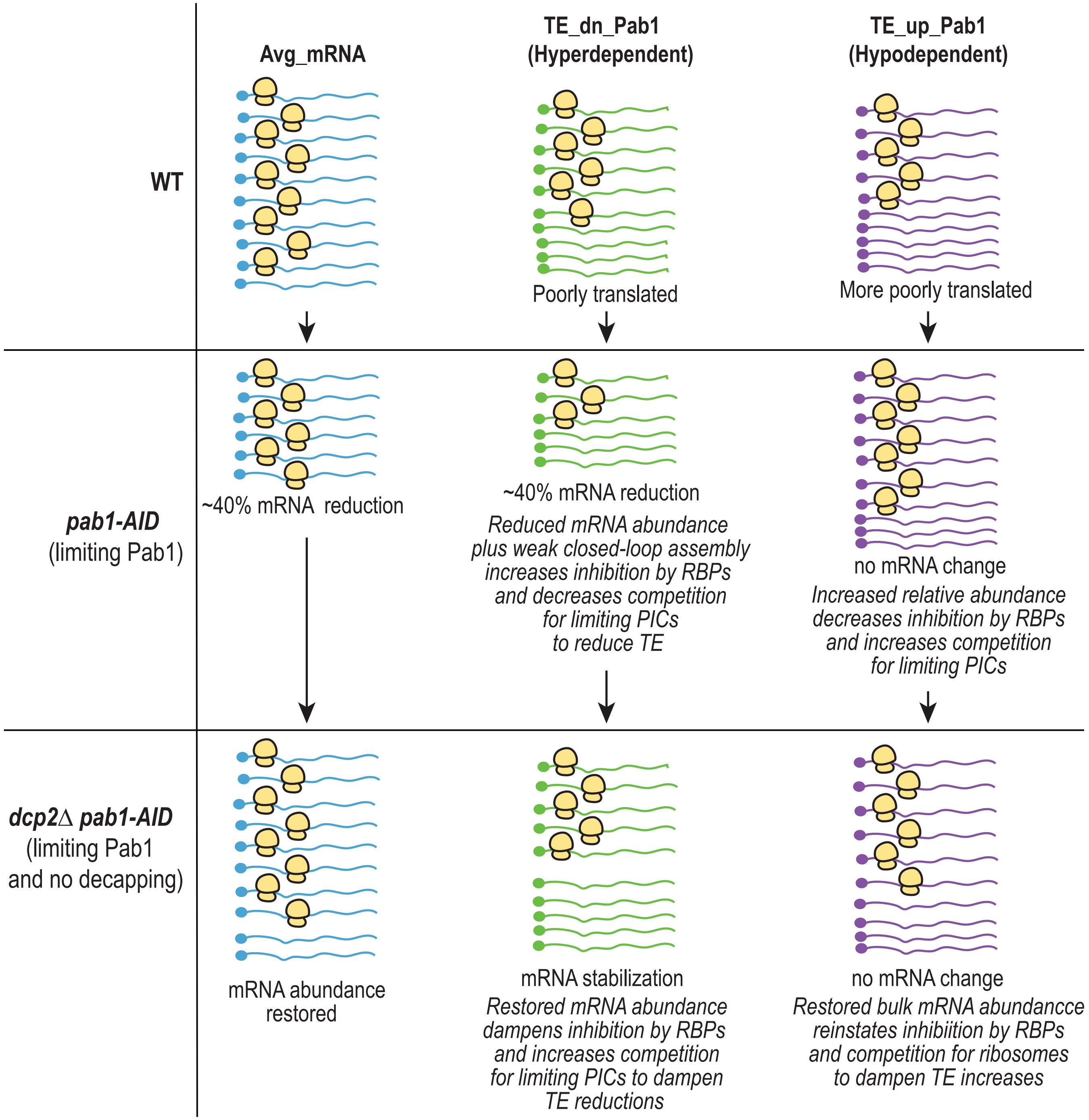

Avg_mRNA
TE_dn_Pab1
(Hyperdependent)
TE_up_Pab1
(Hypodependent)
WT
Poorly translated
More poorly translated
pab1-AID
(limiting Pab1)
~40% mRNA reduction
~40% mRNA reduction
Reduced mRNA abundance
plus weak closed-loop assembly
increases inhibition by RBPs
and decreases competition
for limiting PICs
to reduce TE
no mRNA change
Increased relative abundance
decreases inhibition by RBPs
and increases competition
for limiting PICs
dcp2∆ pab1-AID
(limiting Pab1
and no decapping)
mRNA abundance
restored
mRNA stabilization
Restored mRNA abundance
dampens inhibition by RBPs
and increases competition
for limiting PICs to dampen
TE reductions
no mRNA change
Restored bulk mRNA abundancce
reinstates inhibiition by RBPs
and competition for ribosomes
to dampen TE increases


Fig.S12

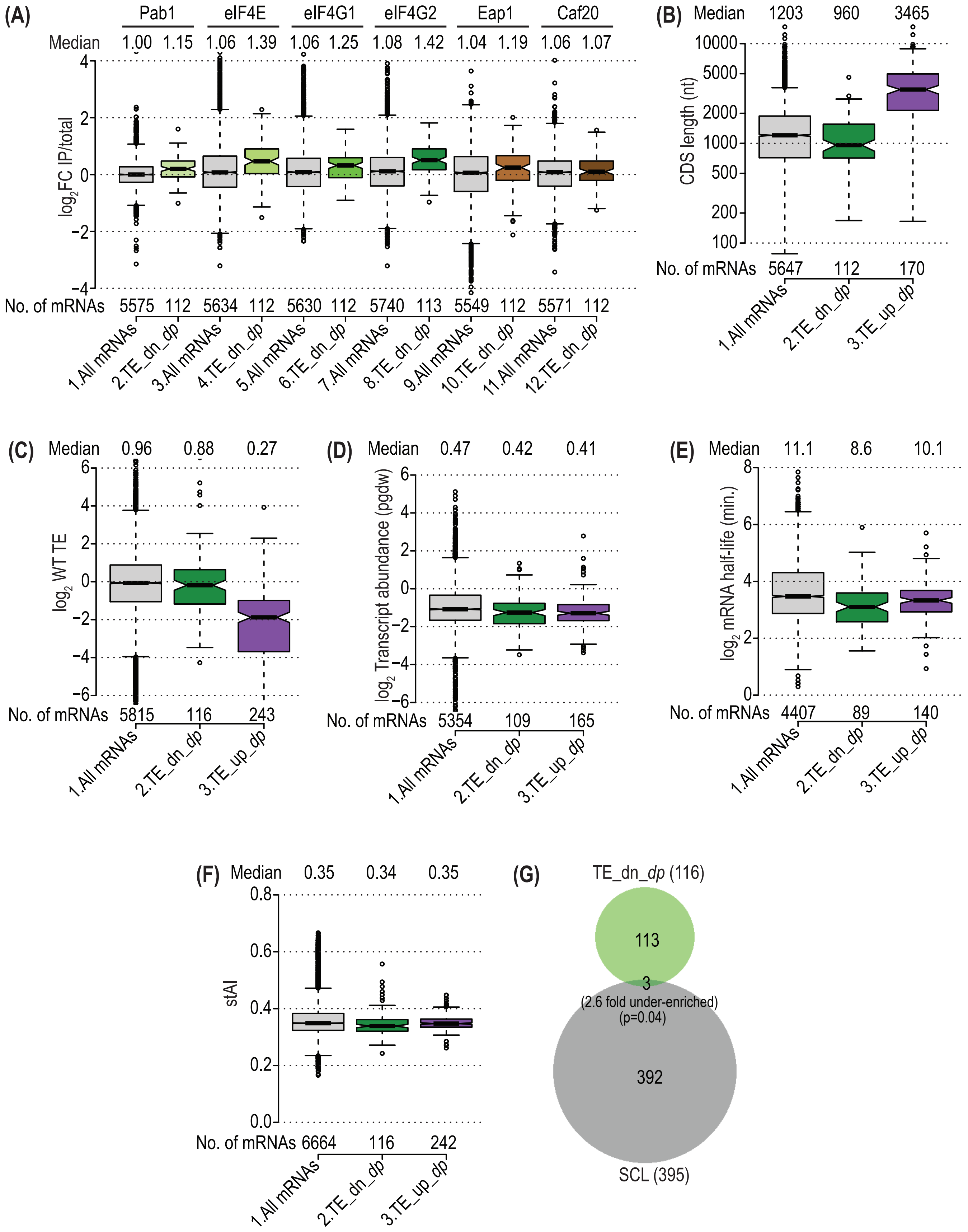

(A)
Pab1
eIF4E
eIF4G1
eIF4G2
Eap1
Caf20
Median
1.00 1.15 1.06 1.39 1.06 1.25 1.08 1.42 1.04 1.19 1.06 1.07
log2FC IP/total
No. of mRNAs
5575 112 5634 112 5630 112 5740 113 5549 112 5571 112
1.All mRNAs
2.TE_dn_dp
3.All mRNAs
4.TE_dn_dp
5.All mRNAs
6.TE_dn_dp
7.All mRNAs
8.TE_dn_dp
9.All mRNAs
10.TE_dn_dp
11.All mRNAs
12.TE_dn_dp
(B)
Median 1203 960 3465
CDS length (nt)
No. of mRNAs 5647 112 170
1.All mRNAs
2.TE_dn_dp
3.TE_up_dp
(C)
Median 0.96 0.88 0.27
log2 WT TE
No. of mRNAs 5815 116 243
1.All mRNAs
2.TE_dn_dp
3.TE_up_dp
(D)
Median 0.47 0.42 0.41
log2 Transcript abundance (pgdw)
No. of mRNAs 5354 109 165
1.All mRNAs
2.TE_dn_dp
3.TE_up_dp
(E)
Median 11.1 8.6 10.1
log2 mRNA half-life (min.)
No. of mRNAs 4407 89 140
1.All mRNAs
2.TE_dn_dp
3.TE_up_dp
(F)
Median 0.35 0.34 0.35
stAI
No. of mRNAs 6664 116 242
1.All mRNAs
2.TE_dn_dp
3.TE_up_dp
(G)
TE_dn_dp (116)
113
3
(2.6 fold under-enriched)
(p=0.04)
392
SCL (395)


**Fig.S13**
